# Heritable single-cell gene expression states shape functional variability in innate immune responses

**DOI:** 10.64898/2026.06.17.732820

**Authors:** Juan Alfonso Redondo, Truong Co Nguyen, Nissrin Alachkar, Josephine Moran, Apurv Srivastav, Marek Kochanczyk, Mark Muldoon, Ian Roberts, Agata Krawczyk-Balska, Abhyduai Singh, Pawel Paszek

## Abstract

Activation of innate immunity at the single-cell level is inherently heterogeneous, yet the mechanisms underlying this variability remain incompletely understood. Here, we integrate transcriptomics, high-content imaging and mathematical modelling to quantify transcriptional heritability within the evolutionarily conserved Toll-like receptor (TLR) system. RNA-seq-based fluctuation tests identified a subset of TLR4-dependent genes, including cytokines and immune effectors, that retain transcriptional heritability for more than 25 cell divisions in clonal macrophage populations. High-content microscopy confirmed gene-specific propagation of heritable states and revealed that environmental context shapes their persistence and expression. CD36, a scavenger receptor involved in bacterial recognition and lipid uptake, exhibited a stable, cell density-reinforced heritable state, whereas the inflammatory programme exemplified by IL1β was transient, with heritability decaying upon clonal expansion. The interplay between heritable transcriptional states and population context generated emergent spatial organisation in high-density populations, with CD36-high cells forming discrete pockets and IL1β-high cells enriched in surrounding regions. Functionally, CD36 expression determined clonal susceptibility to *Listeria monocytogenes* infection, linking transcriptional heritability to heterogeneous infection outcomes. Together, these findings identify transcriptional heritability as a key determinant of innate immune heterogeneity and demonstrate how heritable cellular states interact with population context to generate complex immune behaviours.

**Key point summary:**

- MemorySeq and scRNA-seq identify long-term heritable gene expression states within TLR4-induced macrophage populations.
- High-content imaging across thousands of clonal populations reveals gene-specific dynamics of heritable states at the protein level.
- *Il1β* and *Cd36* define mutually exclusive heritable states that are differently regulated by population context and drive spatial organisation in cellular monolayers.
- Heritable CD36 protein expression shapes heterogeneous outcomes during *Listeria monocytogenes* infection.

## Introduction

In mammals, the innate immune defence against foreign threats is initiated by evolutionarily conserved pattern recognition receptors, including the TLR system [1]. Paradoxically, the activation of TLR signalling is inherently a heterogeneous process with target genes exhibiting substantial transcriptional variability at the single-cell level [2]. Consequently, effector molecules such as Tumour Necrosis Factor α (TNFα), Interleukin 1β (IL1β) or type I interferons (IFN-I) are produced by small (and often non-overlapping) subsets of genetically identical innate immune cells [3–6]. This response heterogeneity is evolutionarily conserved across species [7] suggesting an important and beneficial role during inflammation [8]. However, a fundamental question remains unresolved: does this variability arise *de novo* through stochastic activation, or is it predetermined by the existing cellular states within innate immune populations?

The single-cell TLR-mediated transcriptional response reflects both cell-intrinsic and extrinsic signalling events, such as activation of key transcription factors such as Nuclear Factor κB (NF-κB) [9–13], paracrine signalling, epigenetic regulation, and pathogen-driven cues [3, 5, 14, 15]. Genetically identical cells may behave differently because they occupy distinct cellular states [16–19] or experience stochastic fluctuations in their local environment [20, 21]. Transcriptional variability has been predominantly attributed to transcriptional bursting, which involves stochastic transitions between active and inactive gene states that generate probabilistic transcription events across diverse cell types and tissues [22–28], including the immune signalling context [4, 15, 29–31]. However, recent studies demonstrate that non-genetic heterogeneity, including responses of rare precocious cells, can persist across multiple generations in proliferating populations, revealing long-term transcriptional memory [16, 32, 33]. Within the TLR system, NF-κB signalling exhibits correlated dynamical responses not only over one or a few cell divisions [17, 34], but over days and months in clonally-derived populations [35]. The seemingly stochastic biphasic TLR4-dependent expression of IL1β and IL1α, located in the same gene cluster, is highly correlated and subjected to epigenetic control [4]. Moreover, heritable cellular fates determine the early IFN responses [6], and heritable receptor expression regulates “all-or-nothing” NF-κB activation patterns [34]. Collectively, these findings indicate that transcriptional heritability in innate immunity cannot be explained solely by stochastic gene expression, reflecting instead stable and heritable cellular states.

The existence of the heritable cellular states in innate immunity raises the key mechanistic question of whether such states exert functional consequences during immune responses. During infection, the outcomes of host-pathogen interactions at the single-cell level arise from a complex interplay between heterogeneous host states and bacterial variability [36]. Individual cells infected with common bacterial pathogens, including *Listeria monocytogenes* (*Lm*), *Salmonella enterica*, or *Mycobacterium tuberculosis*, display strikingly divergent fates, ranging from pathogen clearance to bacterial replication or dormancy [37–40]. While microbial-intrinsic gene expression programs contribute to this diversity [14, 41–43], host-intrinsic factors including cell state, tissue microenvironment, and local cell density strongly influence susceptibility to infection [37, 39, 44–46]. For example, single-cell metabolic dynamics of human macrophages predicts *Legionella* intracellular replication [47], whereas susceptibility to *L. monocytogenes* infection has been linked to host cell-to-cell variation in bacterial adhesion [46]. Importantly, heritability within skin cells has been shown to encode resistance to viral infection [48]. Critically, primary innate immune cells, including macrophages, proliferate in the steady-state and in response to inflammation [49–54], allowing propagation of heritable states across cell divisions in their local environment [55]. Together, these observations suggest that pre-existing transcriptional or morphological states can determine infection outcomes, raising the possibility that heritable expression traits may mechanistically underlie such variability.

To distinguish between heritable and non-heritable gene expression traits in innate immune signalling we applied MemorySeq, an RNA-seq-based adaptation of the classical Luria-Delbrück fluctuation test [16], to assay genome-wide TLR4-dependent responses of clonal populations of immortalised murine bone marrow derived macrophages (iBMDMs). Approximately 7% of TLR-dependent genes, including multiple chemokines and cytokines, displayed heritable expression after ∼25 cell divisions, compared to only 2% of TLR-independent genes. Single-cell RNA sequencing (scRNA-seq) confirmed this heterogeneity in mRNA distributions. To validate selected immune-relevant genes at the protein level, we performed high-content immunostaining across thousands of clonal populations spanning ∼10 generations, revealing that transcriptional heritable states persist over multiple cell divisions but are modulated by environmental and population context, resulting in spatial organisations in cellular monolayers. Among these genes, CD36, a scavenger receptor involved in bacterial recognition [56], exhibited an exceptionally high, long-term heritability, prompting further investigation into its role in macrophage susceptibility to *Lm* infection. Together, our findings establish a framework for quantifying and functionally linking transcriptional heritability with cellular immune phenotypes.

## Results

### MemorySeq reveals heritable gene expression patterns in TLR signalling

If a gene exhibits cell-to-cell variability (e.g., high versus low cytokine expression levels following stimulation) and these expression states are maintained across multiple cell divisions, clonal populations derived from individual cells will diverge and show substantial inter-clone variability; some clonal populations will contain mostly high-expressing cells, whereas others will contain mostly low- (or non-) expressing cells. In contrast, if expression fluctuates rapidly and reversibly, clone-to-clone differences will disappear within a few generations, and clonal responses will show similar expression profile to the parental population. Thus, persistent inter-clonal variability, relative to parental populations, provides a quantitative measure of transcriptional heritability (Fig. 1A) [16, 18, 32, 33, 57–60].

**Figure 1:**
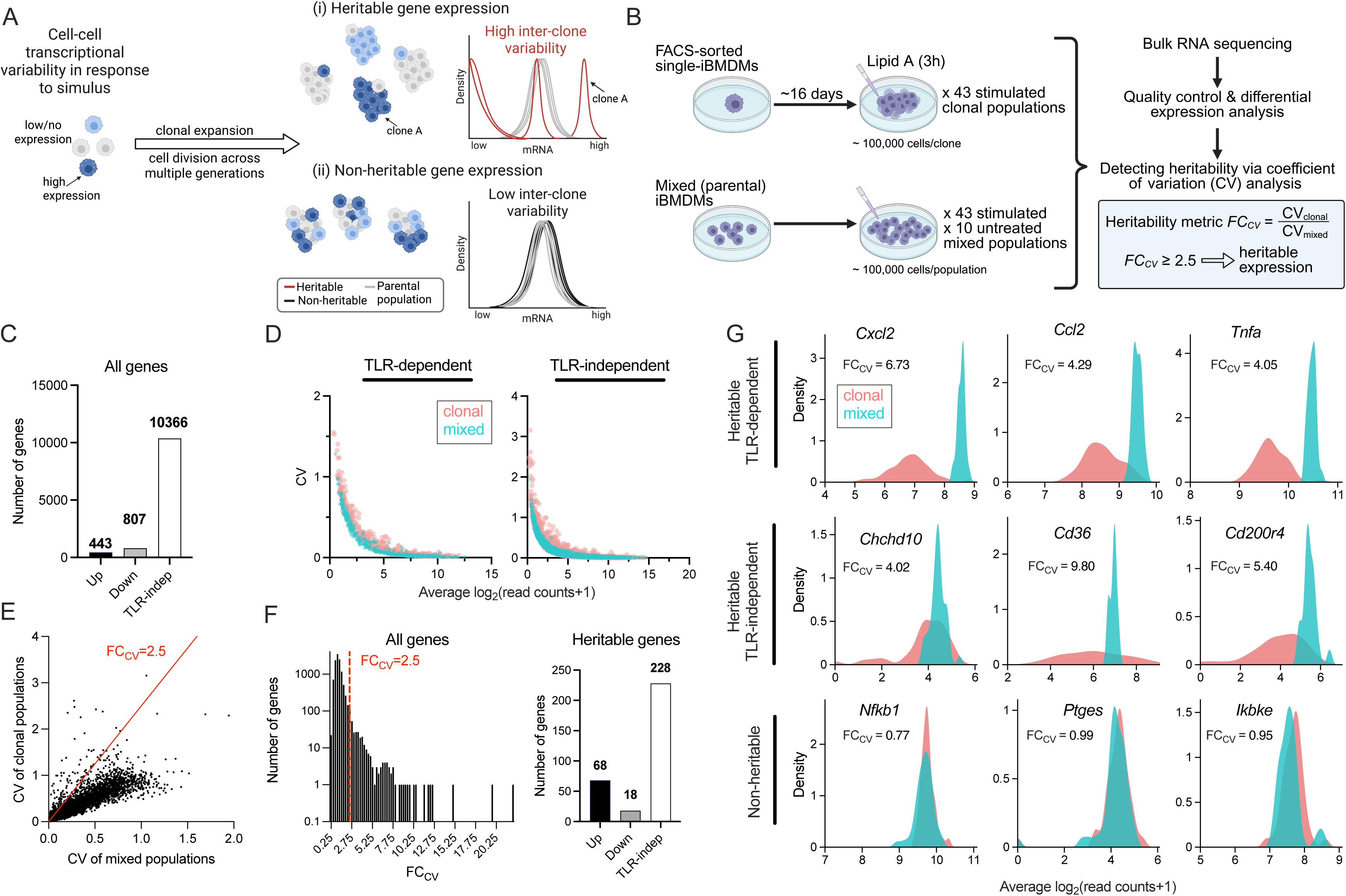
MemorySeq reveals heritable gene expression patterns in the TLR system. **A**. Conceptual framework for transcriptional heritability analysis using single-cell-derived clonal populations. Following stimulation, single cells display heterogeneous gene expression patterns. Upon expansion, transcriptional heritability generates distinct clonal populations with high inter-clone variability, whereas non-heritable expression produces mRNA distributions similar to the parental population. **B**. Schematic representation of the MemorySeq experiment. Individual iBMDM cells were FACS-sorted into separate wells and cultured for 16 days. 43 MemorySeq clones and 43 matching mixed-population controls were stimulated with 500 ng/ml of lipid A for 3 h before RNA sequencing. Ten untreated mixed-population samples were included as controls. The coefficient of variation (CV) across clonal and mixed samples was used to identify heritable genes. **C.** Number of genes identified as TLR-dependent (up- and down-regulated) and TLR-independent using mixed-population data. **D**. Relationship between the CV and mean expression levels for TLR-dependent (left) and TLR-independent (right) genes. Each dot represents the average expression value for a gene across clonal (pink) and mixed (turquoise) samples. **E**. CV of clonal versus mixed samples for the 11,618 MemorySeq genes. The red line corresponds to CV_clonal_ = 2.5 CV_mixed_, FC_CV_ denotes fold change CV of clonal samples vs. CV of mixed samples. **F**. Number of heritable genes. (Left) Histogram of FC_CV_ for the 11,618 MemorySeq genes. The dashed red line marks the heritability fold change threshold FC_CV_ = 2.5, above which 314 genes were identified as heritable. (Right) number of heritable genes classified as TLR-dependent (up- and down-regulated) and TLR-independent. **G**. Examples of heritable (top and middle rows) and non-heritable (bottom) genes. Shown are distributions of expression levels for clonal (pink) and mixed (turquoise) samples from MemorySeq. The FC_CV_ value for each gene is indicated below its name.

To extend this concept to genome-wide innate immune TLR-induced responses, we applied the MemorySeq framework (Fig. 1B), which quantifies heritability based on bulk RNA-seq of clonal populations relative to their parental populations [16]. MemorySeq clones were generated using isogenic iBMDMs, a well-established model of macrophage TLR signalling [14, 61]. Unlike primary macrophages, which have limited proliferative capacity *in vitro*, iBMDMs provide an experimentally tractable system for studying propagation of heritable cellular states across multiple cell divisions. Each clone, derived from a single cell by fluorescence-activated cell sorting (FACS), was expanded to approximately 100,000 cells over 15–16 days, corresponding to ∼25 cell divisions. As a reference, we cultured and sequenced matched parental iBMDM populations (hereafter referred to as mixed populations), which were seeded one day prior to the experiment at densities adjusted to yield the same final cell number as the clonal populations, thereby controlling for technical and sampling variability. Both clonal and mixed populations were stimulated for 3 h with 500 ng/ml lipid A, the bioactive component of bacterial lipopolysaccharide and a TLR4 ligand [62], prior to bulk RNA sequencing. From an initial set of 96 samples comprising 43 lipid A-stimulated clonal populations, 43 stimulated mixed populations, and 10 untreated mixed populations, we retained 41 stimulated MemorySeq clones, along with 43 stimulated and 10 untreated mixed populations after library preparation, sequencing, and quality control (Fig. 1B). Principal component analyses (PCA) demonstrated a clear separation between conditions, except for one clonal sample which was subsequently excluded (Fig. S1). Sequencing depth exceeded 500,000 reads per sample (mean of 2.9 million reads). Differential expression analysis of the mixed cell populations identified 1,251 TLR4-dependent genes (referred herein as TRL-dependent) significantly regulated by lipid A (443 upregulated and 807 downregulated), whereas 10,366 genes were unaffected and classified as TLR-independent (Fig. 1C).

To quantify transcriptional variability across clonal and mixed populations, we calculated the coefficient of variation (CV) for each gene, defined as the standard deviation normalized by the mean of read counts. Gene expression variability decreased with increasing mean expression whereas a subset of genes exhibited substantially higher inter-clone variability than that observed in the corresponding mixed populations (Fig. 1D). We quantified transcriptional heritability using the ratio of clonal-to-mixed CV (F_CV_ = CV_clonal_/CV_mixed_), defining heritable genes as those with *FC_cv_* ≥ 2.5 (Fig. 1E). This analysis identified 314 genes with heritable expression patterns (Fig. 1F). Among these, 86 genes were TLR-dependent (including 68 upregulated; corresponding to 7% of all TRL-dependent genes and 15% of the upregulated subset), whereas 228 genes (2%) were classified as TLR-independent (Fig. 1G for representative example genes, see Table S1 for full gene lists and).

Genes identified as heritable in our assay did not overlap with previously reported resistance-associated loci from non-phagocytic cell lines [16, 18], but instead reflected macrophage-specific innate immune functions. Heritable TLR-dependent genes included key inflammatory chemokines (*Cxcl2*, *Ccl2*, *Ccl3*, *Ccl17*) and cytokines (*Tnfa*, *Il6*, *Il1b*, *Csf1*, *Csf2*) (Fig. 1G), together with their cognate receptors (*Ccrl2* and *Tnfrsf1b*, encoding TNFR2). The inflammasome component *Nlrp3* [63] and *Ptgs2* (encoding COX-2) [64] were also heritable. Within the core TLR-NF-κB signalling network [65], none of the principal transcription factors (*Rela*, *Relb*, *Nfkb1*, *Nfkb2*), their inhibitors (*Nfkbia*, *Nfkbib*, *Nfkbieb*) or the kinases *Ikbkb* and *Ikbkg*, meet the heritability threshold. In contrast, *Tnfaip3*, encoding the ubiquitin-editing enzyme A20, a pivotal negative regulator of NF-κB activity [66, 67], displayed strong heritability. No heritable expression was detected for components of the Janus kinase-Signal Transducer and Activator of Transcription (JAK-STAT) pathway [68] or for type I/II IFN receptors. However, their inhibitor, Suppressor of Cytokine Signalling 3 (*Socs3),* but not *Socs1,* was heritable. Among interferon-regulated transcription factors, only *Irf1* exhibited heritability, consistent with its broad regulatory role in macrophage activation [69]. Finally, several TLR-independent heritable genes were associated with regulation of phagocytosis (*Cd36*, *Msr1*, *Abca1*, *Marcks*, *Plekho*, *Slc11a1*, *Rab20*, *Clcn7*, *Lyz1/2*, *Ctsc)* [70], extracellular matrix organisation (*Itga5-8*) [71], the complement system (*C1qa*, *C1qb*, *C1qc*, *C1ra*) [72], and macrophage differentiation and signalling (see Table S2 for gene ontology analysis).

Overall, these analyses demonstrate the long-term persistence of heritable gene expression states within the TLR signalling network.

### Single-cell RNA-seq demonstrates distinct inter-clone expression distributions of heritable genes

We identified over 300 genes that were classified as heritable using population-based MemorySeq approach. To validate these findings at single-cell resolution, we performed scRNA-seq on individual clonal and mixed populations and quantified transcriptional heritability using the same CV-based framework, enabling direct comparison between these two approaches.

Using a multiplexed 10x Genomics protocol (see Methods), we simultaneously profiled eight newly derived clonal and four mixed populations, all stimulated with lipid A for 3 h (Fig. 2A). Following quality control, we obtained 9,166 single-cell transcriptomes, averaging 764 ± 166 cells per clonal population. Cells contained a median of ∼56,000 unique molecular identifiers (UMIs) and ∼6,900 detected genes per cell, with ∼20,000 genes identified across all populations. t-distributed stochastic neighbour embedding (t-SNE) analysis of single-cell gene expression levels revealed that individual clonal and mixed cells formed overlapping clusters (Fig. S2). Transcriptomes were broadly stratified by cell-cycle phase, consistent with expected phase distributions [73] (Fig. S3). Importantly, cell-cycle composition was uniform across clones and mixed controls, indicating that observed expression heterogeneity was not driven by differences in proliferative state. From the scRNA-seq data, we calculated mean expression across all populations and identified 5,791 robustly expressed genes, of which 5,701 overlapped with the MemorySeq dataset. For these genes, heritability metric (*FC_cv_*) was calculated (Table S3), revealing numerous genes with substantial inter-clone variability (Table S4). Unlike MemorySeq, which measures bulk averages, scRNA-seq captures mRNA distributions at single-cell resolution. However, the smaller number of samples (8 clonal and 4 mixed populations, compared to >40 MemorySeq samples) reduced statistical power. To account for sampling effects, we evaluated multiple *FC_cv_* thresholds and assessed reproducibility between datasets. A threshold of *FC_cv_* = 3 minimized the rate of false positives (genes classified as heritable in scRNA-seq but not in MemorySeq) to ∼20%, while allowing a moderate increase in false negative rates (Fig. 2B).

**Figure 2:**
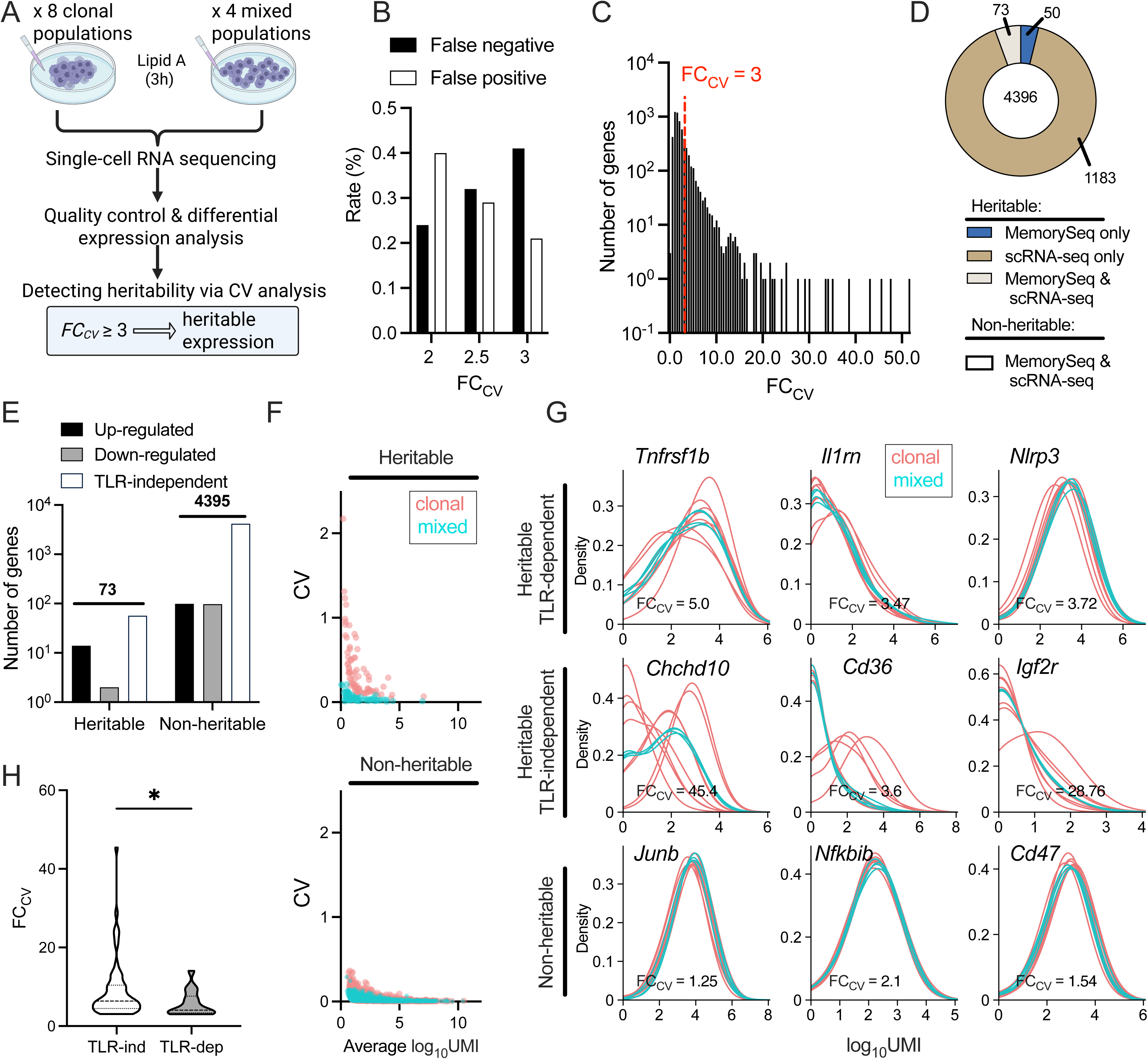
scRNA-seq analysis of gene expression heritability. **A.** Schematic representation of the scRNA-seq experiment, involving 8 clonal and 4 mixed populations stimulated with 500 ng/ml of lipid A for 3 h. Analysis of CV across clonal samples versus mixed samples was used to identify heritable immune genes using a 3-fold change threshold. Genes over this threshold were subsequently compared to MemorySeq data. **B**. Sensitivity analysis of the heritability metric FC_CV_. Shown are false-positive and false-negative rates (%) for the identification of heritable genes in scRNA-seq data, relative to MemorySeq inference, across different FC_CV_ thresholds (2-, 2.5- and 3-fold). **C.** Histogram of FC_CV_ values for clonal versus mixed samples across 5,701 genes in the scRNA-seq dataset. The dashed red line indicates the FC_CV_ = 3 threshold, above which 1,256 genes were classified as heritable. **D.** Overlap between MemorySeq and scRNA-seq datasets. Pie chart showing genes identified as heritable in both experiments (grey), in MemorySeq only (blue), in scRNA-seq only (brown), or non-heritable in both (white). **E.** Number of TLR-dependent (up- and down-regulated) and TLR-independent genes identified as heritable or non-heritable in both MemorySeq and scRNA-seq experiments. **F.** Relationship between CV and mean expression levels across samples for the 73 genes classified as heritable in both MemorySeq and scRNA-seq datasets (top) and for the 4,396 non-heritable genes (bottom). Each dot represents the average expression value for a gene across its clonal (pink) and mixed (turquoise) samples. **G**. Examples of heritable (top and middle rows) and non-heritable (bottom) genes from E. Shown are scRNA-seq gene expression distributions for clonal (pink) and mixed (turquoise) populations. The FC_CV_ value for each gene is indicated below its name. **H**. Comparison of FC_CV_ between the 73 TLR-dependent and -independent heritable genes. Statistical significance was assessed using Mann-Whitney test (\**p*-value < 0.05).

We identified 1,256 genes that met the heritability criterion in scRNA-seq (Fig. 2C). The concordance between datasets reached 78% across 5,701 shared genes (Fig. 2D). Of the 314 genes classified as heritable by MemorySeq, 123 were robustly detected in the scRNA-seq dataset (39%). Among these, 73 genes (59%) were independently classified as heritable in scRNA-seq, comprising 16 TLR-dependent genes (14 upregulated by lipid A) and 57 TLR-independent genes (Fig. 2E). This overlap between datasets represents ∼2.7-fold enrichment over chance (Fisher exact, p-value = 1.6×10^-19^). As expected, inter-clone variability, CV, was markedly higher for heritable genes compared with both mixed controls and non-heritable genes (Fig. 2F). Single-cell mRNA distributions further supported these findings: mixed populations exhibited highly overlapping profiles across genes, resembling non-heritable clonal distributions, whereas heritable genes displayed distinct and variable distribution shapes, including shifts in modality (Fig. 2G). Among the heritable TLR-dependent genes identified in both datasets were *Tnfrsf1b*, *Il1rn*, *Csf1*, and *Nlrp3*, all associated with acute inflammatory responses. Among the TLR-independent heritable genes, *Chchd10*, *Cd36* and *Igf2r*, three genes that cooperatively regulate lipid metabolism, mitochondrial homeostasis, and lysosomal trafficking, showed the highest clone-clone differences (Fig. 2G; Table S5). Notably, the heritability metric differed significantly between TLR-dependent and TLR-independent genes (Fig. 2H), indicating that TLR-dependent genes generally exhibit weaker transcriptional heritability, reflected by lower inter-clone variability and more homogeneous single-cell mRNA distributions (Fig. 2G). This suggests that core TLR-responsive genes are subject to tighter regulatory control, limiting long-term transcriptional divergence across clones.

Mechanistically, heritable genes may exhibit coordinated fluctuations reflecting shared transcriptional regulation, such as common upstream transcription factors or engagement of broader regulatory networks [16]. Accordingly, if a clonal population or a single cell displays elevated expression of one heritable gene, transcripts that co-fluctuate with it are expected to show correlated abundance patterns. To quantify these relationships, we calculated pairwise Pearson’s correlation coefficients among the 73 heritable genes across both the MemorySeq and scRNA-seq datasets. This analysis revealed distinct modules of co-regulated genes exhibiting strong positive correlations in MemorySeq and, albeit with greater noise, similar patterns in the scRNA-seq data (Fig. S4A). All TLR-dependent genes (except *Napepld*, which encodes a phospholipid-modifying enzyme) displayed uniformly high cross-correlations, consistent with coordinated stimulus-induced activation (Fig. S4B). We also identified clusters of co-fluctuating TLR-independent genes in both datasets (Fig. S4C), including components of the complement system (*C1qc*, *C1qb*) and several transmembrane proteins (*Cd36*, *Cd33*, *Cd200r4*), among others. Together, these results demonstrate strong concordance between datasets and suggest that transcriptional heritability operates at the level of regulatory modules rather than isolated genes.

Overall, these scRNA-seq analyses validate the heritable gene expression patterns identified by MemorySeq and reveal distinct heritability architectures between TLR-dependent and - independent gene classes.

### High-content microscopy reveals heritability at the protein level

To assess the temporal stability of transcriptional heritability and validate RNA-seq findings beyond a single time point, we performed high-content microscopy to monitor protein expression in clonal iBMDM populations (Fig. 3A). We first focused on the TLR-dependent cytokines TNFα and IL1β, key mediators of inflammation and infection [4], both identified as heritable at the mRNA level (FC_cv_ = 4.05 and 2.9, respectively, in MemorySeq). Single iBMDM cells were seeded at low density (20 cells per well, 96-well plates) to ensure spatial segregation of growing clones and were analysed on days 3 and 6. In parallel, mixed (parental) populations were plated at four densities (500, 2,500, 5,000 and 10,000 cells per well; 1.6×10³–3.1×10⁴ cells/cm²) and incubated for 24 h. All populations were seeded on the same 96-well plate on staggered days to enable simultaneous analysis across conditions. For each condition, half of the samples were left untreated, and half were stimulated with 500 ng/ml lipid A for 3 h (see Fig. S5A for the plate layout). Images were quantified using ShuttleTracker software to distinguish nuclear and perinuclear (cytoplasmic) signals and extract morphological features [74] (see Table S6 for segmentation parameters). Across six independent experiments, we analysed 1,172 lipid A-treated and 839 untreated clones, encompassing approximately 260,000 cells. Clone sizes spanned three orders of magnitude (4–1,500 cells; displayed in log_2_ scale in Fig. 3B). The largest clones (>1000 cells at day 6) were consistent with the measured doubling time of 14.4 ± 1.5 h (Table S7), confirming their origin from single founder cells. Smaller clones in the dataset likely resulted from spatial dispersion of proliferating cells. All clonal populations were manually delineated following automated segmentation.

**Figure 3.**
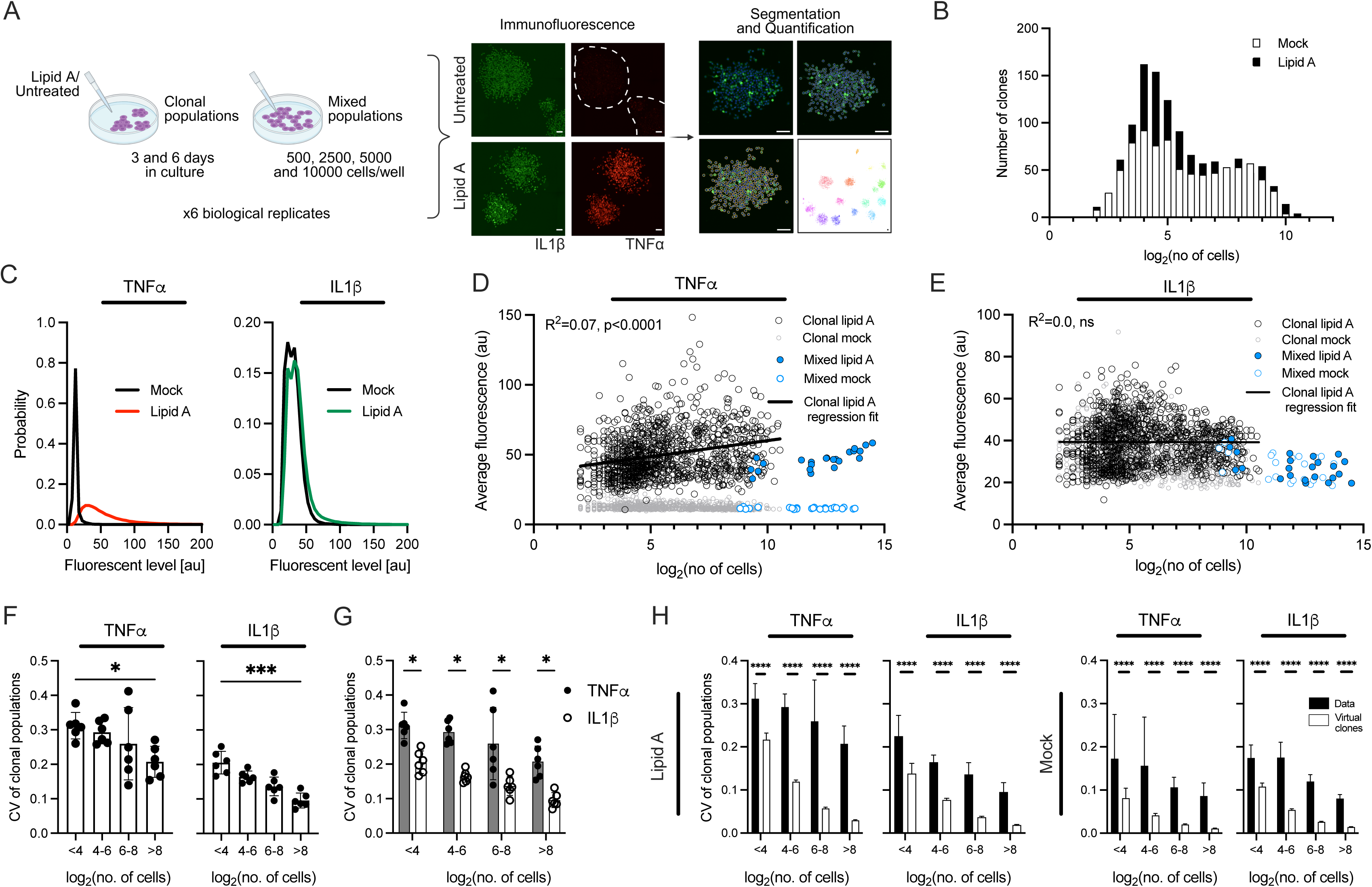
High content microscopy of IL1β and TNFα protein expression patterns in clonal populations. **A.** Schematic representation of high-content imaging experiments and image segmentation workflow. iBMDM cells were seeded at 20 cells per well in 96-well plates and assayed on days 3 and 6, either untreated or after stimulation with 500 ng/ml lipid A for 3 h. Mixed parental populations were seeded at 500, 2,500, 5,000, and 10,000 cells per well and assayed 24 h later under identical treatment conditions. Representative images of clonal populations co-stained for IL1β and TNFα show single-cell segmentation and quantification of protein expression using ShuttleTracker software. Dashed outlines highlight the absence of TNFα-expression in untreated samples. Scale bar, 50 μm. **B.** Histogram of clonal population size distributions. Shown are log₂-transformed clone sizes for untreated (Mock) and lipid A-stimulated conditions. Data pooled across six biological replicates (plates, as in A). **C.** Probability distributions of perinuclear TNFα and IL1β fluorescence signal intensities (arbitrary units) in the clonal populations shown in B. **D.** Clonal populations exhibit heterogeneity in TNFα expression. Shown is the relationship between average TNFα signal intensity per clone and clone size (log_2_-transformed number of cells). Black circles indicate individual lipid A-stimulated clonal populations; grey circles represent untreated (Mock) clonal populations. Filled blue circles show average TNFα expression in lipid A-stimulated mixed populations across a range of seeding densities (expressed as average signal per well versus the total cell number per well across six plates), while open blue circles correspond to untreated mixed populations for a range of seeding densities. The black line indicates the linear regression fit for lipid A-stimulated clonal populations; the coefficient of determination (R^2^) and the *p*-value for the non-zero slope of the regression line are shown. **E.** Clonal populations exhibit heterogeneity in IL1β expression. Relationship between average IL1β signal intensity per clone and clone size (log₂-transformed). Figure formatting as in D; ns, non-significant. **F.** Inter-clonal variability as function of clone size for data from D and E. Shown are CVs for lipid A-induced average TNFα (left) and IL1β (right) fluorescent intensities, binned into four population size categories: log₂N ≤ 4, 4 < log₂N ≤ 6, 6 < log₂N ≤8, log₂N >8, where N is the number of cells per clone. Bar chart represents means ± SDs across six biological replicates (plates); circles denote individual plate data. Statistical significance was assessed using the Kruskal-Wallis test with multiple comparisons (\**p* < 0.05; \*\*\**p* < 0.001). **G.** TNFα exhibits a higher level of heritability than IL1β. Comparison of CVs for TNFα and IL1β (as in F). Bar chart represents means ± SDs of six biological replicates (plates). Circles represent individual replicate data. Statistical significance was assessed using the multiple paired t-test with Holm-Sidak correction (\**p* < 0.05). **H.** Inter-clone heritability in clonal TNFα and IL1β expression cannot be explained by chance. Comparison of CVs for average TNFα and IL1β signal intensities between experimental data and virtual clones. Data were binned into four population size categories (as in F). For virtual clones, the number and size of individual clones were matched to the experimental data, but single-cell expression values were randomly sampled from control mixed populations (seeded at 10,000 cells per well). Bar charts for virtual clones show means ± SDs derived from bootstrap resampling across six individual plates. Statistical significance was assessed by comparing experimental mean CVs to corresponding bootstrap distributions per plate; combined *p*-values were calculated using Fisher’s method (\*\*\**p* < 0.0001).

We next quantified the heritability of TNFα and IL1β protein expression across clonal populations. TNFα levels exhibited robust lipid A-induced activation in nearly all cells, whereas untreated populations showed minimal signal (Fig. 3A and C; see Fig. S5B-C for biological replicates data). In contrast, only a small subset of clonal and mixed-population cells produced robust IL1β following stimulation. These distinct activation patterns are consistent with our previous single-cell mRNA analyses, in which *Tnfa* transcripts showed relatively uniform induction across cells, whereas *Il1b* expression was highly heterogeneous, ranging from tens to thousands of mRNA copies in genetically identical cells [4].

Upon visual inspection, individual clones displayed distinct TNFα expression levels and variable fractions of IL1β-producing cells following stimulation (Fig. 3A). Quantitatively, clones of comparable size (*N*) differed substantially in their average protein expression levels (Fig. 3D-E). When analysed as a function of clone size, approximating the number of cell divisions (log₂*N*), the average TNFα intensity showed a modest but statistically significant positive correlation with clone size (R^2^ = 0.07; Fig. 3D). In contrast, average IL1β levels varied widely among clones but showed no dependence on clone size (Fig. 3E). In both cases, mixed-population controls exhibited markedly lower variability than clonal populations, consistent with inter-clone heterogeneity rather than measurement noise. We next quantified clone-to-clone variability in TNFα and IL1β protein expression using the coefficient of variation (CV; Fig. 3F-H). Data were grouped into four approximately equal-sized bins based on log₂N (≤ 4, 4–6, 6–8, and > 8), spanning from small (<16 cells) to large clones (>256 cells). Overall, inter-clone variability was highest among small clones and decreased progressively with clone size (Fig. 3F-G). Although IL1β expression is more heterogeneous at the single-cell level, TNFα exhibited significantly stronger inter-clone divergence across all size categories (e.g., CV = 0.31 ± 0.04 vs. 0.21 ± 0.03 for log₂*N* < 4; 0.21 ± 0.05 vs. 0.10 ± 0.02 for log₂*N* > 8; Fig. 3G). Consistent with the MemorySeq data, this suggests more stable propagation of its expression state across divisions. To determine whether the observed variability could arise by chance, we computationally generated “virtual clones” matching the experimental clone-size distribution but randomly sampled with single-cell expression values from mixed populations (Fig. 3H). Virtual clones exhibited markedly lower clone-to-clone variability for both TNFα and IL1β, even among small populations (log₂*N* ≤ 4), where stochastic effects are expected to be strongest. These results indicate that the experimentally observed inter-clone variability in protein expression substantially exceeds that predicted from random fluctuations alone.

Building on these observations, we expanded our high-content immunostaining analyses to additional targets, to further validate the RNA-seq findings (Fig. 4A). We selected the heritable TLR4-dependent *Tnfrsf1b* (encoding TNFR2) alongside the non-heritable NF-κB family members RelA and RelB [75]. To assess heritability beyond the TLR axis, we also included two TLR-independent genes: *Cd36*, encoding platelet glycoprotein 4 involved in bacterial recognition and lipid metabolism [56], classified as heritable in the MemorySeq dataset; and *Adgre1* (encoding F4/80) [76], a canonical macrophage marker, classified as non-heritable (see Fig. S6 and S7 for plate layouts and distributions on clonal versus mixed populations). Comparisons of heritability scores between MemorySeq and scRNA-seq datasets confirmed overall concordance for these targets (Fig. S8A). This panel enabled direct comparison of heritable and non-heritable genes within and outside the core TLR signalling pathway.

**Figure 4.**
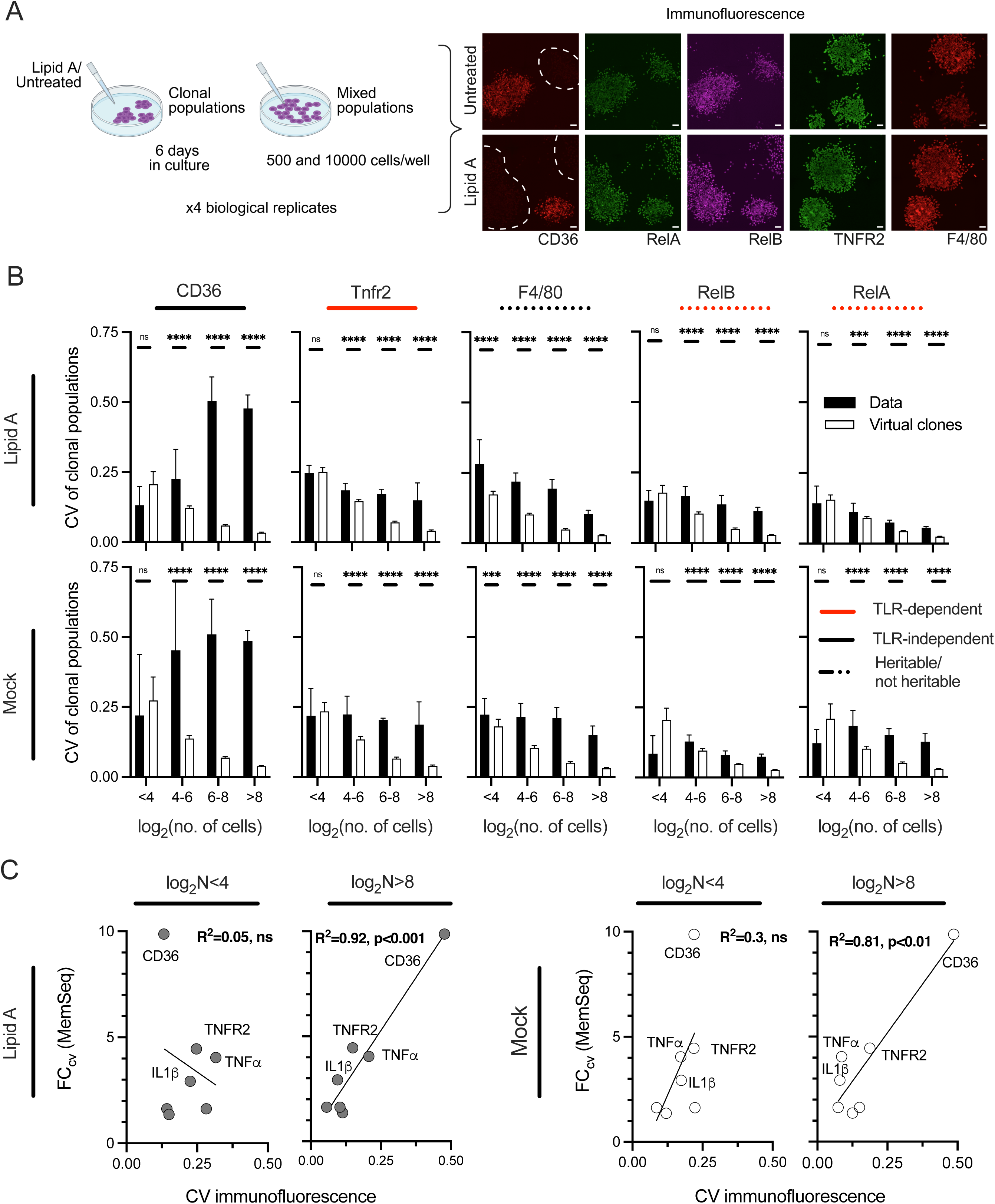
Heritability analysis across diverse TLR-dependent and -independent targets. **A.** Schematic representation of high-content imaging experiments for RelA, RelB, CD36, TNFR2 and F4/80. iBMDM cells were seeded at 20 cells per well in a 96-well plate and assayed on day 6, either untreated or after stimulation with 500 ng/ml lipid A for 3 h. Mixed populations were seeded at 500 and 10,000 cells per well and assayed 24 hours after seeding, under the same treatment conditions. Representative images of clonal populations co-stained for RelA, RelB and CD36, and separately co-stained for TNFR2 and F4/80. Dashed outlines highlight CD36-negative populations. Scale bar, 50 µm. **B.** Patterns of observed inter-clonal variability differ from random chance. Shown is the comparison of CVs for average signal intensities between experimental data and simulated virtual clones for the targets from A. Data were binned into four population size categories (as in Fig. 3F-H). Experimental data represented as means ± SDs of four biological replicates (plates). For virtual clones the number and size of individual clones were matched to the experimental data, but single-cell expression values were randomly sampled from the control mixed populations (seeded at 10,000 cells per well). Bar charts for virtual clones show means ± SDs derived from bootstrap resampling across four individual plates. Statistical significance was assessed by comparing experimental mean CVs to corresponding bootstrap distributions for individual plates; combined *p*-values were calculated using Fisher’s method (ns - non-significant, \*\*\**p* < 0.001, \*\*\*\**p* < 0.0001). **C.** Inter-clonal variability levels are consistent across imaging and RNA-seq datasets. Shown is the relationship between CVs obtained from immunofluorescence data (as in B and Fig. 3H) and the heritability metric (FC_CV_) derived from the MemorySeq analysis for TNFα, IL1β, RelA, RelB, CD36, TNFR2 and F4/80. Each circle represents an individual gene, with the determination coefficient (R^2^) and *p*-value for non-zero slope of the regression line indicated. Data are shown for two clone size categories, log₂N ≤ 4 and log₂N > 8, stratified by the number of cells per clone N, for untreated (Mock, right) and lipid A-stimulated (left) conditions. Heritable gene names are labelled.

For clonal analyses, single iBMDM cells were seeded at low density (20 cells per well, 96-well plates) and analysed on day 6, either untreated or following 3 h stimulation with 500 ng/ml lipid A. Mixed-population controls (seeded at 500 and 10,000 cells per well, 1.6×10³ and 3.1×10⁴ cells/cm²) were plated in parallel and incubated for 24 h (Figs. S6AB and S7A). Approximately 1,100 clones (∼220,000 cells) were co-stained for RelA, RelB and CD36, and ∼950 clones (∼130,000 cells) were co-stained for TNFR2 and F4/80. Clone size spanned three orders of magnitude under both treatment conditions (Fig. S8C). Consistent with established stimuli-induced single-cell NF-κB dynamics [77], lipid A treatment induced enhanced nuclear localisation of RelA and RelB in a subset of cells (Fig. S6D-E). CD36 expression was heterogeneous, with only a fraction of cells within both clonal and mixed populations exhibiting strong staining (Fig. 4A, Fig. S8B). Although previous studies reported TLR-mediated transcriptional repression of CD36 [78], we observed only a modest downregulation in lipid A-treated samples compared with untreated controls (Fig. S6B-C), likely reflecting its relatively long (∼16 h) protein half-life [79] compared with the 3 h stimulation period used in our assays. Finally, TNFR2 and F4/80 were predominantly cytoplasmic and showed minimal change upon stimulation, despite *Tnfrsf1b* mRNA upregulation in the MemorySeq dataset, likely reflecting high basal protein expression (Fig. S7B-C).

Among the analysed targets, CD36 exhibited the strongest heritability by far (FC_CV_ = 9.87 and 3.6 in the MemorySeq and scRNA-seq analyses, respectively; Fig. 1G, Fig 2G, and Fig. S8A). Consistent with this, visual inspection revealed marked clonal divergence: some clones comprised predominantly CD36-positive cells, whereas others lacked detectable expression entirely (Fig. 4A). Following image segmentation and quantification, average protein expression per clone was calculated as a function of clone size (Fig. S8D), and inter-clone variability was quantified using CV across four clonal-population-size categories and compared with virtual clones (Fig. 4B). CD36 displayed the highest CV values among all targets, and variability increased with clone size under both untreated and lipid A-stimulated conditions. This variability substantially exceeded that observed in virtual clones, except in the smallest populations (log₂N ≤ 4), where sampling noise is expected to be higher due to a smaller sample size. TNFR2 showed lower overall variability (CV = 0.48 ± 0.04 for CD36 vs. 0.15 ± 0.05 for TNFR2 under lipid A stimulation, log₂N > 8) but variability remained stable across clone sizes (log₂N > 4) and significantly exceeded that of virtual clones. Interestingly, non-heritable genes such as RelA, RelB, and F4/80 also displayed modest size-dependent heritability signatures, with larger clones showing greater variability than expected by chance. This suggests that even genes classified as non-heritable at the mRNA level, may exhibit heritability for at least 10 divisions.

To systematically compare inter-clone variability across all seven targets, we correlated MemorySeq heritability scores with CV values obtained from immunostaining analyses (Fig. 4C). For large clonal populations (log₂N > 8), the two measures were strongly correlated (R^2^ = 0.92 for lipid A-treated and 0.81 for untreated samples), demonstrating excellent concordance between RNA-seq and protein-based datasets. In contrast, correlations were weaker for small clones (log₂N < 4), as a limited number of cell divisions preserves apparent inter-clone variability across most targets, including genes classified as non-heritable. Consequently, discrimination between heritable and non-heritable genes is reduced in smaller populations compared to large populations that have undergone extensive expansion. To further validate this point, we leveraged collected imaging datasets (500 cells per well) and computationally identified daughter-cells pairs as cells positioned in close proximity to each other but spatially isolated from neighbouring cells (Fig. S9A, see Methods). Across all target genes, we observed high Pearson’s correlations within daughter-cell pairs, ranging from approximately 0.55 for inducible TNFα and IL1β to 0.70–0.85 for the remaining genes (Fig. S9B). These findings indicate that recently dividing cells retain highly similar protein expression states.

Together, these analyses reveal that transcriptional heterogeneity in the TLR signalling system exhibits varying degrees of protein level-heritability, with both strongly and weakly heritable targets retaining measurable memory across multiple generations.

### CD36 and IL1β exhibit distinct heritability modes shaped by population context

Our analyses support a model in which recently divided daughter cells are highly correlated, generating inter-clone variability that progressively diminishes over successive divisions in a gene-specific manner. However, the immune activation is strongly influenced by the local tissue environment and population context [9, 80–82]. To examine how transcriptional heritability within the innate immune system interacts with population context, we compared single-cell protein responses for IL1β, a low-frequency, stimulus-induced TLR target, and CD36, which exhibits a stable, heritable expression that persists across many generations.

We quantified IL1β and CD36 single-cell protein expression by classifying cells as responsive (“positive”) or non-responsive (“negative”) and characterising each clonal population by the fraction of responding/positive cells (Fig. 5A; see Methods). Lipid A-induced IL1β protein expression was rare at the single-cell level, particularly in mixed populations (up to 3.2 ± 0.8% at the lowest seeding density, Fig. 5B; Fig. S10A-B for mixed populations) [81, 82], but clonal populations exhibited pronounced inter-clone heterogeneity, with high fractions of IL1β-positive cells in some small clones (up to 75%, Fig. 5B, Fig. S10A). As clone size increased, the fraction of responding cells progressively converged toward levels observed in mixed populations, suggesting gradual dilution of heritable traits and an increasing influence of population-level cues. Nevertheless, even the largest clones retained elevated IL1β responsiveness (up to 25% responding cells per clone) relative to mixed population controls (Fig. 5B), indicating a persistent heritable component. This was supported by high coordination between daughter cells (odds ratio (OR) = 9.6 ± 5.5, Fig. 5C), indicating that concordant pairs (ON-ON or OFF-OFF) were nearly 10-times more likely than discordant pairs (ON-OFF), and a reduction in inter-clone variability with increasing clone size (Fig. 5D).

**Figure 5.**
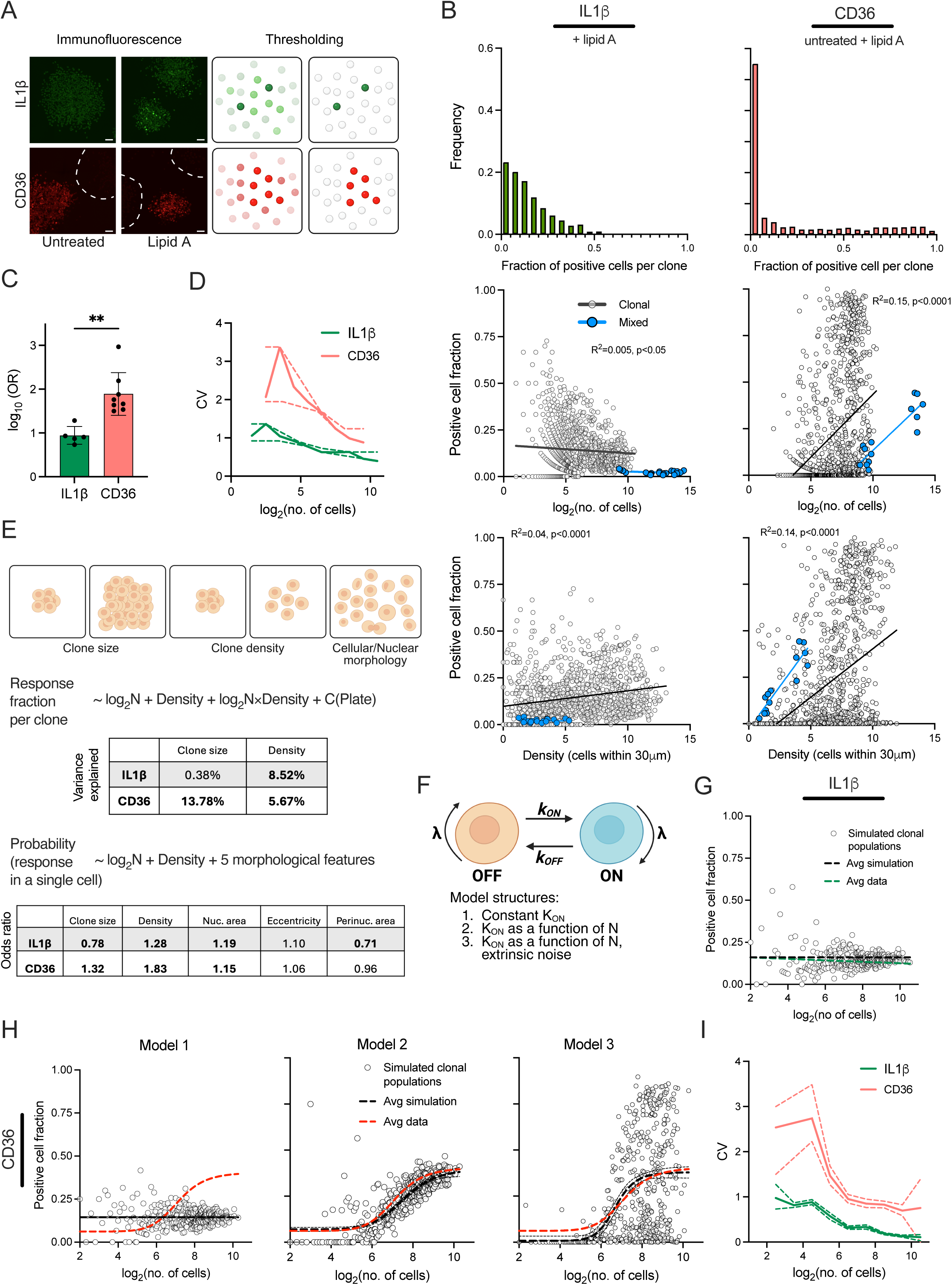
Population context contributes to transcriptional heritability. **A.** Representative examples of clonal populations immunostained for IL1β and CD36. Dashed outlines indicate CD36-negative clones. Scale bar, 50 µm. Individual cells were thresholded based on fluorescence intensity and classified as either responding/positive or non-responding/negative, yielding a binary representation of single-cell protein expression. **B.** Fraction of IL1β and CD36 responding cells exhibit different regulatory modes. Top: Histogram of the fraction of IL1β-positive (lipid A-stimulated) and CD36-positive cells (merged across untreated and lipid A-stimulated) within clonal populations. Middle: Relationship between the fraction of IL1β responding and CD36 expressing cells per clone and clone size (log₂-transformed) calculated for clonal and control mixed populations. Black circles represent individual clonal populations; blue circles denote control mixed populations across a range of seeding densities (fraction of responding cells averaged per well versus total cell number per well across replicate plates). The black line indicates a linear regression fit for clonal populations; the coefficient of determination (R²) and *p*-value for the non-zero slope are shown. The blue line indicates a linear regression fit for mixed populations. Bottom: Relationship between the fraction of IL1β and CD36 responding cells per clone and cell density (expressed as an average number of cells within 30 μm radius per population) calculated for clonal and control mixed populations (formatted as above). Left: Analysis of IL1β expression in lipid A-stimulated cells, pooled from six biological replicates (see Fig. 3). Right: Analysis of CD36 expression, pooled across treatment conditions and four biological replicates (see Fig. 4). **C.** Daughter cell responses are highly coordinated. Shown are odds ratios (log_10_-transformed) for the binary responses of daughter cell pairs. Circles represent individual replicates/plates, with mean ± SD across biological replicates for IL1β (1750 lipid-A stimulated pairs; data from Fig. 3) and CD36 (1337 untreated and lipid A-stimulated pairs; data from Fig. 4). Statistical significance was assessed using the Mann-Whitney test (\*\**p* < 0.01). **D**. CV of the fraction of IL1β- and CD36-positive cells across clonal populations from B. Solid lines represent bootstrap estimates per bins (Δlog₂N = 1), with dashed lines indicating 95% confidence intervals. **E**. Analysis of IL1β and CD36 responsiveness in clonal populations. Top: Schematic representation of the effect of population size, local cell density and cell morphology on clonal responses. Middle: Summary of fitted linear mixed-effects regression models describing the fraction of responding cells across clonal populations. Models included clone size (log₂N), local density, and their interaction as fixed effects, with plate identity included as a random intercept to account for batch variability. Variance contributions of individual predictors are shown with dominant predictors highlighted in bold. Bottom: Summary of fitted generalized linear models for the odds of a cell being a responder. Models included clone size (log₂N), local density as well as morphological features (nuclear area, nuclear eccentricity, perinuclear area, solidity and *longest_hv_stretch*). Odds ratios (ORs) for a subset of significant predictors are shown, with dominant predictors in bold. Models were fitted using the data described in B. **F**. Schematic representation of the stochastic cell-state switching models for clonal expansion. Cells transition between a transcriptionally inactive OFF state (beige) and an active ON state (blue) governed by transition rates *k_ON_* and *k_OFF_*, with a growth rate λ. Models of increasing complexity were evaluated; (1) constant *k_ON_* and *k_OFF_* rates; (2) *k_ON_* as a function of clone size (N); (3) *k_ON_* as a function of clone size with added extrinsic noise. **G**. Simulated fraction, per clone, of lipid A-stimulated cells expressing IL1β for Model 1 in F. Black circles denote individual simulated clonal populations, matched in number and size to experimental data in B. Black line shows average fraction of positive cells from simulation (constant fit); green line represents the corresponding average from experimental data (linear fit). **H**. Simulated fraction of CD36-expressing cells per clone across models in F. Black circles denote individual simulated clonal populations, matched in number and size to experimental data in B. Black line shows average behaviour of model fits (Model 1: constant; Models 2-3: four-parameter logistic); red line represents fit to experimental data (four-parameter logistic). **I**. CV of the fraction of IL1β- (from G) and CD36-positive cells (from H, Model 3) across simulated clonal populations. Figure formatting as in D.

CD36 exhibited the opposite pattern. Clonal populations showed higher inter-clone heterogeneity (Fig. 5B and C) and elevated daughter-cell correlations in comparison to IL1β (Fig. 5C). The fraction of CD36-positive cells increased with clone size, from ∼10% at log₂N = 6 to ∼40% at log₂N = 10, while inter-clone variability decreased (Fig. 5D). In mixed populations, CD36 expression also increased with cell density, independently of stimulation status (Fig. S10C-D), indicating that local density reinforces CD36 expression and contributes to transcriptional heritability. Notably, however, the density dependence on CD36 expression did not uniformly apply across all clones; a subset of clones characterised by high density exhibited low or none CD36 expression, suggesting that not all cells respond equivalently to population context. Thus, rather than simply driving a uniform shift in expression, density appears to selectively amplify heritable states in a subset of clonal populations.

To assess the influence of population context on clonal expression, we quantified local cell density from the immunofluorescence images by measuring the number of neighbouring cells within a defined proximity of 30 μm (see Methods). Local density varied across datasets (from less than one to approximately 13 neighbouring cells, Fig. S11A) and increased with clone size (R² up to 0.5), indicating that expanding clones experience increasing crowding (Fig. S11B). Density significantly influenced expression across multiple targets, with strongest effects for TNFα, CD36 and IL1β (Fig. S11C). When analysing the fraction of positive cells, CD36 showed consistent associations with both local clone density and clone size (R^2^ = 0.14–0.15), whereas IL1β showed weaker but still significant associations (Fig. 5B).

To disentangle the heritable and environmental contributions, we statistically modelled the fraction of responding cells as function of clone size, local density, and their interaction using multiple linear regression, while controlling for plate-to-plate variability (Fig. 5E and Table S8). For CD36, both clone size (13.8%) and local density (5.7%) contributed to expression, with a positive interaction indicating that local density reinforces CD36 expression in expanding clones. In contrast, IL1β responsiveness was primarily driven by local density, with minimal contribution from clone size. At the single-cell level, logistic regression confirmed local density as the dominant positive predictor of activation of both markers (Fig. 5E, Table S8). Clone size exerted opposing effects, reducing the likelihood of IL1β activation, while increasing CD36 expression. Morphological descriptors suggested an additional regulatory layer, for example, larger perinuclear area suppressed and nuclear size increased responsiveness of both targets.

We used mathematical modelling to recapitulate kinetics of IL1β and CD36 expression during clonal expansion. We implemented a stochastic two-state model in which cells switch between transcriptionally active (ON) and inactive (OFF) states, governed by the transition rates *k_ON_* and *k_OFF_*, and proliferate with rate λ (Fig. 5F). We first considered a model with constant transition rates (Model 1), in which *k_ON_* and *k_OFF_* were estimated from daughter-pair data (see Methods). This model assumes that clonal expansion is independent of clone size and population context and predicts dilution of heritability as clonal size increases. Model 1 qualitatively reproduced the behaviour observed for IL1β (Fig. 5G), however; it predicted a substantially higher steady-state fraction of responding cells than observed in parental populations (∼15% vs 2.5%, respectively; Figure 5B). This discrepancy indicates that population context constrains the steady state distribution of IL1β expression. In contrast, Model 1 failed to recapitulate CD36 dynamics, indicating that constant switching rates derived from daughter-pairs are insufficient to explain the observed behaviour (Fig. 5H). We therefore introduced the dependence of the activation rate *k_ON_* on the population size using Michaelis-Menten kinetics (Model 2, Fig. 5H). Model extension captured the non-linear increase in the ‘average’ CD36 responsiveness during clonal expansion but failed to reproduce the observed inter-clone variability. To account for this, we incorporated extrinsic noise by allowing the maximal activation rate (*k_ON_^max^*) to vary between clones, while remaining constant within each clone (Model 3, Fig. 5H). This model successfully recapitulated the experimental data, demonstrating that CD36 activation can emerge during clonal expansion and can stably propagate through cell divisions, while maintaining high inter-clonal variability (Fig. 5I). Importantly, this extrinsic noise formulation implies that sensitivity to population context is clone-specific and heritable, providing a mechanistic explanation for the observed heterogeneity in the CD36 expression.

### Heritability drives spatial organisation in dense environments

Having characterised two distinct heritable modes for CD36 and IL1β with contrasting regulatory dynamics, we next asked how these states are organized within the same cells and populations. Specifically, we investigated whether these programmes coexist at the single-cell level or constitute mutually exclusive states, as well as how population context influences this relationship. To address this, we examined CD36 and IL1β expression simultaneously, co-staining both targets, in 6-day clonal populations (seeded at 20 cells per well, Fig. 6A) and mixed parental populations, (Fig S12A-C). In both settings, single-cell CD36 and IL1β expression showed a tendency toward mutual exclusivity (Fig. 6B), with individual cells typically expressing one marker but not the other (Fig. 6C, Fig. S12D-E). This negative relationship was also present at clonal level where average CD36 and IL1β expression showed a significant negative correlation, even though there were many clones which expressed both protein targets (Spearman’s ρ = -0.16, p < 0.05, Fig. S12F). This observation suggests that these two states may represent; distinct and potentially competing activation programmes.

**Figure 6.**
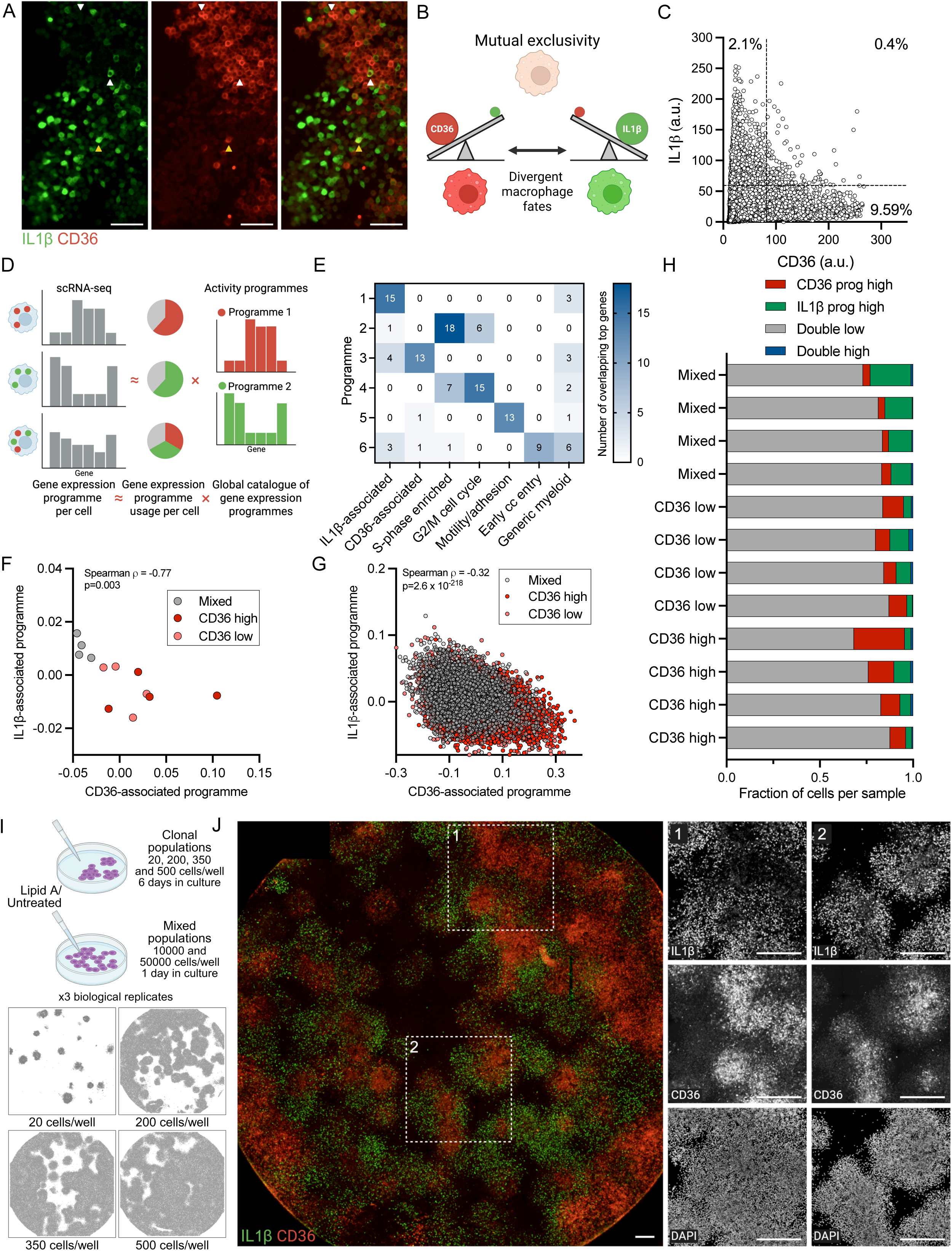
Heritability and cell density drive emergent spatial organisation. **A.** Representative composite fluorescence images of CD36 (red) and IL1β (green) staining in clonal populations (seeded at 20 cells per well and expanded for 6 days) after stimulation with 500 ng/ml lipid A for 3 hours. Data representative of three biological replicates. White arrowheads pointing up: representative example of high-IL1β expressing cell; white arrowhead pointing down: representative example of high-CD36 expressing cell; yellow arrowhead: high-IL1β and -CD36 expressing cell. Scale bar 50 µm. **B.** Schematic representation of heritable exclusivity suggesting that CD36 and IL1β expression may represent reciprocal activation programmes. **C.** Cell-level relationship between CD36 and IL1β protein expression. Shown are single cell fluorescent levels across clonal populations from A after Lipid A stimulation. Dashed lines indicate thresholds for top 2.5% IL1β and 10% CD36 expressing cells, respectively. Percentage of CD36 high, IL1β high and double high expressing cells highlighted. **D.** Schematic representation of NMF-decomposition of scRNA-seq data on lipid A-stimulated clonal and mixed populations into latent transcriptional programmes. **E.** Overlap between NMF-derived programmes and curated gene sets. Heatmap shows the number of overlapping genes between each inferred programme and reference signatures, including IL1β-associated inflammatory response, CD36-associated remodelling, cell cycle phases (S-phase enriched, G2/M), motility/adhesion, early replication, and generic myeloid identity. **F.** Sample-level relationship between CD36 and IL1β programme usage after regression for the cell cycle effect. Each point represents an average per sample coloured by group (CD36 high, CD36 low, mixed). Correlation between CD36 and IL1β programme activity assessed using Spearman’s ρ (p-value indicated). **G.** Cell-level relationship between CD36 and IL1β programme usage after regression for the cell cycle effect. Figure formatting as in F. Spearman’s ρ (p-value indicated). **H.** Distribution of programme-defined cellular states across samples. Cells were classified based on CD36 and IL1β programmes into four states. Cells in the top 10% were defined as programme high: CD36 programme high, IL1β programme high, double high, double low. Bars represent the fraction of cells in each state per sample, grouped by sample. **I.** Experimental design to analyse CD36 and IL1β expression in clonally derived populations embedded within dense environments. iBMDM cells were seeded in 96-well plates to generate low-density (20 cells per well) and high-density (200, 350 and 500 cells per well) clonal populations during 6 days of culture. Mixed (parental) populations were seeded at 10,000 and 50,000 cells per well and cultured for 24 h. All samples were assayed either untreated or stimulated with lipid A (500 ng/ml, 3 h). Shown are segmented cells across representative wells for the clonal populations based on three biological replicates. **J.** Representative fluorescence images of CD36 (red) and IL1β (green) staining for high-density clonal populations (seeded at 500 cells per well) assayed on day 6 following lipid A stimulation. Shown is a composite image of a single representative well and the individual microscopy channels (DAPI, CD36 and IL1β) for the highlighted insert regions (dashed boxes). Data representative of three biological replicates. Scale bar, 100 μm.

To determine whether this relationship reflects underlying transcriptional regulation, we next analysed our scRNA-seq dataset of clonal and mixed populations. In agreement with imaging analyses, at the single cell-level, CD36 and IL1β expression exhibited a tendency towards mutual exclusivity, with the top 10% of CD36-expressing cells characterised by low IL1β expression, and *vice versa* (Fig. S13A). A similar inverse relationship was also observed at the sample level using population averaged expression, where high-CD36 clones were associated with low IL1β expression (Fig S13B).

To identify transcriptional programmes underlying this heterogeneity, we applied non-negative matrix factorisation (NMF) [83] (Fig. 6D). This analysis revealed two robust and distinct programmes associated with CD36 and IL1β, respectively, based on gene loadings (Fig. 6E, Fig. S13C-I, Table S9). The CD36-associated programme included genes such as *Ctsk, Gpnmb, Hmox1, Igf1* and *Metrnl* consistent with a tissue-remodelling and lipid-associated macrophage state [84, 85]. In contrast, the IL1β-associated programme was defined by genes including *Acod1, Tnfa, Nlrp3, Cxcl2*, and *Nfkbia*, representing a canonical inflammatory NF-κB-driven response [5, 65] (Fig. S13C). We also identified a programme associated with adhesion and motility, suggesting further functional diversification, and additional programmes associated with cell cycle progression (Fig. 6E, Table S9). CD36- and IL1β-associated programmes were significantly negatively correlated across both samples and cells, before (Fig. S13J-K) and after (Fig. 6F-G) cell-cycle regression, and showed depletion of co-activation at high expression levels (Fig. 6H) as suggested by imaging data, indicating that they represent largely independent transcriptional axes. Importantly, this reciprocal regulation between CD36- associated lipid remodelling and IL1β-associated inflammatory macrophage states has also been previously observed across multiple systems. In human induced pluripotent stem cell (iPSC)-derived macrophages, a model of tissue resident macrophages [82], both CD36 and IL1β exhibited reciprocal density-dependent regulation, which was mirrored by associated programme genes (Fig. S13L). A similar reciprocal organisation of CD36-associated lipid remodelling and inflammatory macrophage programmes has also been described in atherosclerosis [86, 87] and in tumour associated macrophages, where expression of CD36 is associated with suppression of pro-inflammatory responses [88].

Having established that CD36 and IL1β define opposing heritable transcriptional programmes, we next asked how population context influences their relationship. Previous work has shown that cell density can attenuate transcriptional heritability through population-level feedback mechanisms, including quorum sensing, in the early interferon responses [6]. This, in addition to the observation of density-reinforced heritability of CD36 expression and attenuated IL1β responses (Fig. 5), prompted us to examine whether increasing cell density could alter the distribution of these transcriptional states.

To move beyond the behaviour of isolated clones, we generated clonal populations seeded at high density in 96-well plates (200, 350 and 500 cells per well), alongside low-density clones (20 cells per well), that were expanded for 6 days. This design introduced local high-density cell-to-cell interactions between neighbouring clones. Mixed-population controls (seeded at 10,000 and 50,000 cells per well, 3.1×10⁴–1.5×10^5^ cells/cm²) were analysed in parallel (Fig. 6I, Fig. S14A-C). Under high-density conditions, CD36 protein expression formed discrete spatial pockets corresponding to locally dense regions, likely derived from founder cells (Fig. 6J). These regions were embedded within a continuous monolayer spanning most of the well area, consistent with merging neighbouring clones. In contrast, IL1β was relatively depleted within high-CD36 regions and enriched in surrounding cells following lipid A stimulation.

To quantify this spatial cross-correlation, we applied bivariate Moran’s I, a global spatial-autocorrelation statistic that quantifies whether high values of one variable occur preferentially near high (or low) values of another [89]. Using single-cell CD36 and IL1β intensities, we constructed a spatial-weights matrix based on neighbouring cells (100 for high-density clonal and mixed populations, and 20 for low-density clones, consistent with the local analyses) and computed bivariate Moran’s I for each well across technical and biological replicates (Fig. S14D). This analysis revealed a strong negative spatial correlation between CD36 and IL1β in high-density clonal populations following stimulation (e.g. I = -0.54 ± 0.09 for 500 cell per well seeding), confirming that CD36-enriched regions are surrounded by IL1β-enriched neighbouring cells, rather than co-localising with them. This significant negative relationship, although weaker, was also observed in stimulated mixed populations, especially at 50,000 cells per well seeding density (I = -0.24 ± 0.05). As expected, this correlation was absent in the non-treated samples, due to the low basal expression of IL1β (Fig. S14D). Thus, while CD36 and IL1β programmes are mutually exclusive at the single-cell level in parental populations and low-density clones, their spatial segregation emerges specifically in high density environments, giving rise to stable, locally organized niches.

Taken together, these findings suggest that both transcriptional heritability and population context interact to organize macrophage responses into spatially structured and transcriptionally distinct states.

### Heritability in CD36 expression regulates susceptibility to *L. monocytogenes*

Our findings demonstrate that innate immune responses are shaped by heritable transcriptional states that persist across multiple cell divisions. Such sustained differences in activation state, cytokine production, and receptor expression may have important implications for bacterial infection dynamics [6, 34, 90]. In proliferating primary bone marrow-derived and tissue-resident macrophages [49–52], these inherited states are predicted to create clonal subpopulations with distinct capacities to sense, internalise, and restrict invading pathogens. To directly test whether transcriptional heritability translates into functional differences in susceptibility to bacterial infection, we focused on CD36 and examined its role during infection with the important foodborne bacteria *L. monocytogenes* [91]. The ability of *Lm* to cause systemic infection depends on its capacity to enter and subvert macrophage activation. Following the receptor-mediated uptake, *Lm* rapidly escapes the phagosome to replicate within the cytosol [92]. The efficiency of these early invasion steps is strongly influenced by the repertoire and abundance of host surface receptors [93–96]. Among these, CD36 has emerged as a key facilitator of the internalisation of Gram-positive bacteria, including *Lm*, through its ability to bind bacterial lipoteichoic acids and to organise lipid microdomains required for efficient uptake in phagocytic cells [56, 97].

To assess the functional contribution of CD36 to *Lm* infection, we infected mixed populations of iBMDMs with a fluorescent *Lm*-GFP strain [38]. Cells were seeded at two densities (10,000 or 50,000 cells per well, 3.1×10⁴–1.5×10^5^ cells/cm²), infected at multiplicity of infection (MOI) of 5 for 45 mins, and analysed 24 h post-infection following CD36 immunostaining. In the high-density condition, cells were additionally treated with the previously-characterised specific CD36 inhibitor sulfosuccinimidyl oleate (SSO) [98] (Fig. 7A, Fig. S15A). High magnification microscopy (40x) was used to optimise detection of the *Lm*-GFP signal, and images were processed using our established analysis pipeline (Fig. S15B). Infection resulted in robust intracellular replication and cell-to-cell spread. Higher seeding density significantly increased infection burden, as quantified by average perinuclear *Lm*-GFP intensity per cell, whereas low-density cultures exhibited fewer replicative events and reduced bacterial spread (Fig. 7B). Importantly, SSO treatment markedly reduced infection levels (Fig. 7A-B, Fig. S15C), consistent with CD36 contributing to bacterial uptake [97]. In parallel, CD36 expression showed a strong density-dependent upregulation (Fig. 7C), in line with our earlier analyses. Notably, SSO treatment did not significantly alter CD36 protein abundance, consistent with its known mechanism of action: irreversible modification of Lys164 on CD36 that blocks ligand engagement without affecting receptor expression [98].

**Figure 7.**
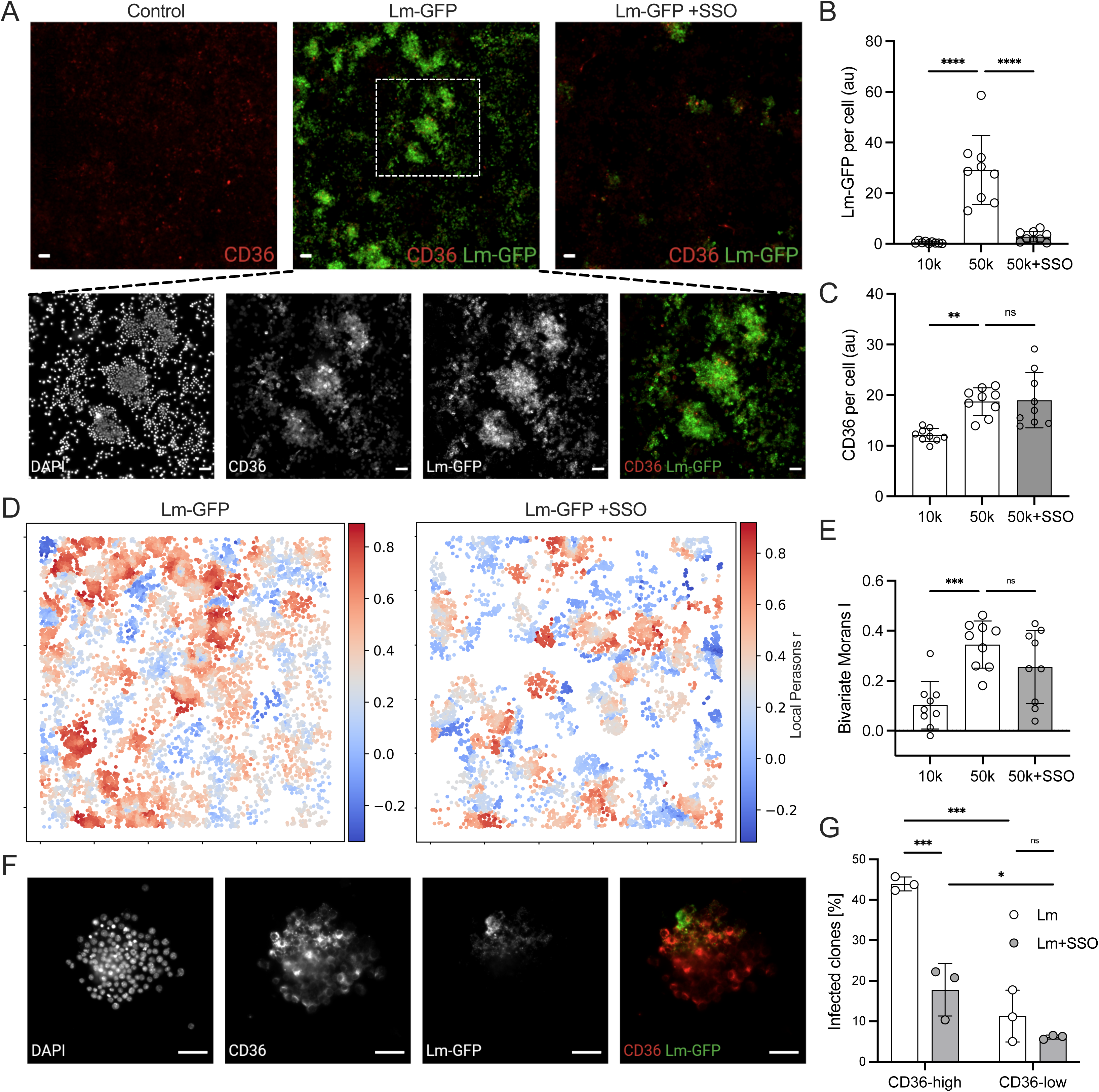
Heritability of CD36 expression affects susceptibility to *L. monocytogenes*. **A.** CD36 expression determines *Lm* infection course. Mixed iBMDM populations were seeded at two densities (10,000 or 50,000 cells per well), left uninfected (control) or infected with *Lm*-GFP at MOI = 5 for 45 min, and immunoassayed for CD36 at 24 h post-infection. High density cells were infected in the absence or presence of 200 µM of the CD36 inhibitor SSO. Top: Representative composite fluorescence images of CD36 staining (red) and *Lm*-GFP (green) across indicated conditions for 50,000 cells seeded per well. Bottom: Individual microscopy channels (DAPI, CD36 and *Lm*-GFP) and the corresponding composite image for CD36 and *Lm*-GFP for the insert region highlighted above (dashed box). Data representative of three biological replicates. Scale bar, 50 μm. **B.** SSO reduces infection burden. Shown is the mean single-cell perinuclear *Lm*-GFP signal quantified per well (individual wells shown as circles), with mean ± SD across three biological replicates/plates for two cell densities infected with *Lm* and treated with SSO (as in A). Statistical significance assessed using ordinary one-way ANOVA with Dunnett’s correction for multiple comparisons (\*\*\*\**p* < 0.0001). **C.** Mean single-cell perinuclear CD36 signal quantified per well (individual wells shown as circles), with mean ± SD across three biological replicates/plates (as in B). Statistical significance assessed using ordinary one-way ANOVA with Dunnett’s correction for multiple comparisons (ns - non-significant, \*\**p* < 0.01). **D.** Local linear association between *Lm*-GFP and CD36 perinuclear signals. Shown are local correlation maps (Pearson’s r) computed with a KNN method (k = 100). Analyses are shown for the microscopy images in panel A (*Lm*-GFP, *Lm*-GFP + SSO) and Fig. S15C. Experimental conditions as in panel A. **E.** Spatial relationship between *Lm*-GFP intensity and CD36 expression. Shown is the global spatial-autocorrelation (bivariate Moran’s I) between perinuclear *Lm*-GFP and CD36 signals using a kNN method (*k* = 100). Individual wells are shown as circles, with mean ± SD across three biological replicates/plates. Statistical significance assessed using ordinary one-way ANOVA with Dunnett’s correction for multiple comparisons (ns – non-significant, \*\*\**p* < 0.001). **F.** Analysis of infection susceptibility in CD36 clonal populations. iBMDMs were seeded at 20 cells per well and expanded for six days to generate clonal populations. Clones were then infected with *Lm*-GFP at MOI = 5 in the absence or presence of 200 µM SSO, and stained for CD36 24 h post-infection. Shown are individual microscopy channels (DAPI, CD36, and *Lm*- GFP) and the corresponding composite image for CD36 and *Lm*-GFP for a representative CD36-high *Lm*-infected clone. Data are representative of three biological replicates/plates. Scale bar: 50 µm. **G.** CD36 expression increases susceptibility to *Lm*. Shown is the average fraction of *Lm*-infected clones stratified into CD36-high or CD36-low (individual plates shown as circles), with mean ± SD across three biological replicates/plates. Experimental conditions correspond to those in panel F. Statistical significance assessed using ordinary two-way ANOVA with Holm-Sidak correction for multiple comparisons (ns – non- significant, \**p* < 0.05, \*\*\**p* < 0.001).

Infected cultures exhibited striking spatial organisation, particularly at high seeding density. *Lm*-GFP formed discrete multicellular pockets, or infection foci, in which bacteria established replicative niches and subsequently spread to neighbouring cells [99] (see the insert in Fig. 7A). These *Lm*-GFP infection foci spatially coincided with regions of elevated CD36 expression. Notably, this spatial enrichment persisted in SSO-treated wells, where the remaining few localised infections were likewise concentrated in CD36-high cells (Fig. S15C). To quantify this structure at the microscale, we computed local Pearson’s correlation coefficient (r), measuring the linear association between *Lm*-GFP intensity and CD36 expression levels within local spatial neighbourhoods. Local-correlation maps revealed predominant high-infection/high-CD36 hotspots (local r approaching 0.8), corresponding to multicellular infection foci embedded within CD36-rich microenvironments (Fig. 7D). In contrast, high-infection/low-CD36 regions were rare, indicating that infection is preferentially sustained in CD36-enriched regions. We observed a strong positive spatial association at high seeding density (bivariate Moran’s I = 0.34 ± 0.09), whereas spatial coupling was significantly weaker at low density (I = 0.10 ± 0.09), consistent with reduced formation of multicellular infection foci (Fig. 7E). SSO treatment decreased global spatial association (I = 0.26 ± 0.15), indicating partial disruption of CD36-dependent spatial coupling. Nevertheless, infection remained preferentially localised in CD36-high cells, likely reflecting incomplete inhibition of CD36 function (Fig. 7E). Collectively, these analyses demonstrate that local CD36 expression structures the spatial organisation of *Lm* infection, supporting a microenvironment-level mechanism by which CD36 expression shapes infection dynamics.

Finally, to directly test whether clonal variation in CD36 expression determines susceptibility to *Lm*, we generated clonal iBMDM populations by seeding 20 cells per well and expanding them for six days prior to infection. Clones were then infected with *Lm*-GFP for 24 h at an MOI of 5, to ensure that each clone had multiple interactions with *Lm*, in the absence or presence of SSO. Across three biological replicates, we analysed 623 clones (438 untreated, and 185 SSO-treated), which were manually stratified into CD36-high and CD36-low groups and scored for the presence or absence of replicative *Lm* foci, defined as regions containing large numbers of bacteria distributed across multiple neighbouring host cells (Fig. 7F). Approximately 50% of the clones were classified as CD36-low, consistent with our previous quantification of CD36 responsiveness (Fig. 5B). Strikingly, 43.9 ± 1.7% of CD36-high clones supported replicative infection, compared to only 11.3 ± 6.4% of CD36-low clones (Fig. 7G). SSO treatment significantly reduced infection among CD36-high clones to 17.7 ± 6.5%, confirming the CD36-dependent mechanism. CD36-low clones treated with SSO showed detectable infection, 6.1 ± 0.5%, consistent with partial inhibition or CD36-independent entry mechanisms.

Taken together, these results demonstrate that transcriptionally inherited variation in CD36 expression determined clonal susceptibility to *Lm* infection, providing direct functional evidence that transcriptional heritability shapes outcomes of host-pathogen interactions.

## Discussion

Innate immune activation is inherently heterogeneous; genetically identical cells differ in cytokine production [11, 73] and infection control [14, 37, 38]. While such variability is often attributed to stochastic signalling [4, 8, 29, 30, 100, 101], accumulating evidence suggests that gene expression states persist through cell divisions [16, 32]. Here we demonstrate that TLR-driven responses exhibit transcriptional heritability, and that this heritability interacts with population context to shape functional immune outcomes.

Using fluctuation-based genomics, we identified a subset of TLR-responsive genes whose expression exhibits long term transcriptional heritability in clonal iBMDMs. Heritability was enriched among inflammatory mediators, receptors and negative-feedback regulators such as A20, SOCS-3 and IRF1, whereas core pathway components such as NF-κB and JAK-STAT systems were largely non-heritable (Fig. 1). This suggests that transcriptional heritability may preferably operate at the level of regulatory tuning rather than primary signal propagation [6, 102, 103]. Additional heritable loci including *Nlrp3* and *Ptgs2*, raise the possibility that binary inflammasome activation is also imprinted [104]. Concordance between bulk cell MemorySeq, scRNA-seq (Fig. 2) and high content imaging (Figs. 3–4) demonstrates that transcriptional heritability is mirrored at the protein level across at least 10 generations. Strong inter-daughter cell correlations (Pearson’s r from 0.55 to 0.85) indicate that early clonal divergence arises from inherited expression states that progressively dilute as clones expand. However, subsets of genes including TNFα and IL1β may retain heritable signatures for longer than others.

Local environment is a critical regulator of macrophage activation, with cell density known to shape cytokine output and signalling thresholds [9, 80–82]. We found that population context modulates heritability and identified two regulatory regimes (Fig. 5). CD36 exemplifies a stable, heritable programme that is reinforced by environmental feedback; as local cell density increases, it amplifies pre-existing CD36 states, increasing clonal divergence. Critically, while mixed populations display a density effect, approximately half of the clonal populations fail to express CD36 after several divisions under high density, suggesting that the capacity for density-dependent CD36 induction must itself be an inherited state. This observation is consistent with our mathematical modelling analyses. In contrast, IL1β exemplifies a rapid stimuli-induced inflammatory response that is shaped by environmental context in an opposite manner; as clonal populations expand, the fraction of IL1β-positive cells declines. Although IL1β activation is correlated between daughter cells, the overall fraction of responding cells across clonal populations is substantially lower than predicted from these daughter cell-pair rates. This indicates that early heritable activation states are progressively constrained at the population level as density increases. A similar regulatory logic has been described for IFN first responders, where the fraction of responding cells is limited by population size through a quorum sensing mechanism [6].

Surprisingly, we found that different heritability modes are interconnected; at the single-cell level, expression of IL1β and CD36 are mutually exclusive, whereas under high-density conditions, they give rise to spatial organisation within the cellular monolayer (Fig. 6). Our scRNA-seq data analyses reveal that the CD36-associated and IL1β-associated programmes are transcriptionally orthogonal, consistently with previous analyses of macrophage heterogeneity in different cell systems. Studies in atherosclerosis identified distinct inflammatory IL1β-high macrophages and foamy CD36-high macrophages enriched for lipid metabolism. In murine plaques, nonfoamy macrophages preferentially expressed inflammatory genes including *Il1b*, *Nfkbia*, *Tlr2* and *Tnfa*, whereas foamy macrophages expressed *Cd36*, *Mertk* and *Nr1h3* [86]. Similarly, multimodal single-cell profiling of human carotid atherosclerosis identified IL1β-high macrophages alongside foamy macrophages expressing CD36 [87]. A related reciprocal organisation has also been described in tumour-associated macrophages, where CD36 expression was associated with immunosuppressive and tissue-remodelling phenotypes, while loss of CD36 promoted inflammatory macrophage activation increasing IL1β and IFNβ production and delaying tumour growth [88]. Mechanistically, this reciprocal regulation may be consistent with a regulatory axis in which NF-E2-related factor-2 (NRF2) promotes a CD36-associated macrophage state while suppressing pro-inflammatory cytokine programmes. NRF2 has been shown to directly repress transcription of inflammatory genes, including IL1β, by limiting polymerase II recruitment [105], while directly regulating CD36 expression through antioxidant response elements in its promoter [106]. However, contributions of other CD36-regulators, including PPARγ [107] or signalling downstream of CD36, such as p38 MAPK [88] cannot be excluded.

Transcriptionally inherited states have clear functional consequences. Heritable CD36 expression creates clonal lineages biased toward high or low receptor abundance, and generates spatial organisation in homogeneous macrophage populations, spatially pre-patterning macrophage susceptibility to infection. Consistent with this, CD36-positive clones were significantly more susceptible to *L. monocytogenes* (Fig. 7), directly linking transcriptional heritability to divergent infection outcomes [37, 39, 44–46]. Density-dependent reinforcement of CD36 expression has previously been observed in a human stem-cell derived model of tissue-resident macrophages [82], and may have broader physiological implications. CD36 participates in phagocytosis of Gram-positive bacteria [78], recognition of *Plasmodium falciparum*-infected erythrocytes [108], and uptake of oxidised LDL during foam-cell formation in atherogenesis [109]. Tightly packed macrophage clusters may therefore function as self-amplifying niches with altered lipid handling and inflammatory signalling, potentially regulating susceptibility to infections and chronic inflammatory pathology including cancer. In turn, the IL1β-associated programme reflects the canonical inflammatory response to innate immune stimulation. However, this programme is also closely related to macrophage states implicated in pathogenic inflammation, where IL1β signalling has established therapeutic relevance [110, 111]. Beyond CD36, heritable expression extends to additional phagocytic receptors (*Msr1* [93], *Fcgr2b* [112], complement *C1q* complex [72]) and to downstream antimicrobial machinery (*Marcks, Myo1d, Plekho1, Slc11a1, Rab20, Atp6v0d2, Clcn7, Lyz1/2, Ctsc*) (Table S1) [70], suggesting that pathogen recognition and intracellular processing could be clonally predetermined.

Mechanistically, heritable genes exhibited coordinated expression patterns, consistent with shared regulatory architecture [16]. We observed blocks of co-regulated genes in MemorySeq and, albeit with greater noise, in the scRNA-seq dataset (Fig. S4), suggesting a higher-order regulatory control. For CD36, canonical regulators such as *Pparγ* or *Nrf2* [109] were not themselves heritable or correlated with its expression, implying that additional regulatory mechanisms may underlie its heritable and density-dependent expression. Epigenetic inheritance offers a plausible explanation, as specific histone modifications can be mitotically inherited to confer long-term transcriptional memory [113]. More broadly, while current models of single-cell transcriptional bursting assume stochastic activation within homogeneous populations [4, 29–31], our data support a framework in which slow, epigenetically driven transitions establish latent heritable states that are subsequently modulated via population context. Incorporating such slow state transitions and density-dependent modifiers into stochastic transcription models will be essential for capturing the architecture of transcriptional heritability.

The use of immortalised BMDMS enabled quantification of transcriptional heritability through multiple cell divisions, providing an experimental framework that is not feasible in primary *in vitro* macrophage systems [14, 61]. Notably, several of the macrophage programmes identified here are consistent with observations in primary and tissue-derived macrophages populations, including reciprocal CD36- and IL1β-associated states described in atherosclerosis and tumour environments [86–88]. Primary and tissue-resident macrophages, including microglia, proliferate during steady-state and inflammation [49–53], providing opportunities for heritable states to propagate *in vivo*. For example, *L. monocytogenes* induces local proliferation of monocyte-derived macrophages in the liver [54]. Critically, phenotypic heritability has been directly demonstrated for proliferating microglia, which adopt clonally-restricted electrophysiological phenotypes following cerebral ischemia [55]. Heritable states may therefore represent a mechanism for maintaining persistent population-level heterogeneity within tissues [114]. Another possibility is that tissue-resident macrophages capable of restricting pathogen growth may propagate their heritable expression states, promoting lineage-level resistance within local tissues. Future lineage tracing and fate-mapping studies [55, 115, 116] will be required to determine how such stable states arise and propagate *in vivo*.

Overall, we demonstrate that acute TLR signalling can exhibit gene-specific transcription heritability that persists across multiple generations. By integrating fluctuation-based genomics, single-cell transcriptomics, high-content imaging, and functional infection assays, we provide a quantitative framework linking heritable gene expression states to macrophage phenotypes. These findings show how inherited transcriptional programmes and microenvironmental cues jointly shape infection heterogeneity.

## Materials and Methods

### Cell lines and culture

iBMDMs [61] were cultured in Dulbecco’s modified Eagle’s medium (DMEM) high glucose with sodium pyruvate and sodium bicarbonate (Sigma-Aldrich, D6429) supplemented with 10% heat inactivated foetal bovine serum (FBS, Gibco, 10500064) and 1% (w/v) penicillin/streptomycin (Gibco, 15070063). Cells were incubated at 37°C and 5% (w/v) CO_2_, under laminar air flow unless otherwise stated. Lipid A diphosphoryl from *Salmonella enterica* serotype Minnesota Re595 (Sigma-Aldrich, L0774) stock was prepared at 500 µg/ml in 50% DMSO and stored at -20°C until use.

### Bacterial strain culture conditions

*Lm* EGDe inlA BUG2479 pPL2 Phyper-GFP (*Lm*-GFP) [38] was grown in tryptone soya broth (TSB, Oxoid, CM0129B) with 7 µg/ml of chloramphenicol (Sigma Aldrich, C0378). *Lm*-GFP mid-log phase aliquots (OD_600_ 0.5–0.6), stored at -80 °C in 15% (v/v) glycerol (Sigma Aldrich, G5516) in PBS, were used for infection. Bacterial counts (CFU/ml) were determined by plating serial dilutions onto TSB agar (Oxoid, CM0131B) supplemented with chloramphenicol.

### Doubling time analysis

iBMDM cells were regularly passaged over a three-month period, and for each passage the number of seeded (*N_0_*) and harvested (*N*) cells, as well as the elapsed time (*t*, in hours), were recorded (see Table S7). The doubling time, *T_d_* was calculated according to the exponential growth formula 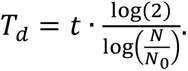 The number of doublings required to reach a given population size (N_c_) was determined as *n* = *log*_2_(*N*_*c*_). The largest clones obtained in our assays (1,000–1,500 cells) correspond to approximately 10.0–10.6 doublings over the 6-day period. This is consistent with the estimated average doubling time of iBMDMs (14.4 ± 1.51 h), which predicts theoretical clonal population sizes of ∼540–2,300 cells after 6 days. In comparison, we estimate an average 27 doubling times for clonal populations used for RNA-seq assays at day 16.

### Preparation of clonal populations for RNA-seq analyses

Single iBMDMs were FACS-sorted into prewarmed complete DMEM in 96-well plates and incubated to generate clonal populations. To reduce clumping during growth from single cells, cells were dispersed by gently reverse pipetting on days 5 and 8. After reaching 60-80% confluency (day 12-14), clonal populations were transferred to 6-well plates and cultured until they reached 100,000-200,000 cells per well (day 15-16) for subsequent experiments. Clonal or mixed populations at approximately 50% confluence were stimulated with 500 ng/ml lipid A for 3h. Untreated controls were treated with an equivalent volume of 50% DMSO.

### MemorySeq library construction, sequencing, and differential expression analysis

43 stimulated clonal populations, 43 stimulated mixed populations and 10 untreated mixed populations were prepared. Cells were harvested by scraping and stored in RNA*later* (Invitrogen, AM7020) for stable storage. Cells were pelleted by centrifugation and RNA*later was* removed prior to RNA extraction with the RNeasy Plus mini kit (Qiagen, 74136) according to the manufacturer’s instructions. RNA was stored at -80°C. Total RNA was submitted to the Genomic Technologies Core Facility (GTCF). Quality and integrity of the RNA samples were assessed using a 4200 TapeStation (Agilent Technologies) and then libraries generated using the Illumina® Stranded mRNA Prep. Ligation kit (Illumina, Inc.) according to the manufacturer’s protocol. Briefly, total RNA (typically 0.025-1 µg) was used as input material from which polyadenylated mRNA was purified using poly-T, oligo-attached, magnetic beads. Next, the mRNA was fragmented under elevated temperature and then reverse transcribed into first strand cDNA using random hexamer primers in the presence of Actinomycin D to improve strand specificity whilst mitigating spurious DNA-dependent synthesis. Following removal of the template RNA, second strand cDNA was then synthesized to yield blunt-ended, double-stranded cDNA fragments. Strand specificity was maintained by the incorporation of deoxyuridine triphosphate (dUTP) in place of dTTP to quench the second strand during subsequent amplification. Following a single adenine (A) base addition, adapters with a corresponding, complementary thymine (T) overhang were ligated to the cDNA fragments. Pre-index anchors were then ligated to the ends of the double-stranded cDNA fragments to prepare them for dual indexing. A subsequent PCR amplification step added the index adapter sequences to create the final cDNA library. The adapter indices enabled the multiplexing of the libraries, which were pooled prior to clustering on a flow-cell from a High-Output NextSeq 500/550 v2.5 kit. The loaded flow-cell was then paired-end sequenced (76 + 76 cycles, plus indices) on an Illumina NextSeq500 instrument. Finally, the output data was demultiplexed and the BCL-to-Fastq conversion was performed using Illumina’s bcl2fastq software, version 2.20.0.422.

Unmapped paired-reads of 74bp from an Illumina NextSeq500 sequencer were interrogated using a quality control pipeline consisting of FastQC v0.11.3 (http://www.bioinformatics.babraham.ac.uk/projects/fastqc/) and FastQ Screen v0.13.0 (https://www.bioinformatics.babraham.ac.uk/projects/fastq_screen/). The reads were trimmed to remove any adapter or poor quality sequence using Trimmomatic v0.39 [117]; reads were truncated at a sliding 4bp window, starting 5’, with a mean quality <Q20, and removed if the final length was less than 36bp. Additional flags included: ‘ILLUMINACLIP:./Truseq3-PE-2_Nextera-PE.fa:2:30:10 SLIDINGWINDOW:4:20 MINLEN:36’.The filtered reads were mapped to the mouse reference sequence (mm10/GRCm38) from the UCSC browser [118], using STAR v2.7.7a [119]. The genome index was created using the comprehensive mouse Gencode vM25 gene annotation [120] applying a read overhang (--sjdbOverhang 75). During mapping, the flags ‘--quantMode GeneCounts’ were used to generate read counts into genes. Normalisation and differential expression analysis was performed using DESeq2 v1.30.1 [121] on R v4.0.4. Log fold change shrinkage was applied using the lfcShrink function along with the “apeglm” algorithm [122].

Following mapping and analyses we retained all but 2 clonal samples with a depth of at least 500,000 reads per library (2.9 million reads per sample on average). One sample was removed due to poor coverage, while another (sample 18) was subsequently removed because it clustered with mixed cell populations (Fig. S1).

### scRNA-seq single cell isolation, library construction and sequencing

8 stimulated clonal populations and 4 stimulated mixed populations were prepared. Cells were harvested by scraping, and then clonal populations were labelled prior to pooling with the 10x genomics 3’ CellPlex kit according to the manufacturer’s instructions for >80% viable cells (protocol 1 document number CG000391). Library preparation and sequencing was performed by the University of Manchester Genomic Technologies Core Facility. Gene expression and Cell Multiplexing libraries were prepared from Cell Multiplexing Oligo labelled cells using the Chromium Controller and Single Cell 3ʹ Reagent Kits v3.1 (10x Genomics, Inc. Pleasanton) according to the manufacturer’s protocol (CG000388 Rev B). Briefly, nanoliter-scale Gel Beads-in-emulsion (GEMs) were generated by combining barcoded Gel Beads, a master mix containing cells, and partitioning oil onto a Chromium chip. Cells were delivered at a limiting dilution, such that the majority (90-99%) of generated GEMs contain no cell, while the remainder largely contain a single cell. The Gel Beads were then dissolved, primers released, and any co-partitioned cells lysed.

Primers containing an Illumina TruSeq Read 1 sequencing primer, a 16-nucleotide 10x Barcode, a 12-nucleotide unique molecular identifier (UMI) and a 30-nucleotide poly(dT) sequence were then mixed with the cell lysate and a master mix containing reverse transcription (RT) reagents along with primers containing an Illumina Nextera Read 1, a 16-nucleotide 10x Barcode, a 12-nucleotide UMI and a capture sequence. Incubation of the GEMs yielded barcoded cDNA from poly-adenylated mRNA and barcoded DNA from the Cell Multiplexing Oligo Feature barcode. Following incubation, GEMs were broken, and pooled fractions recovered. The cell barcoded cDNA was then purified from the post GEM-RT reaction mixture using silane magnetic beads and amplified via PCR to generate sufficient mass for library constructions. The amplified cDNA molecules for 3’ Gene Expression were separated from those for Cell Multiplexing library construction by size selection. For the 3’ Gene Expression library enzymatic fragmentation and size selection were then used to optimize the cDNA amplicon size. Illumina P5, P7, i7 & i5 sample indexes, and TruSeq Read 2 sequence were added via end repair, A-tailing, adaptor ligation, and PCR to yield final Illumina-compatible sequencing libraries. For the Cell Multiplexing library P5, P7, i7 & i5 sample indexes and Nextera Read 2 were added via PCR to amplified DNA from Cell Multiplexing Oligo Feature Barcodes to yield final Illumina-compatible sequencing libraries.

Single Cell 3’ libraries comprised standard Illumina paired-end constructs flanked with P5 and P7 sequences. The 16 bp 10x Barcode and 12 bp UMI were encoded in Read 1, while Read 2 was used to sequence the cDNA fragment in 3’ Gene Expression libraries while read 2N was used to sequence the DNA from Cell Multiplexing Feature barcode. Sample index sequences were incorporated as the i7 and i5 index read. Paired-end sequencing (28:90) was performed on the Illumina NovaSeq 6000 platform. The .bcl sequence data were processed for QC purposes using bcl2fastq software (v. 2.20.0.422) and the resulting .fastq files assessed using FastQC (v. 0.11.3), FastqScreen (v. 0.14.0) and FastqStrand (v. 1.11.1) prior to pre-processing with the CellRanger pipeline.

### scRNA-seq data processing, cell filtering and cell cycle assignment

The 10x Genomics Cell Ranger pipeline (v7.0.0) was used to process raw sequencing data. The base call (BCL) files produced by the sequencer were demultiplexed and converted to FASTQ files using “cellranger mkfastq”. The gene expression and multiplexing capture FASTQ files were processed using “cellranger multi” and mapped against the pre-built Mouse reference package from 10x Genomics (mm10-2020-A) to generate the per-sample gene-cell barcode matrix. The single-cell data were processed in the R environment (v4.1) following the workflow documented in Orchestrating Single-Cell Analysis with Bioconductor [123]. Briefly, for each sample, the HDF5 file generated by Cell Ranger was imported into R to create a SingleCellExperiment object. A combination of median absolute deviation (MAD), as implemented by the “isOutlier” function in the scuttle R package (v1.4.0) and exact thresholds was used to identify and subsequently remove low quality cells before data integration. Cell cycle phase classification was performed using the “cyclone” function and pre-trained classifiers from the scran R package (v1.22.1) and obtained the predicted phase for each cell.

### scRNA-seq data integration, visualisation, and cell clustering

The “multiBatchNorm” function from the batchelor R package (v1.10.0) was used to re-compute the log-normalized expression values of the combined single-cell data. Mutual nearest neighbours (MNN) approach available from the batchelor R package was used to perform batch correction on top 2000 highly variable genes. Then, the first 50 dimensions of the MNN low-dimensional corrected coordinates were used as input to produce the t-stochastic neighbour embedding (t-SNE) projection using the “runTSNE” function from the scatter R package (v1.22.0) respectively.

### Heritability analysis for RNA-seq data

Heritability metric was defined as the ratio of coefficient of variations (CV) of the log_2_(read counts +1) in the clonal and mixed samples, respectively. In the MemorySeq analysis this metric was calculated for 11617 genes with a total number of read counts >100 across all samples. In the scRNA-seq, the metric was calculated based on average UMIs per sample for 5701 genes that were also expressed in MemorySeq (genes with at least one clonal sample with log_10_(UMIs)>1 were considered expressed, with total of 5791 genes passing the threshold). Cut-offs of 2.5 and 3 for MemorySeq and noisier scRNA-seq data, respectively, were applied to define heritable genes. Gene ontology (GO) Biological Process enrichment analysis was performed using g:Profiler (gprofiler2 v0.2.3, Ensembl gene sets, FDR-corrected using the Benjamini-Hochberg method, significance threshold p < 0.05) [124]. For each pair of 73 heritable genes identified in both MemorySeq and scRNA-seq datasets, we calculated the Pearson’s correlation coefficient between their expression across all samples (clonal and mixed). Correlation matrices were represented by heatmaps with the same order of genes (as per MemeorySeq).

### Latent variable decomposition for scRNA-seq data

Genes with low total transcript counts across all conditions (<1000 log_2_UMI) were excluded, retaining 10,631 genes across 9,166 cells. The top 2,000 most variable genes (based on variance across cells) were selected for downstream analysis. To identify transcriptional programmes, NMF [83] was applied to the expression matrix (*cells x genes*) using the skearn.decomposition.NMF implementation in Python. The number of components was set to *k = 6*, based on empirical evaluation of stability across a range of factorisation ranks. NMF decomposed the expression matrix *X* into *X* = *W* × *H*, where *W* represented programme usage per cell (*cells x programmes*) and H represented gene weights per programme (*programmes x cells*). Identified transcriptional programmes were annotated post-hoc based on manual inspection of top programme-defining genes and overlap with curated marker gene sets representing IL1β-associated inflammatory, Cd36-associated remodelling, cell-cycle, motility, and myeloid states (Table S9). Programmes were assigned to the marker set with the highest overlap, subject to a minimum threshold of three shared genes. Curated marker sets included IL1β-associated inflammatory programme (*Il1β, Nfkbia, Tnfα, Cxcl2, Nlrp3, Acod1, Ccl4*) [5, 65], CD36-associated remodelling programme (*Cd36, Ctsk, Gpnmb, Hmox1, Igf1, Metrnl*) [84, 85], S-phase enriched cell cycle reflecting active synthesis and chromatin assembly including histone cluster genes (*Hist1h1b, Hist1h1e, Hist1h2ae*) [125], replication-associated factors (*Pclaf*) [126], cell cycle regulators (*Top2a, Mki67, Cdk1, Cenpf*) [127], G2/M cell cycle programme (*Ube2c, Cdc20, Tpx2, Birc5, Ccnb1)* [128–130], motility/adhesion-like programme (*Itga4, Dock2, Diaph2, Peak1, Ahnak, Calcrl, Pla2g4a, Ralgapa2, Srgap2, and Gphn*) [131–135] and early cell cycle entry programme (*Nasp, Dut, Ccnd1, Pcna, Mcm7*) [136, 137]. To remove the effect of cell-cycle, a linear regression model was used:

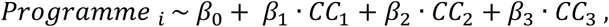

where *CC_1_*, *CC_2_* and *CC_3_* correspond to three cell cycle-related programmes. Residuals form this model were used as cell-cycle corrected programmes.

To determine whether macrophage density influences expression of the CD36- and IL1B-associated transcriptional programmes, we reanalysed the published macrophage density RNA-seq dataset [82]. Gene-level TPM values were extracted for representative genes, samples were grouped according to the original low-, medium-, and high-density culture conditions.

### Preparation of clonal and mixed populations for imaging analyses

For experiments with clonal populations, iBMDM cells were seeded in flat bottom 96-well TC treated imaging plates (Greiner, 655090) at a density of 20 cells/well (6- and/or 3 day-clones), 200, 350 and 500 cells/well (6-day high-density clonal populations) and 500, 2500, 5,000, 10,000 or 50,000 cells/well (24 h mixed populations), depending on the experiment, in 50% conditioned medium and incubated for the corresponding time. Conditioned medium was prepared by collecting the medium from routinely passaged iBMDM cells, filtering through a 0.22 µm filter, and mixing (1:1) with fresh complete medium. Cells were seeded on the same plate on staggered days to ensure that all conditions were assayed simultaneously. For experiments containing exclusively mixed populations, 10,000 and 50,000 cells/well were seeded in complete medium in flat bottom 96-well TC treated imaging plates and incubated for 24 hours. For lipid A stimulation, cells were treated with lipid A at a final concentration of 500 ng/ml for 3 h, while untreated controls received an equivalent volume of 50% DMSO. Stimulation was performed in the presence of 10 µg/ml brefeldin A (Thermo Fisher, B7450) for intracellular TNFα detection. For infection assays, cells were infected with *Lm*-GFP at an MOI of 5 in pre-warmed serum-free medium for 45 min, washed once, and then cultured for 24 hours in medium supplemented with 10% FBS and 10 µg/ml gentamicin sulfate (Sigma-Aldrich, 345814). CD36 was inhibited with 200 µM sulfosuccinimidyl oleate sodium (SSO, Abcam, ab145039) overnight prior to infection and re-administered after infection at the time of gentamicin addition.

### Immunofluorescence

For CD36, IL1β, NF-κB RelA, RelB and TNFα detection, cells were fixed with 4% formaldehyde (Sigma Aldrich, 252549) in PBS at room temperature (RT) for 15 min and subsequently permeabilized with 0.1% triton X-100 (Sigma Aldrich, X100) in PBS for 10 min. Cells were blocked in 5% BSA (Sigma Aldrich, A7906) in PBS for 1 hour. For RelA staining, an additional 1 hour blocking step with Mouse-on-Mouse IgG Blocking Solution (Invitrogen, R37621) was performed at RT. The cells were then incubated overnight with primary antibodies in 5% BSA in PBS at 4 °C. Subsequently, secondary antibodies were applied in 5% BSA in PBS for 1 hour at RT in the dark. Nuclear DNA was stained using DAPI (Sigma Aldrich, D9542). For F4/80 and TNFR2 detection, cells were fixed with ice-cold methanol (Sigma Aldrich, 34860) for 5 minutes at -20 °C and subsequently permeabilized with 0.1% triton X-100 in PBS for 15 min at RT. After blocking in 2% BSA in PBS for 1 hour, the cells were incubated with primary antibodies in 0.1% BSA in PBS overnight at 4 °C. Subsequently, secondary antibodies were applied in 0.1% BSA in PBS for 1 hour at RT in the dark. Nuclear DNA was stained using DAPI.

Antibody list:

Anti-CD36 from rat, (Bio-Rad, MCA2748, 1/400)

Anti-F4/80 from rat (Abcam, ab6640, 1/200)

Anti- IL1β from goat (R&D Biosystems, AF-401-NA, 1/250)

Anti-NF-κB RelA/p65 from mouse (Cell Signaling Technology, 6956S, 1/400)

Anti-RelB from rabbit (Cell Signaling Technology, 10544S, 1/200)

Anti-TNFα from rabbit (Abcam, ab215188, 1/500)

Anti-TNFR2 from rabbit (Thermo Fisher, MA5-32618, 1/200)

Donkey anti-goat, Alexa Fluor 488 (Thermo Fisher, A32814, 1/500),

Donkey anti-mouse, Alexa Fluor 488 (Thermo Fisher, A21202, 1/1000)

Donkey anti-rabbit, Alexa Fluor 488 (Thermo Fisher, A21206, 1/1000)

Donkey anti-rabbit, Alexa Fluor 594 (Thermo Fisher, A21207, 1/500)

Donkey anti-rabbit, Alexa Fluor 647 (Thermo Fisher, A32795, 1/1000)

Donkey anti-rat, Alexa Fluor 594 (Thermo Fisher, A21209, 1/1000)

### High-content microscopy and image processing

Image acquisition was performed on the high-throughput Operetta CLS microplate microscope (PerkinElmer) using Harmony software v. 4.9. Tile arrays of images of individual wells were acquired with a 20x air objective (numerical aperture 0.4, part number HH14000404) for lipid A stimulation experiments (Figs. 3–6) and 40x water immersion objective (numerical aperture 1.1, part number HH14000422) for *Lm* infection (Fig. 7). The resulting raw images were exported, flat-field corrected and stitched using Maestro (https://github.com/kochanczyk/maestro). Subsequently, ShuttleTracker (https://pmbm.ippt.pan.pl/web/ShuttleTracker) was used for segmentation based on nuclear staining and fluorescent signal quantification [74]. Nuclear detection parameters were manually tuned on a subset of wells to identify the most suitable settings, after which these parameters were used for automated detection across the remaining wells for each set of replicates. The erroneous contours (e.g., corresponding to mitotic, overlapping, misshapen, or oversized cells) were manually removed. Perinuclear annular extensions were then generated for each nucleus, with manual fine-tuning performed on the first few wells to ensure optimal settings before being applied to the remaining wells for each set of replicates (see Table S6 for the utilised parameter sets for nuclear and perinuclear segmentation). Finally, regions corresponding to individual clones were manually delineated as separate objects. Nuclear and perinuclear fluorescence intensities were extracted using default ShuttleTracker settings and exported for analysis. Following quantification, segmented objects were analysed using Jupyter Notebooks in Python (v3.11). Data were additionally filtered based on cell morphological characteristics, by removing nuclei that have inappropriate size (smaller than 25 μm^2^ and larger than 225 μm^2^), based on published size distributions [10]. Nuclei that were not round (eccentricity score > 0.3, circularity normalised score < 0.145, longest horizontal vs vertical stretch > 12 pixels), or located too close to the edge of the well (<100 pixels) were also removed from the analysis. The average perinuclear (and in selected cases nuclear) fluorescent intensity per object, calculated from the bottom 80th percentile of the corresponding pixel-by-pixel fluorescence distribution, was used for fluorescence intensity quantification in all subsequent analyses.

### Quantification of imaging data

#### Fraction of responding cells

For IL1β, the fraction of “responding/positive cells” was determined as relative to the fluorescent intensity distribution of untreated clonal cells. Responding/positive cells were defined based on the single-cell fluorescent intensity threshold corresponding to the 97^th^ percentile of the overall single cell distribution of all clonal populations per plate (pooled across wells). For TLR-independent CD36, the fraction of CD36 positive and negative clones was first manually assessed. The 97^th^ percentile threshold (per plate) was then adjusted to reproduce obtained proportions of CD36-positive and -negative clones, assuming that a positive clone is characterised by more than 10% of responding cells. For each plate, the corresponding threshold was applied to classify cells as positive or negative across all clonal and mixed populations. The resulting data was used to calculate the fraction of responding/positive cells per clone, and the average fraction of responding/positive cells per well per plate for mixed population (e.g., for different seeding densities).

#### Estimation of local cell density

Local cell density was estimated based on the XY coordinates of the centroid of each segmented nuclear contour by determining the number of neighbouring cells within a radius of 50 pixels (corresponding to approximately 30 µm, given the pixel size of 0.598 µm for the 20x objective). Ensemble averages of individual cell densities were reported for clonal populations. For mixed populations ensemble averages were additionally averaged per well for each replicate-plate.

#### Daughter cell analysis

Daughter cells were identified within mixed populations seeded at 500 cells/well by scanning Euclidean distances between centroids of individual nuclei contours. A pair of daughter cells was defined as two nuclei located within a maximum distance of 30 pixels from each other, and at least 120 pixels away from the nearest third cell (pixel size of 0.598 µm). Wells with less than 50 daughter pairs were not included in the analysis. For odds ratio analysis, each daughter-cell was annotated as responding (‘ON’) and non-responding (‘OFF’), according to the fluorescent intensity thresholds previously established for calculating the fraction of responding cells in the corresponding plate. Odds ratios were calculated as 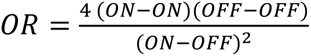 for all the daughter pairs per plate [18].

#### Generation of virtual clones

For each plate, we matched the number and sizes of experimentally observed clones and generated “virtual” clones by randomly sampling, with replacement, single-cell protein intensities from control mixed populations (seeded at 10,000 cells per well), under either treated or untreated conditions. For the plate-level statistics (i.e., difference in the CV between the observed and virtual clones within each clone-size bin), one-sided *p*-values were estimated by bootstrap resampling (n=2,000 iterations). To account for the finite solution of the bootstrap procedure and avoid zero *p*-values, we applied a standard correction 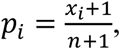 where *x_i_* is the number of bootstrap samples exceeding experimental CV on plate *i*. Overall *p*-values across plates were then computed using Fisher’s method [138]. Specifically, plate-based *p*-values (*p_i_*) for each gene and treatment were combined as 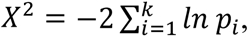 and the overall combined *p*-value was calculated as 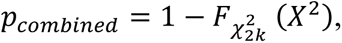 where 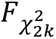 is the cumulative distribution function of the chi-square distribution with *2k* degrees of freedom, *k* is the number of plate-level *p*-values included in the combination.

#### Estimation of variability across clone sizes

CV was used to quantify inter-clone variability for each gene based on the average signal intensity per clone. Data were binned according to the clone size, expressed as log₂(N), with the following ranges: log₂N ≤ 4, 4 < log₂N ≤ 6, 6 < log₂N ≤ 8, log₂N > 8. Data was represented as the mean and SD of biological replicates, per treatment. To calculate the CV of the fraction of responding cells, a bin width of 1.0 was applied, and only bins containing at least five clones were included. Uncertainty in CV estimates was assessed by bootstrap resampling (n = 300 iterations) with replacement from the clones within each bin. The 95% confidence intervals were defined by the 2.5^th^ and 97.5^th^ percentiles of the bootstrap distribution. To visualize trends in variability across clone sizes, the binned CV values were fitted using locally weighted scatterplot smoothing (LOESS) with a smoothing fraction of 0.35. For each gene, a smoothed curve was generated over 250 evenly spaced points (Δlog₂N=1), together with bootstrapped 95% confidence intervals.

#### Regression analysis of the fraction of responding cells

Multiple linear regression using Ordinary Least Squares (OLS) method was performed to model the fraction of CD36 and IL1β responding/positive cells per clone as function of cell size (*log_2_N*) and local density (*Density*). To account for potential non-additive effects, a two way-interaction term (*log_2_N* × *Density*) was included in the model. To control for experimental variability, *Plate* was included as a categorical (batch) factor:

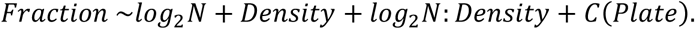

To test whether the interaction term significantly improved model fit, we compared the full model with a reduced model (excluding the interaction term) using a partial F-test, which determined whether the increase in R^2^ was statistically significant. Sequential (Type I) ANOVA was used to partition explained variance and quantify the unique contribution of each predictor to the total model R^2^. To assess potential multicollinearity among predictor variables, Variance Inflation Factor *VIF=1/(1-R^2^)* was calculated for the cell size and density in the regression models, where *R^2^*is the coefficient of determination for the predictor variables. VIF values below 2 indicated negligible multicollinearity in all models.

#### Logistic regression of single-cell responsiveness

To assess how cell morphology, clone size and local density influence responsiveness, logistic regression models were fit separately for IL1β and CD36 expression. Each observation represented a single cell, with response status coded as binary (off = 0, on = 1). Morphological and density features were screened for collinearity; *circularity* and *circularity_normalized* were excluded. The final predictors included *nuclear area*, *nuclear eccentricity*, *perinuclear area, solidity, longest_hv_stretch*, *Density*, *log_2_N*, all standardized (Z-scored) before fitting. Separate generalized linear models (GLMs) with a binomial error distribution and logit link function were fitted for IL1β and CD36 to model the odds of a cell being a responder:

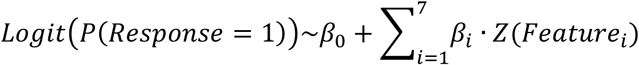

Model coefficients (β) were reported as the change in log-odds of a cell being classified as *responding* (“on”) for a one standard deviation (SD) increase in the predictor variable. Odds ratios OR = exp(β) were calculated to represent the multiplicative change in response odds per SD increase in each predictor. Model performance was evaluated using the Cox and Snell pseudo-R² statistic, quantifying the proportion of variance explained by each model and allowing comparison of explanatory power between markers (Table S8). Models were implemented in Python using the statsmodels library, utilizing the Generalized Linear Model (GLM) module (statsmodels.formula.api.glm).

#### Spatial correlation analysis

Spatial relationships between molecular markers were quantified using Bivariate Moran’s I. To define the local neighbourhood for each cell, a k-nearest-neighbour (kNN) method was employed. For the spatial cross-correlation between IL1β intensity and CD36 expression (Fig. 6), the k value was adjusted based on cell density: k = 100 was applied for high-density clonal (200, 350, and 500 cells per well) and mixed populations, while k = 15 was used for low-density clones (20 cells per well) (Fig. S14D). To eliminate background fluorescence, the analysis was restricted to cells exceeding a 30 a.u. intensity threshold for the average perinuclear IL1β expression. For the correlation between *Lm*-GFP intensity and CD36 expression (Fig. 7), a value of k = 100 was used (Fig. 7E). Additionally, Pearson’s r was calculated specifically for neighbourhoods exhibiting a positive *Lm*-GFP signal (Fig. 7D).

### Analysis of *Lm* infection in clonal populations

After visually selecting the most relevant areas of each well containing clonal populations, each area was scanned with the 40x water immersion objective on successive scans. Clonal infections were then analysed using ShuttleTracker to manually delineate the regions of each clone. The average CD36 expression (high or low) of each clone, as well as the presence of infection foci, was manually recorded and counted.

### Mathematical modelling of clonal expansion

Clonal IL1β and CD36 expression were modelled using a stochastic two-state switching framework in which cells transition between ON and OFF states with rates *k_ON_* and *k_OFF_*, respectively, while proliferating at rate λ (Fig. 5F). Switching rates were inferred based on daughter-cell pair data: the fraction of cells in the ON state (*f_ON_*), and OR between paired cells. Specifically, *f_ON_ = k_ON_/(k_ON_+k_OFF_)*, and *OR = 4ab/c^2^ = (k_ON_+λ)(k_OFF_+λ)/(k_ON_k_OFF_)*, where *a*, *b* and *c* denote the number of ON-ON, OFF-OFF, and mixed-state pairs, respectively [48]. Three model structures were considered: (1) constant-rate model; (2) population-size-dependent model, in which *k_ON_* is defined as a Hill-type function, *k_ON_ = k_ON_^min^ +(k_ON_^max^-k_ON_^min^)N^n^/(K^n^+ N^n^),* where *k ^min^* and *k ^max^* denote the minimal and maximal switching rates, *K* is the population size at half-maximal activation, and *n* is the Hill coefficient; and (3) extrinsic-noise model, in which *k_ON_^max^* is modelled as a Beta-distributed random variable across clones. Parameters were estimated using a two-state bootstrapping procedure to obtain confidence intervals (1000 resamples per bin). For Models 2 and 3, stochastic simulations (Gillespie algorithm), combined with nearest-neighbour matching to experimental clone sizes, were used to compare model outputs with data using a weighted least square objective function (refer to Supplementary Methods for detailed description of methodology and results).

### Statistical Analyses

Statistical analysis was performed using GraphPad Prism 10 software, unless otherwise stated. Data was tested for normality using D’Agostino-Pearson test.

## Data Availability

Generated sequencing data have been deposited in the ArrayExpress database at EMBL-EBI under accession number E-MTAB-11041 (https://www.ebi.ac.uk/biostudies/arrayexpress/studies/E-MTAB-11041) and E-MTAB-13014 (https://www.ebi.ac.uk/biostudies/arrayexpress/studies/E-MTAB-13014). Generated images and image processing data have been deposited in BioImageArchive under accession number S-BIAD2517 (DOI <u>10.6019/S-BIAD2517</u>). Custom scripts used for data analysis and mathematical modelling are available from GitHub repository (https://github.com/cellbiologyppaszek/macrophage-heritability).

## Conflict of Interest

*The authors declare that the research was conducted in the absence of any commercial or financial relationships that could be construed as a potential conflict of interest*.

## Funding

This work was supported by Polish National Agency for Academic Exchange (BPN/PPO/2022/1/00002), National Science Centre Poland (2022/45/B/NZ6/01643) and Biotechnology and Biological Sciences Research Council UK (BB/R007691/1). NA was supported by Wellcome Trust PhD Studentship, A. Srivastav was supported by NIH Award Number T32GM142603. We also thank The Polish high-performance computing infrastructure PLGrid (HPC Centers: ACK Cyfronet AGH, CI TASK) for providing computational facilities and support within computational Grant No. PLG/2025/018818.

## Author contributions

JAR performed microscopy experiments, TCN performed quantification of imaging data, JM conducted genomics experiments, NA performed analyses of genomics data. A. Srivastav and A. Singh performed mathematical modelling. MK supported image quantification. MM, IR, AKB, AS and PP provided supervision and conceptualisation. PP with assistance of JAR, NA and TCN prepared displays and wrote the manuscript. All authors read and approved the final manuscript.

## Acknowledgements

We thank Ian Donaldson, I-Hsuan Lin, Claire Morrisroe and Andy Hayes of the Bioinformatics and Genomic Technologies Core Facilities at the University of Manchester for providing support regarding genomics analyses. Schematic diagrams were created with BioRender.

## Supplementary Tables

**Table S1.** Coefficient of variation fold change of MemorySeq genes.

**Table S2**. Ontology analysis of heritable TLR-independent MemorySeq genes.

**Table S3.** Coefficient of variation fold change of scRNA-seq genes.

**Table S4.** Expressed genes in both MemorySeq and scRNA-seq experiments.

**Table S5.** Heritable genes in both MemorySeq and scRNA-seq datasets.

**Table S6.** Parameter sets used for nuclear detection and perinuclear derivatization in ShuttleTracker.

**Table S7.** Estimation of the doubling time (T_d_) for the cells passaged under standard conditions.

**Table S8.** Regression analysis of IL1β and CD36 expression in clonal populations.

**Table S9.** NMF analysis of scRNA-seq data

## Notes

https://www.ebi.ac.uk/biostudies/arrayexpress/studies/E-MTAB-11041

https://www.ebi.ac.uk/biostudies/arrayexpress/studies/E-MTAB-13014

https://www.ebi.ac.uk/biostudies/bioimages/studies/S-BIAD2517

https://github.com/cellbiologyppaszek/macrophage-heritability

### Competing Interest Statement

The authors have declared no competing interest.

