## Supplementary figures, tables and methods for "Heritable single-cell gene expression states shape functional variability in innate immune responses"

### 1. Supplementary Figures

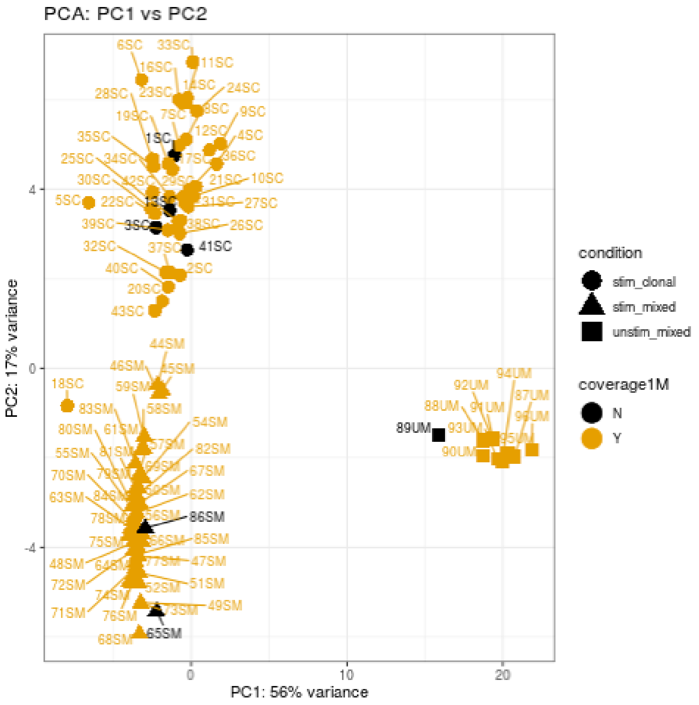

**Figure S1. Analysis of MemorySeq cell populations.** Principal component analysis (PCA) of stimulated clonal (SC, *stim_clonal*), stimulated mixed (SM, *stim_mixed*), and untreated mixed (UM, *unstim_mixed*) populations. The variance explained by each principal component is indicated on the corresponding axis. Samples shown in black had sequencing coverage between 0.5 and 1 million reads. Sample *18SC* was excluded from subsequent analyses.

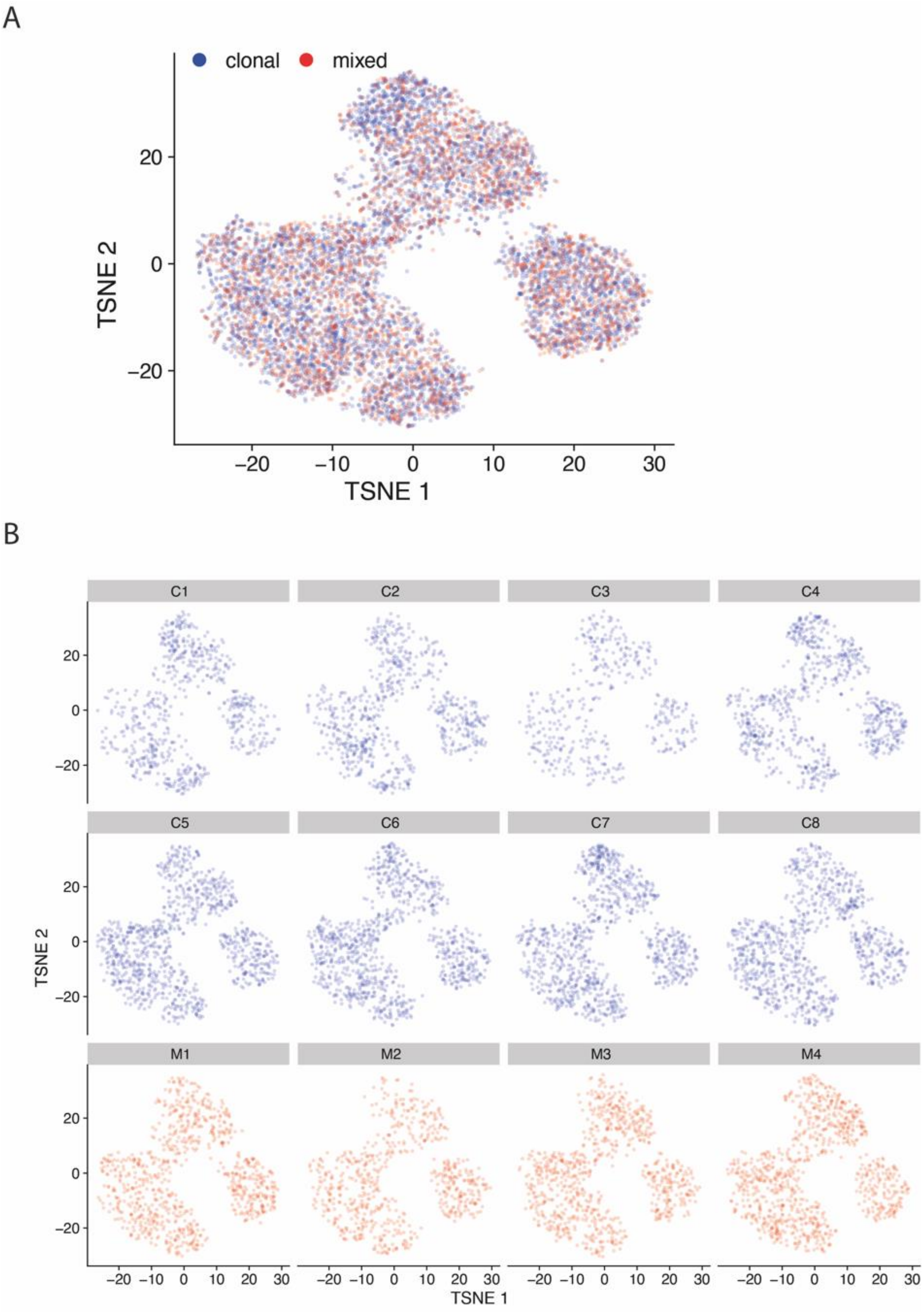

**Figure S2: Analysis of the scRNA-seq populations. A**. t-SNE plot for single-cell gene expression levels for clonal and mixed populations pooled together. **B**. t-SNE plots of individual clonal (C1 to C8) and mixed (M1 to M4) cells.

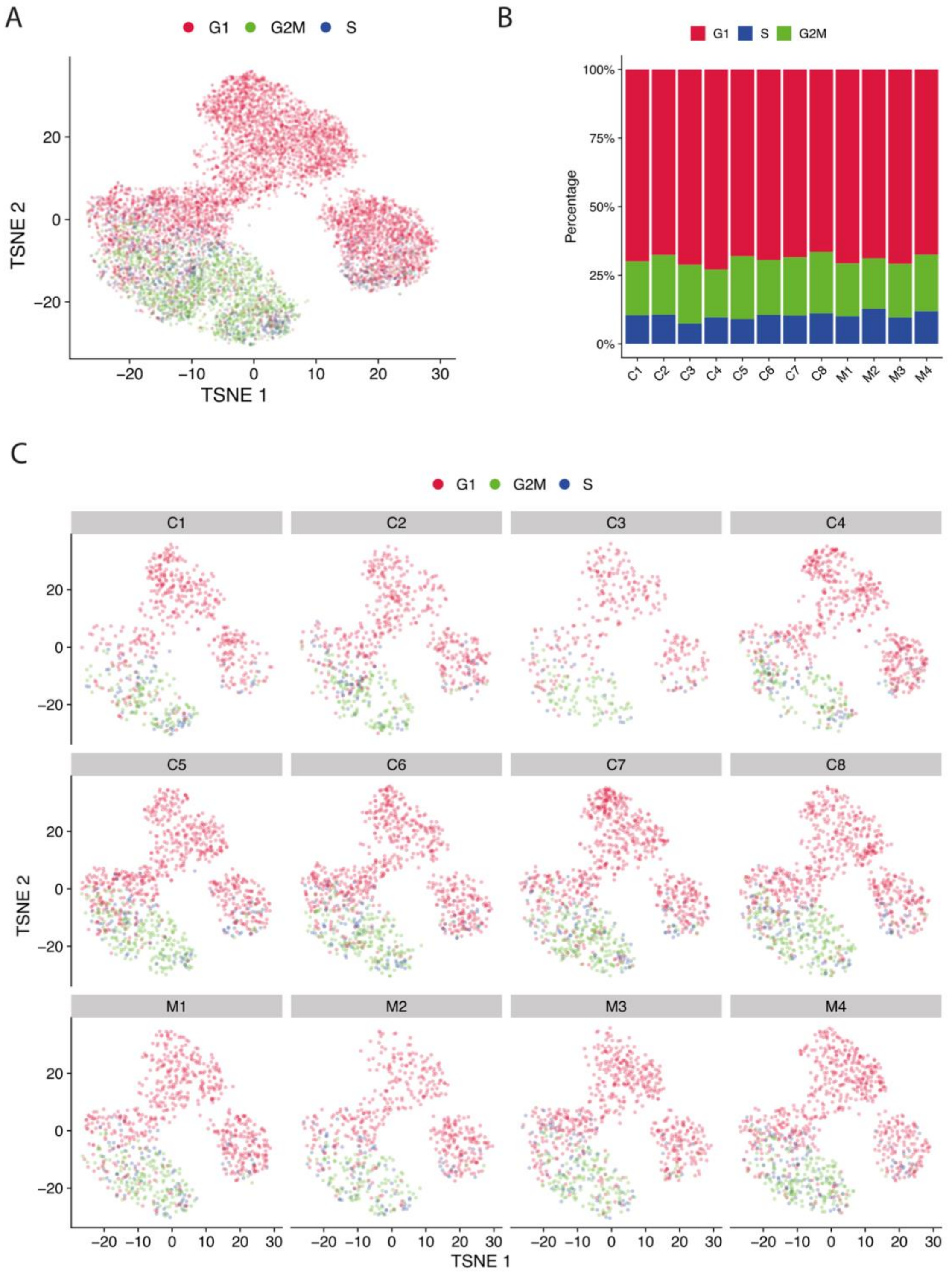

**Figure S3: Cell cycle analysis of the scRNA-seq populations. A**. t-SNE plot of single-cell gene expression levels pooled across clonal and mixed populations, with inferred cell-cycle stages indicated by colour. **B**. Distribution of cell cycle stages across individual clonal (C1 to C8) and mixed (M1 to M4) populations. **C**. t-SNE plots of individual clonal (C1–C8) and mixed (M1–M4) populations coloured by inferred cell-cycle stage.

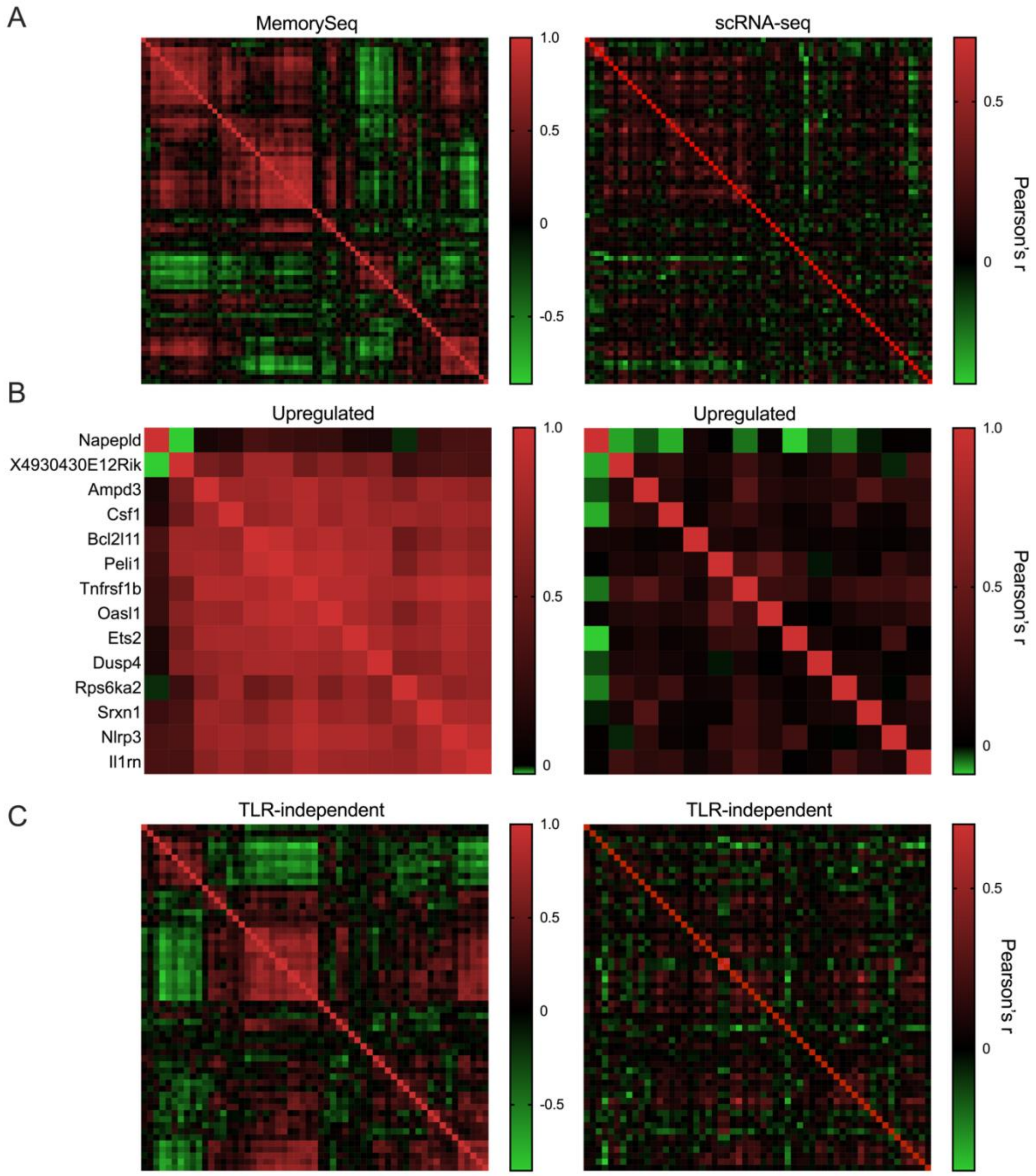

**Figure S4: Heritable genes exhibit co-variability.** Shown are Pearson's correlation heatmaps for all pair combinations of 73 heritable genes in the MemorySeq (left) and scRNA-seq (right) datasets. **A**. Correlation heatmap of all heritable genes. **B**. Correlation heatmaps of up-regulated heritable genes. **C**. Correlation heatmaps of TLR-independent heritable genes. Heatmaps for scRNA-seq data are displayed using the ordering of the MemorySeq correlation matrix.

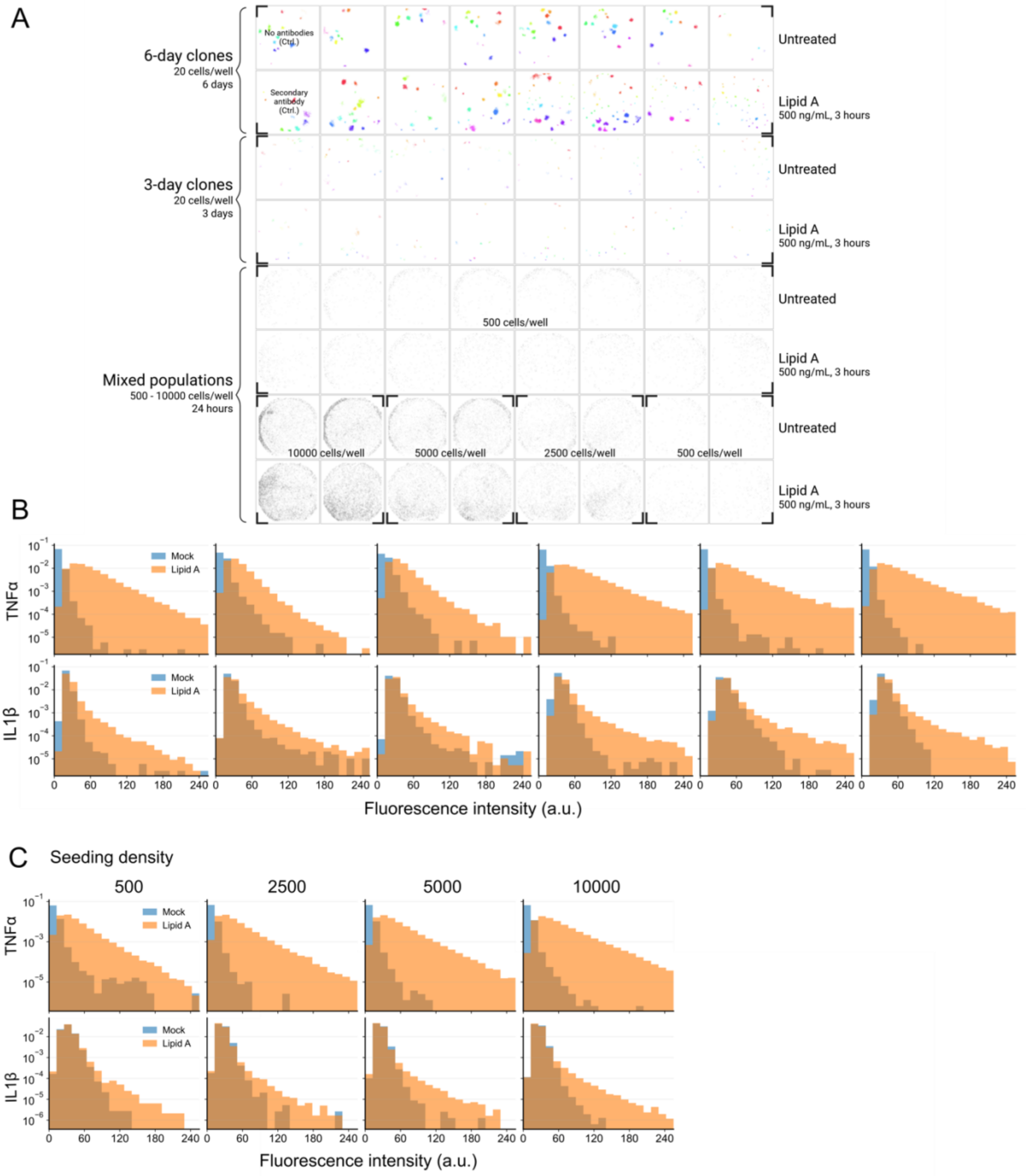

**Figure S5. TNFα and IL1β protein expression in clonal and mixed populations. A.** 96-well plate layout of the immunostaining experiment under different experimental conditions, representative of six biological replicates. Segmented individual clonal populations are highlighted in distinct colours; single non-clonal cells are shown in grey. **B**. Distributions of perinuclear protein expression in clonal populations across biological replicates. Cells stimulated with 500 ng/ml lipid A for 3 h (orange) and untreated cells (blue, Mock) are shown. Data pooled across day 3 and day 6 clones per plate. **C.** Distributions of perinuclear protein expression in mixed-population controls at different seeding densities (500, 2,500, 5,000, and 10,000 cells per well). Data pooled across six replicate experiments.

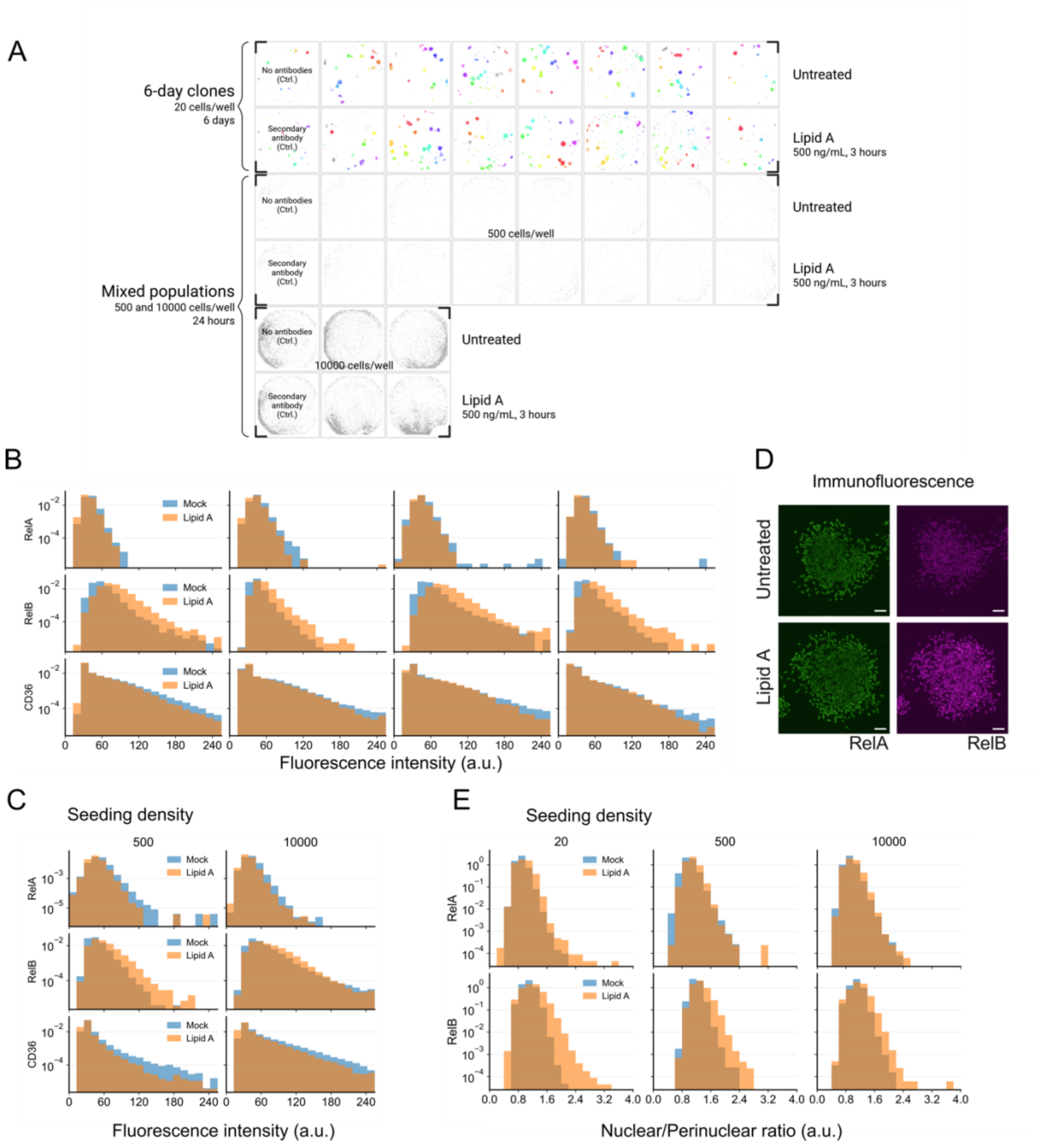

**Figure S6. RelA, RelB and CD36 protein expression in clonal and mixed populations. A.** 96-well plate layout of the immunostaining experiment, representative of four biological replicates. Segmented individual clonal populations are highlighted in distinct colours; single non-clonal cells are shown in grey. **B**. Distributions of perinuclear protein expression in clonal populations across biological replicates. Cells stimulated with 500 ng/ml of lipid A for 3 hours (orange) and untreated cells (blue, Mock) are shown. **C.** Distributions of perinuclear protein expression in mixed-population controls at seeding densities of 500 and 10,000 cells per well. Data pooled across replicate experiments (three for 10,000 and four for 500 seeding density). **D**. Representative microscopy images of untreated and lipid A-stimulated clonal populations for RelA and RelB. Scale bar, 50 µm. **E.** Distributions of nuclear-to-perinuclear RelA and RelB intensity ratios across populations seeded at different densities. Data pooled across replicate experiments (three for 10,000, four for 500 and clonal populations).

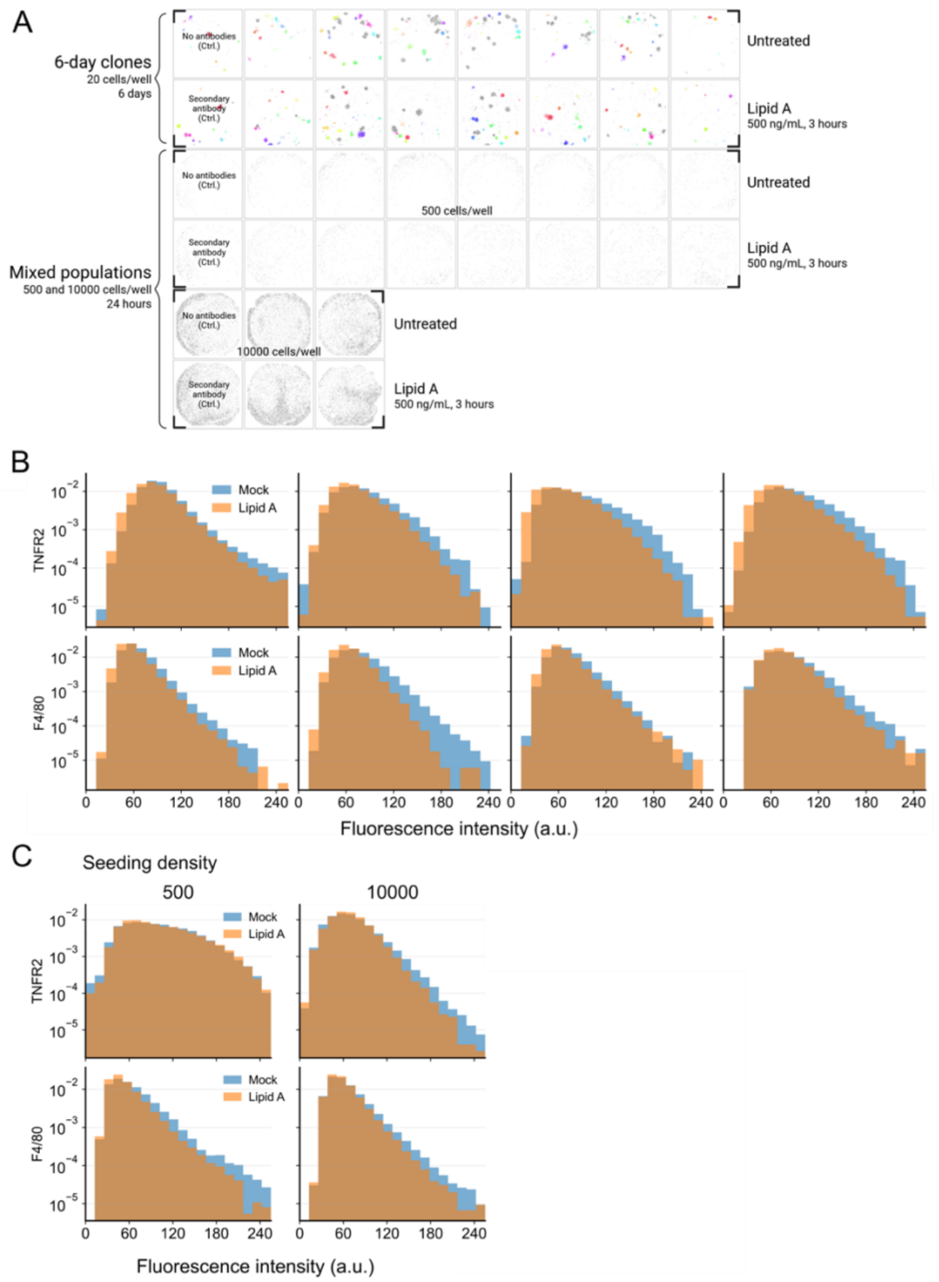

**Figure S7. TNFR2 and F4/80 protein expression in clonal and mixed populations. A.** 96-well plate layout of the immunostaining experiment, representative of four biological replicates. Segmented individual clonal populations are highlighted in distinct colours; single non-clonal cells are shown in grey. **B**. Distributions of perinuclear protein expression in clonal populations across biological replicates. Cells stimulated with 500 ng/ml lipid A for 3 h (orange) and untreated cells (blue, Mock) are shown. **C.** Distributions of perinuclear protein expression in mixed-population controls at seeding densities of 500 and 10,000 cells per well. Data pooled across replicate experiments (three for 10,000, four for 500 seeding density).

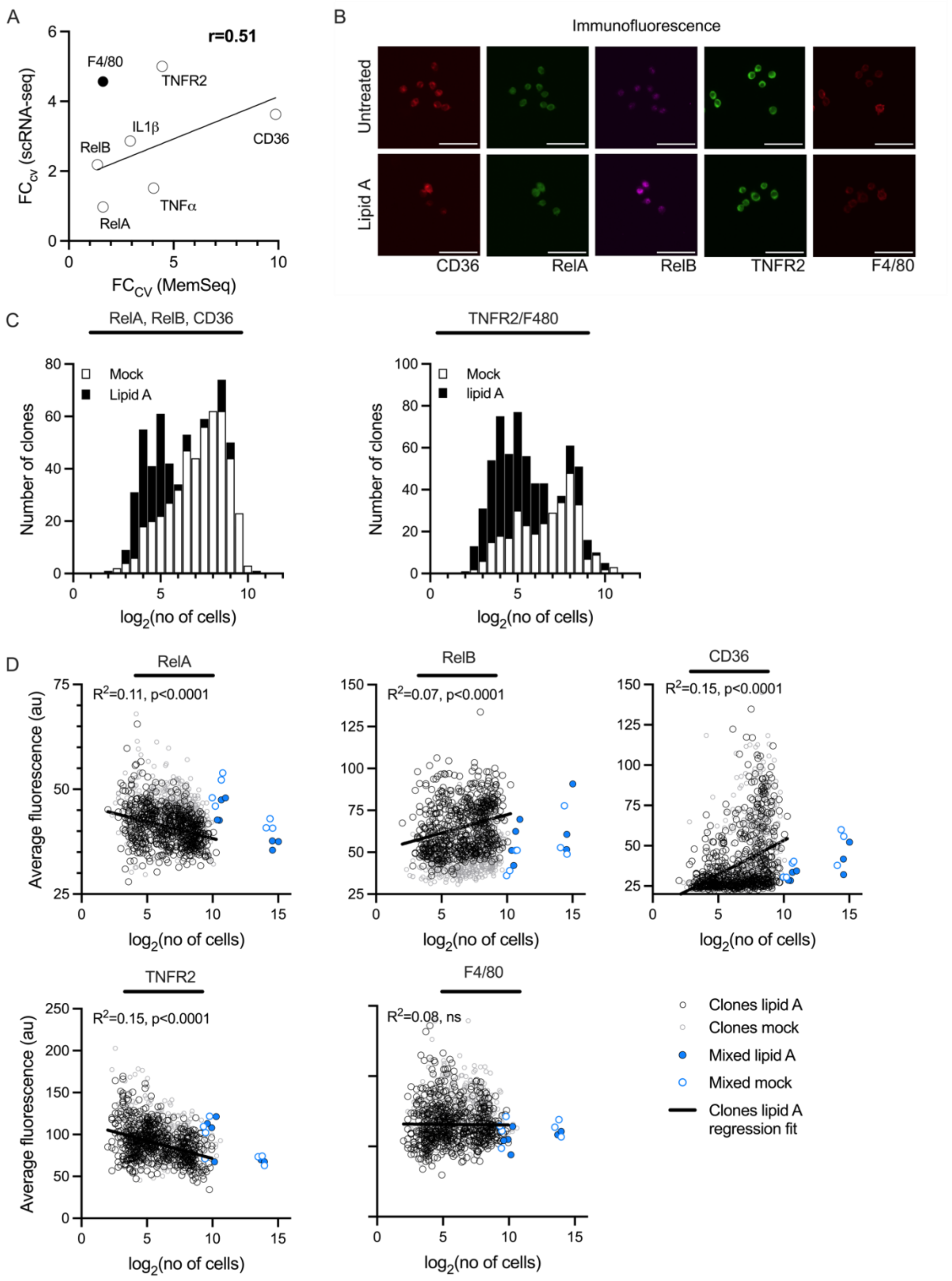

**Figure S8. Heritability analysis of RelA, RelB, CD36, TNFR2, and F4/80 protein expression across clonal populations. A.** Correlation between the heritability index ($FC_{CV}$) from MemorySeq and scRNA-seq data (lipid A-stimulated populations) for the genes selected for immunostaining. Circles represent individual labelled genes. Linear regression fit indicated in black. Pearson's correlation coefficient (*r*) calculated after excluding F4/80. **B**. Representative images of mixed-population controls (seeded at 500 cells per well) co-stained for RelA, RelB, and CD36, and separately co-stained for TNFR2 and F4/80. Scale bar, 50 μm. **C.** Histograms of clonal population size distributions. Shown are $log_2$-transformed clone sizes for untreated (Mock) and lipid A-stimulated conditions. From left to right: populations corresponding to RelA, RelB, and CD36; and TNFR2 and F4/80 immunofluorescence measurements (see Fig. 4). **D**. Relationship between average perinuclear signal intensity per clone and clone size ($log_2$-transformed number of cells per clone) for data from C. Black circles indicate individual lipid A-stimulated clonal populations; grey circles represent untreated (Mock) clonal populations. Filled blue circles show average expression in lipid A-stimulated mixed populations across a range of seeding densities (average signal per well versus total cell number per well across replicate plates); open blue circles correspond to untreated mixed populations. The black line represents the linear regression fit for lipid A-stimulated clonal populations; the coefficient of determination ($R^2$) and *p*-value for the non-zero slope are indicated on the plot.

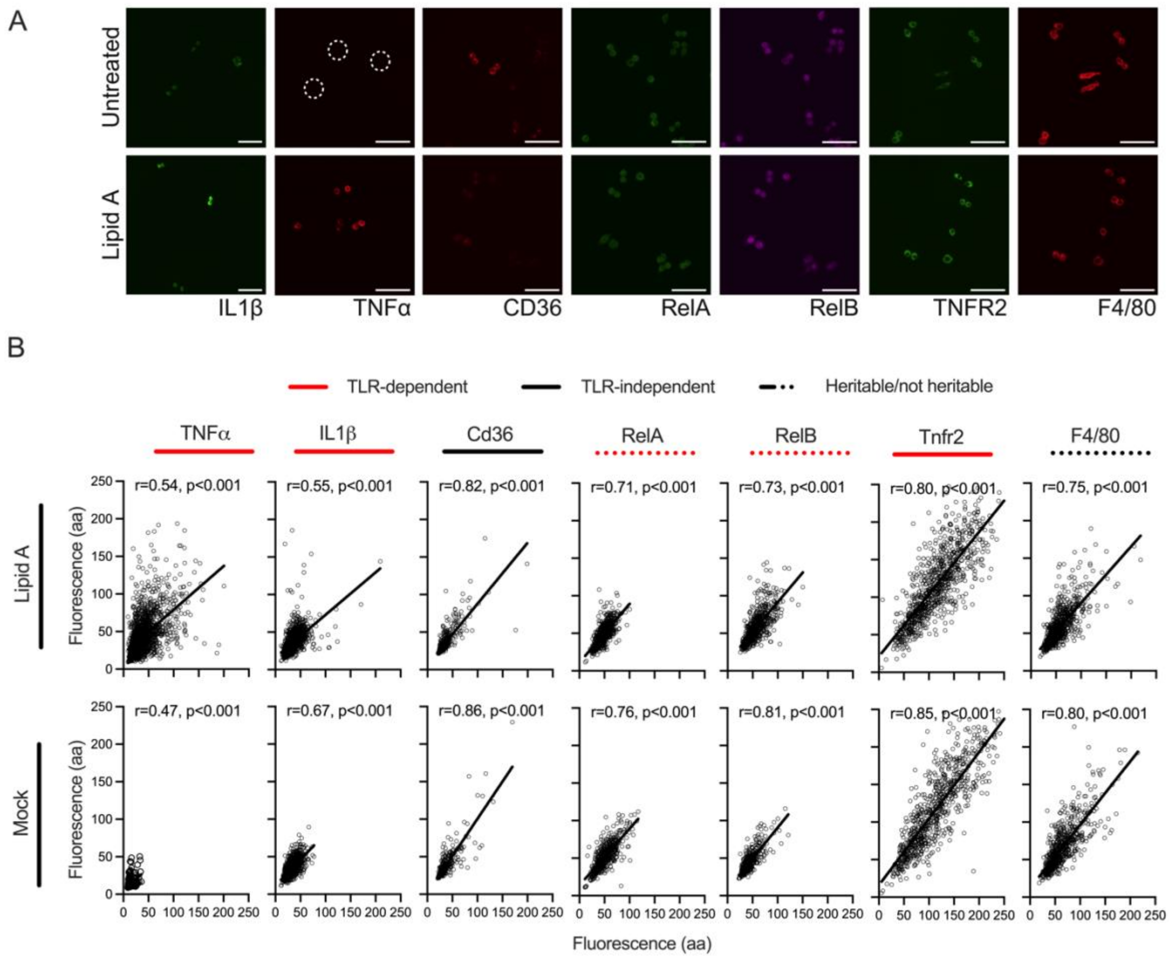

**Figure S9. Correlation between daughter cells**. **A**. Representative images of daughter-cell pairs from mixed populations, seeded at 500 cells per well, co-stained for IL1β and TNFα; RelA, RelB, and CD36, as well as separately co-stained for TNFR2 and F4/80. Scale bar, 50 µm. Dashed circles indicate cells with low TNFα expression under untreated conditions. **B**. Correlation between daughter-cell pairs for protein targets shown in A. Each circle represents perinuclear fluorescence levels in one daughter-cell pair; solid lines show linear-regression fits. Pearson's correlation coefficients (r) and corresponding p-values are indicated in each panel. Daughter-cell pairs were extracted from mixed-cell populations seeded at 500 cells per well based on cell-to-cell distances (see methods). From left to right: TNFα and IL1β (1750 and 1392 pairs, for lipid A and untreated conditions, respectively, from Fig. S5), RelA, RelB, and CD36 (771 and 566 pairs, for lipid A and untreated conditions, respectively, from Fig. S6); and TNFR2 and F4/80 immunofluorescence (889 and 749 pairs, for lipid A and untreated conditions, respectively, from Fig. S7). Heritable genes highlighted with a solid line, non-heritable genes with a broken line; TLR-dependent genes are shown in red, and TLR-independent genes in black as in Fig. 4B.

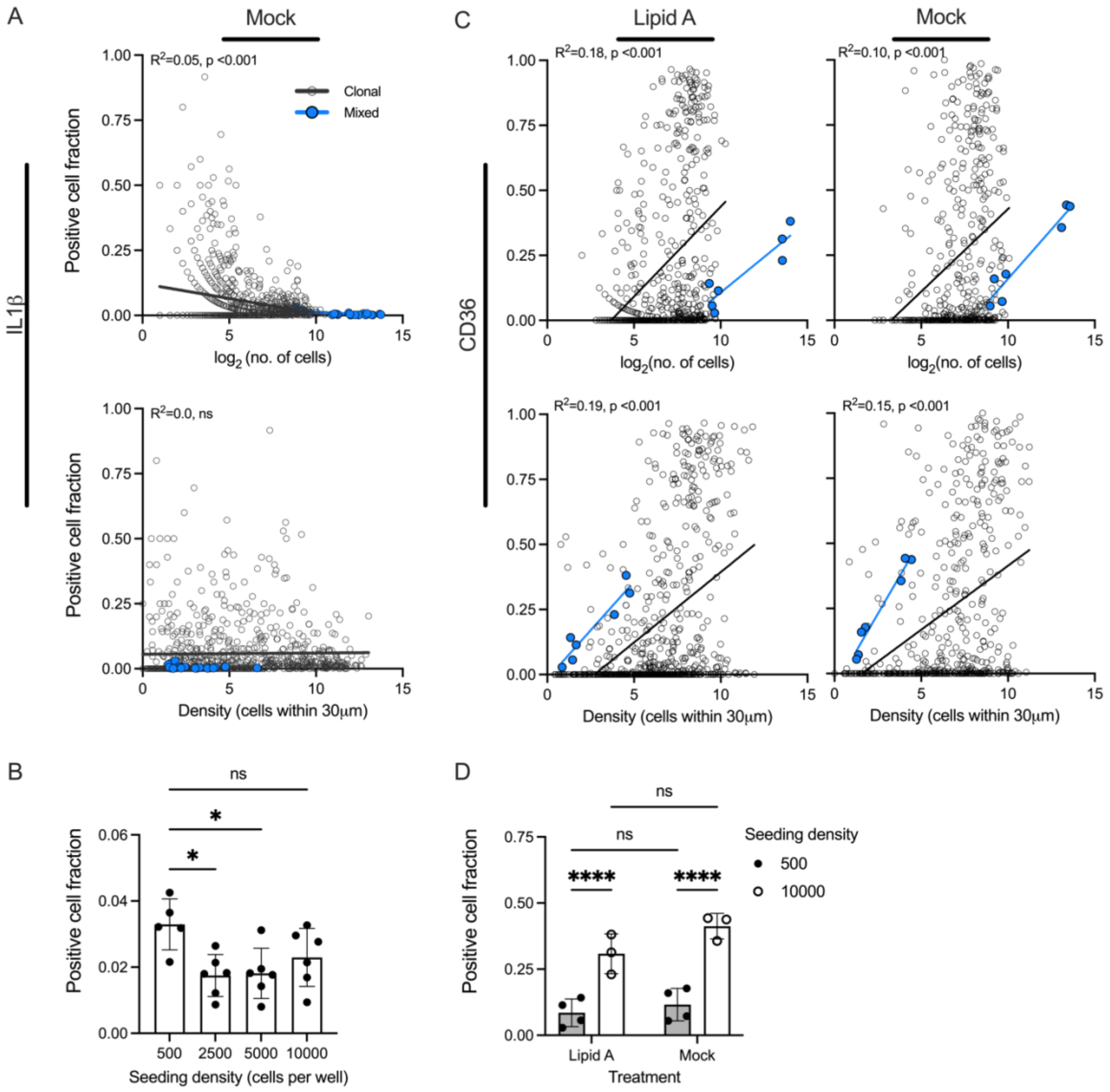

**Figure S10. Fraction of IL1β responding and CD36 expressing cells as function of clone size (*N*) and cell density. A**. Relationship between the fraction of IL1β-positive cells per clone and clone size (*N*, $\log_2$-transformed, top) and cell density (expressed as an average number of cells within 30 μm radius per population, bottom) calculated for untreated (Mock) clonal and mixed populations. Black circles represent individual clonal populations; blue circles denote mixed populations across a range of seeding densities. Solid lines indicate linear regression fits; the coefficient of determination ($R^2$) and *p*-value for the non-zero slope are shown for clonal populations. Data pooled across six biological replicates (see Fig. 3). **B**. Fraction of lipid A-stimulated IL1β-positive cells in mixed populations as function of seeding density. Shown are fractions of responding cells averaged per plate represented with circles, with mean ± SD shown across six biological replicates/plates (see Fig. 3). Statistical significance was determined using the Kruskal-Wallis test with multiple comparisons ($p < 0.05$; ns, non-significant). **C**. Relationship between the fraction of CD36 positive cells per clone and clone size (*N*, $\log_2$-transformed, top) and cell density (expressed as an average number of cells within 30 μm radius per well, bottom) calculated for clonal and control mixed populations (formatted as in A). Shown is the data for lipid A-stimulated and untreated populations (see Fig. 4). **D.** Fraction of lipid A-stimulated (Lipid A) and untreated (Mock) CD36-positive cells in mixed population controls as function of seeding densities, as indicated. Circles represent average fractions per well, with mean ± SD across biological replicates/plates (see Fig. 4). Statistical significance was determined using two-way ANOVA with multiple comparisons (****$p$ < 0.001; ns, non-significant).

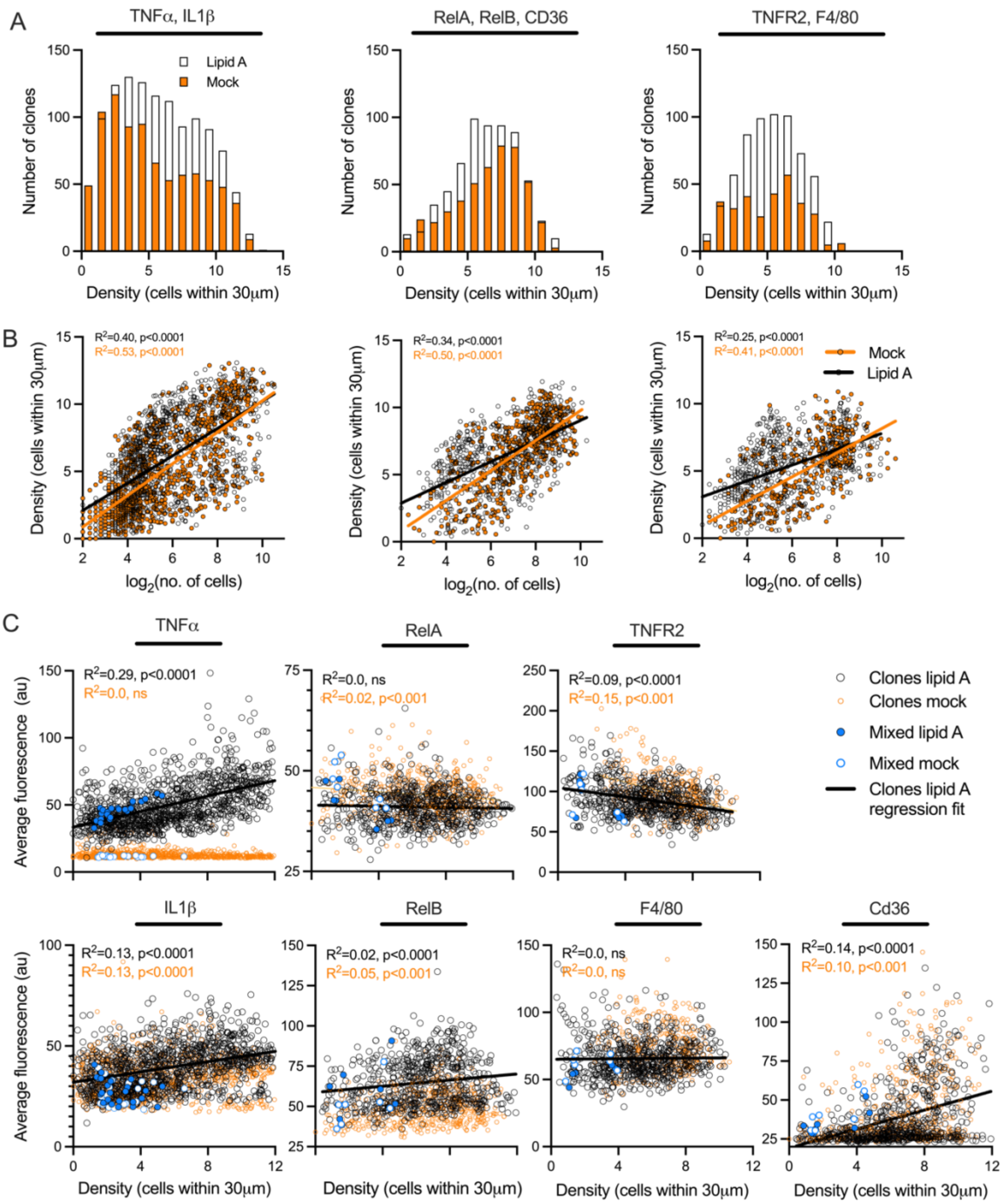

**Figure S11. Effect of cell density on protein expression in clonal populations. A.** Histograms showing the distribution of cell density in lipid A-stimulated (white) and untreated (orange) clonal populations. From left to right: populations corresponding to IL1β and TNFα (see Fig. 3); RelA, RelB, and CD36; and TNFR2 and F4/80 immunofluorescence measurements (see Fig. 4). **B**. Correlation between cell density (expressed as an average number of cells within 30 μm radius per population) and clone size across clonal populations. Scatter plots show cell density versus clone size ($log_2$-transformed number of cells) for the populations in A. Black circles indicate individual lipid A-stimulated clonal populations; orange circles represent untreated (Mock) populations. Solid lines indicate linear regression fits for lipid A-stimulated (black) and untreated (orange) populations. The coefficient of determination ($R^2$) and *p*-value for the non-zero slope are shown on each plot. **C.** Correlation between average perinuclear protein expression and cell density for clonal and mixed populations. Scatter plots show the relationship between average expression and cell density (expressed as an average number of cells within 30 μm radius per population) for the populations in A. Black circles indicate lipid A-stimulated clonal populations; orange circles represent untreated (Mock) populations. Filled blue circles show average expression in lipid A-stimulated mixed populations across a range of seeding densities; open blue circles correspond to untreated mixed populations (averaged per well). Solid lines indicate linear regression fits for lipid A-stimulated populations. The coefficient of determination ($R^2$) and *p*-value for the non-zero slope are indicated on each plot.

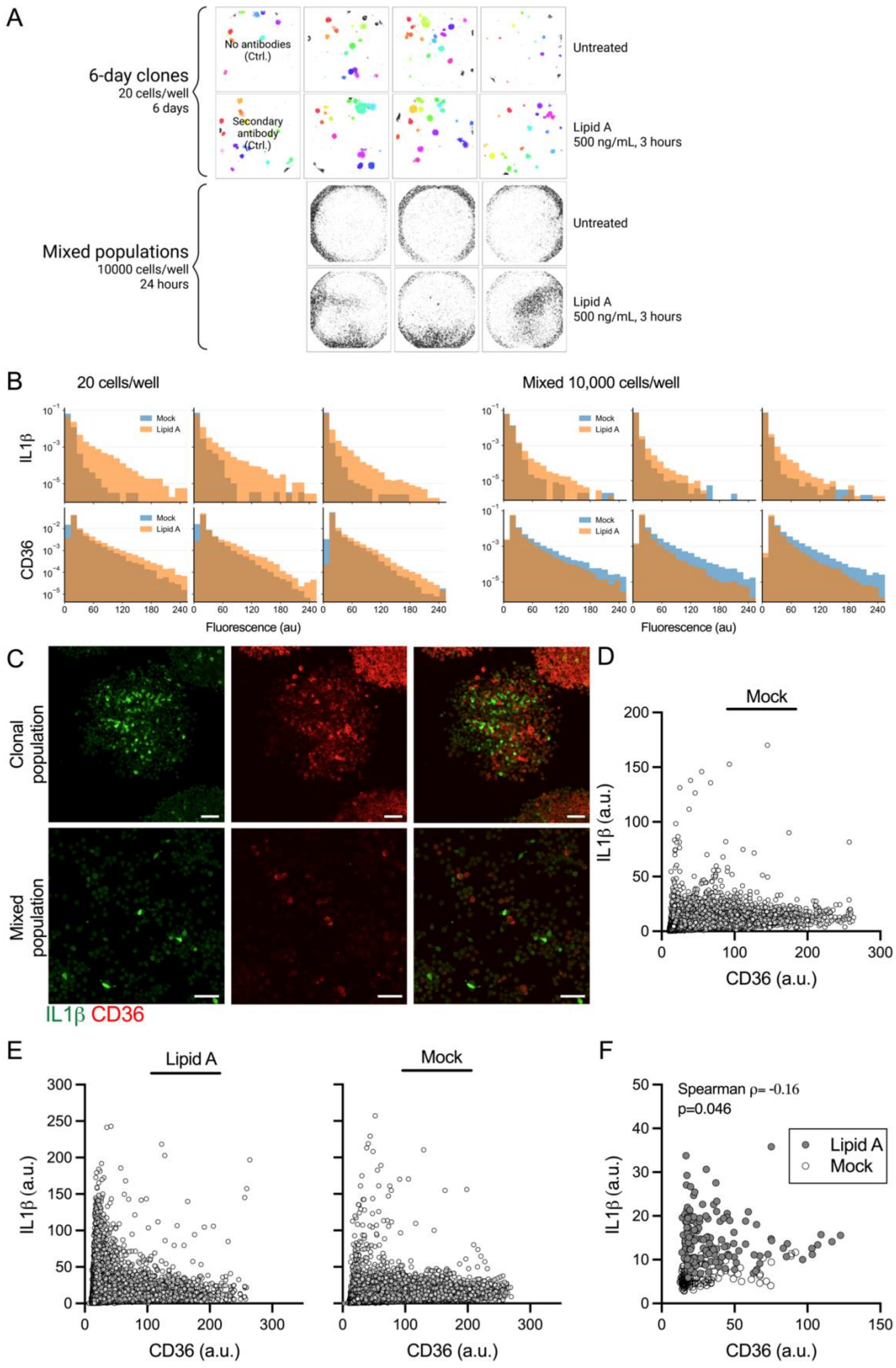

**Figure S12. CD36 and IL1β protein expression in clonal and mixed populations. A.** 96-well plate layout of the immunostaining experiment, representative of three biological replicates. Segmented individual clonal populations are highlighted in distinct colours; single non-clonal cells are shown in grey. **B**. Distributions of perinuclear protein expression in populations from A across biological replicates. Cells stimulated with 500 ng/ml lipid A for 3 h (orange) and untreated cells (blue, Mock) are shown. **C**. Representative composite images of clonal and mixed populations co-stained for IL1β (green) and CD36 (red) following lipid A treatment. Top: Clonal populations seeded at 20 cells per well; bottom: mixed populations seeded at 10,000 cells per well. Scale bar, 50 µm. **D.** Cell-level relationship between CD36 and IL1β protein expression in untreated (Mock) clonal populations. Shown are single cell fluorescent levels across three biological replicates. **E. C**ell-level relationship between CD36 and IL1β protein expression in mixed populations. Shown are single cell fluorescent levels across mixed populations either stimulated with lipid A (left) or untreated (right) across three biological replicates. **F**. Clone-level relationship between CD36 and IL1β protein expression in clonal populations. Shown is average fluorescence per clonal population for Mock (open circles) and lipid A-treated samples (grey circles). Correlation for lipid A-treated clonal populations assessed using Spearman's ρ (p-value indicated).

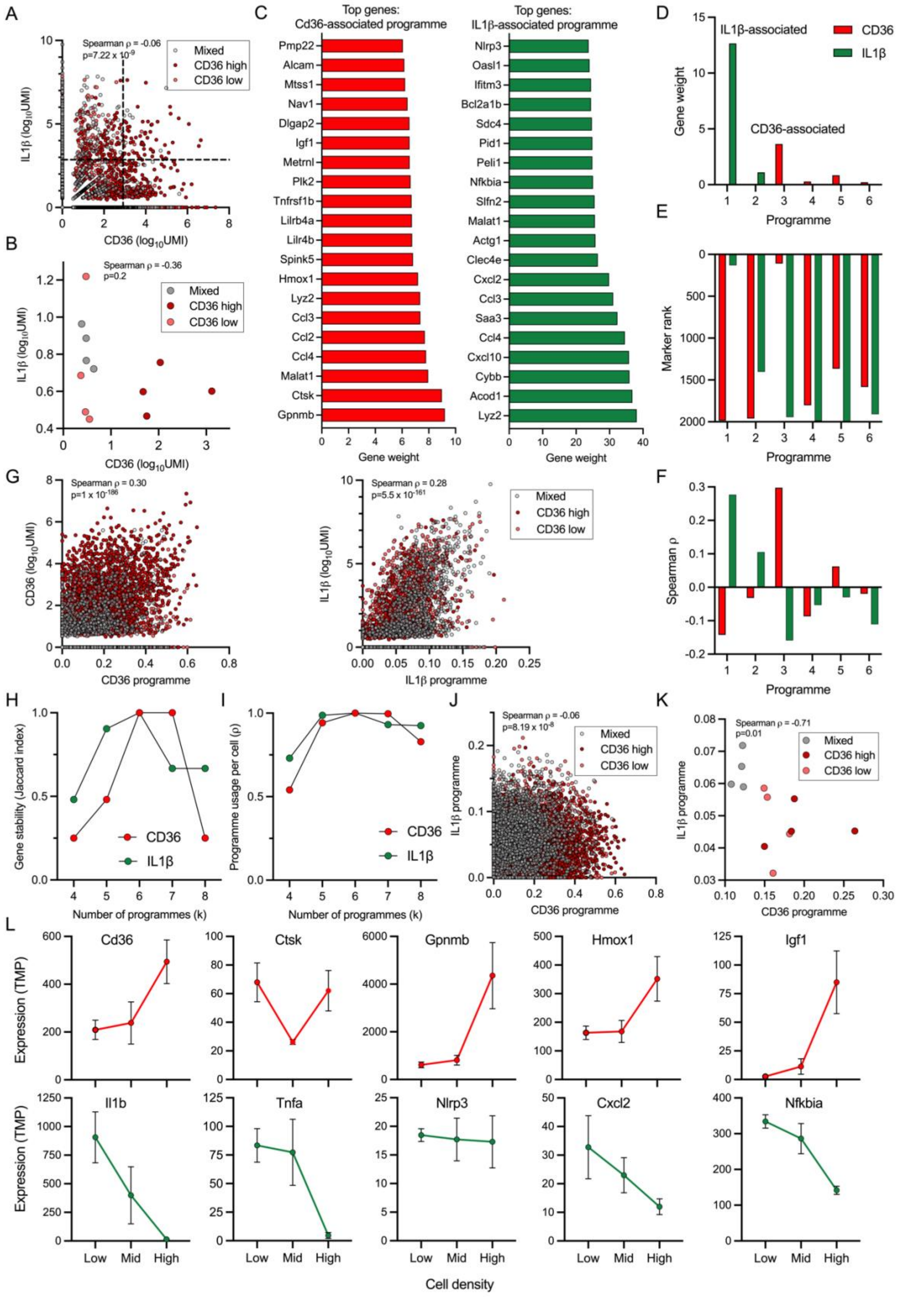

**Figure S13. NMF analysis of scRNA-seq dataset. A**. Cell-level relationship between CD36 and IL1β expression in the scRNA-seq dataset. Each point represents single-cell expression level ($log_{10}$UMI) coloured by group (CD36 high, CD36 low, mixed as determined in B). Correlation assessed using Spearman's ρ (p-value indicated). Dashed lines indicate thresholds for top 10% expressing cells. **B**. Sample-level relationship between average CD36 and IL1β expression in the scRNA-seq dataset. Figure formatting as in A. **C.** Top-weighted genes defining the CD36-associated (in red) and IL1β-associated (in green) transcriptional programmes identified by NMF analysis. Gene weights reflect the individual gene contribution to the respective programme. **D**. Marker loading across NMF programmes. Bar graph showing the contribution of CD36 and IL1β to each inferred programme (as in Fig. 6F). **E.** Rank position of CD36 and IL1β within each programme based on gene weights (rank 1 = highest). The respective gene markers are among the top-ranked genes only within associated programmes. Colour-coding as in D. **F.** Association between CD36 and IL1β expression across programmes. Shown is the Spearman's correlation between CD36 and IL1β expression and programme usage across cells. Colour-coding as in D. **G**. Relationship between programme usage and CD36 (left) and IL1β (right) expression across all groups (CD36 high, CD36 low, mixed) assessed using Spearman's ρ (p-value indicated). **H**. Jaccard index comparing the top 20 genes defining CD36- and IL1β-associated programmes across different number of NMF components (k = 4-8). **I.** Spearman's correlation of CD36 and IL1β per cell programme usage across different number of NMF components (k = 4-8). **J.** Cell-level relationship between CD36 and IL1β programme usage. Panel formatting as in A. **K.** Sample-level relationship between CD36 and IL1β programme usage. Panel formatting as in B. **L.** Density-dependent regulation of resting human iPSC-derived macrophages from [1]. Shown is the mean transcript per million (TPM) and standard deviation of three replicates for the CD36- (top) and IL1β-associated programme genes (bottom) across three cell densities (Low, Mid, High) corresponding to 1.5, 3, and $6x10^4$ cells per well in a 96-well plate (0.44, 0.88, and $1.76x10^5$ cells/$cm^2$ respectively).

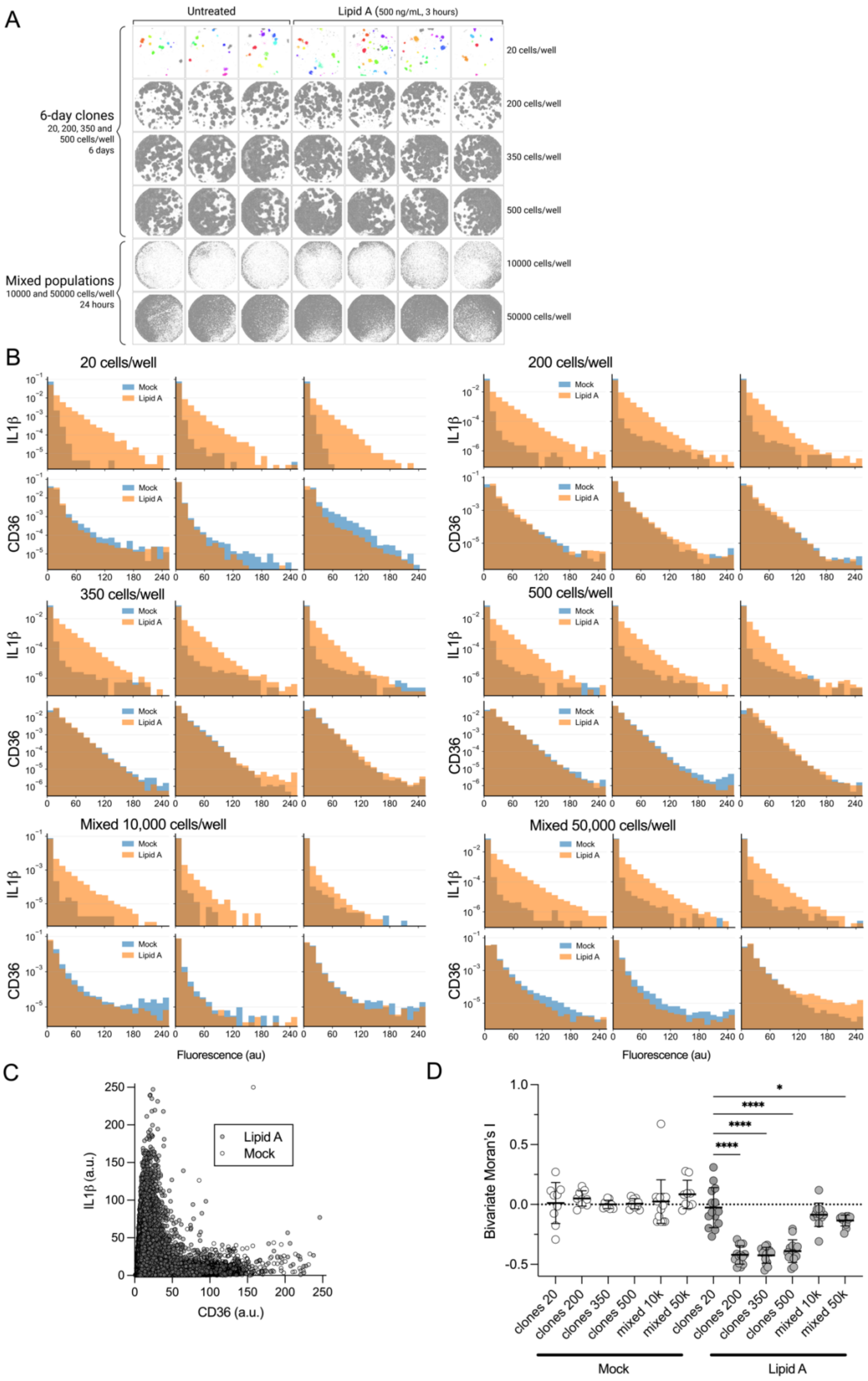

A
Untreated
Lipid A (500 ng/mL, 3 hours)
6-day clones
20, 200, 350 and 500 cells/well
6 days
Mixed populations
10000 and 50000 cells/well
24 hours
20 cells/well
200 cells/well
350 cells/well
500 cells/well
10000 cells/well
50000 cells/well
B
20 cells/well
200 cells/well
350 cells/well
500 cells/well
Mixed 10,000 cells/well
Mixed 50,000 cells/well
IL1β
CD36
Mock
Lipid A
Fluorescence (au)
C
Lipid A
Mock
IL1β (a.u.)
CD36 (a.u.)
D
Bivariate Moran's I
clones 20
clones 200
clones 350
clones 500
mixed 10k
mixed 50k
Mock
Lipid A

**Figure S14. CD36 and IL1β protein expression in low- and high-density clonal and mixed populations. A.** 96-well plate layout of the immunostaining experiment, representative of three biological replicates. Segmented individual clonal populations are highlighted in distinct colours; single non-clonal cells are shown in grey. **B**. Distributions of perinuclear protein expression in populations from A across three biological replicates. Cells stimulated with 500 ng/ml lipid A for 3 h (orange) and untreated cells (blue, Mock) are shown. **C.** Cell-level relationship between CD36 and IL1β protein expression. Shown are single cell fluorescent levels across clonal populations (seeded at 20 cells per well) either stimulated with lipid A (grey circles) or untreated (open circles). **D**. Spatial relationship between CD36 and IL1β expression. Shown is the global spatial-autocorrelation (bivariate Moran's I) between perinuclear CD36 and IL1β signals, in untreated (white) or lipid A-stimulated wells (grey). Individual wells are shown as circles, with mean ± SD across three biological replicates. Statistical significance assessed using ordinary one-way Anova with Dunnet's multiple comparison test against low density clones. Only statistically significant comparisons are shown (*$p < 0.05$, ****$p < 0.0001$).

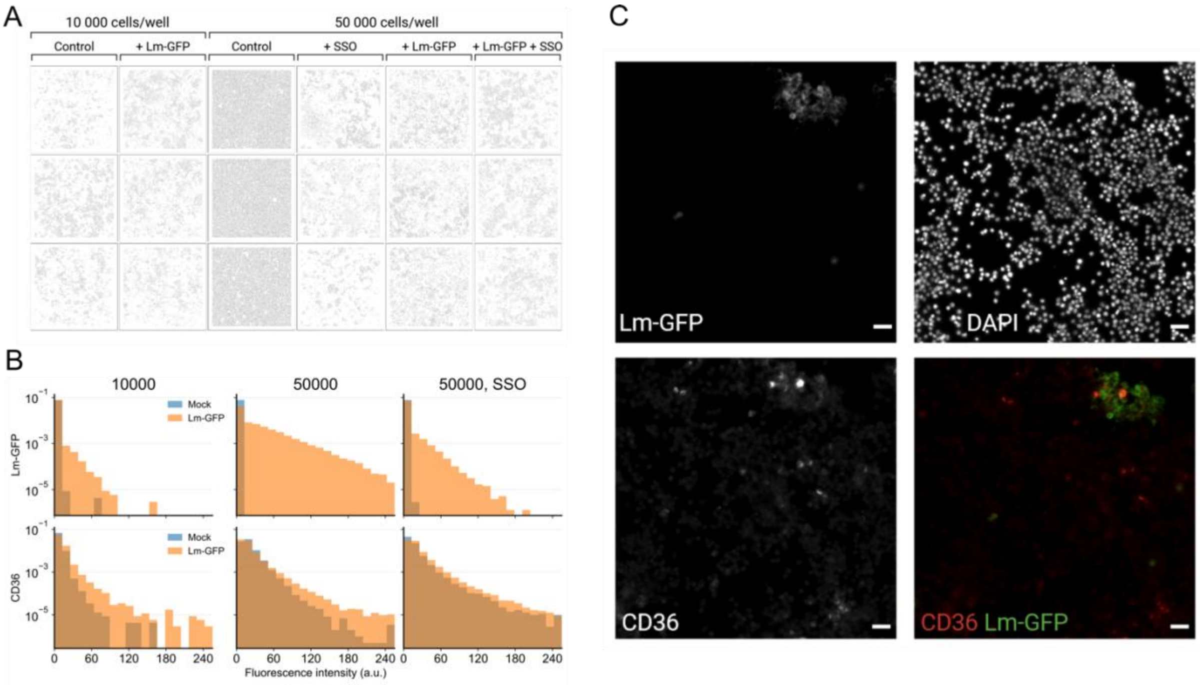

**Figure S15. Analysis of *Lm*-GFP and CD36 protein expression. A.** 96-well plate layout of the immunostaining experiment, representative of three biological replicates. Segmented single cells are shown in grey. Mixed iBMDM populations were seeded at two densities (10,000 or 50,000 cells per well), left uninfected (control) or infected with *Lm*-GFP at MOI = 5 for 45 min, and stained for CD36 at 24 h post-infection. High-density cultures were infected in the absence or presence of 200 µM SSO. **B**. Distributions of perinuclear *Lm*-GFP and CD36 immunostaining signals for the populations in A. Mock and *Lm*-GFP infection conditions are highlighted in blue and orange, respectively. Data pooled from three independent experiments. **C**. Individual microscopy channels (DAPI, CD36, and *Lm*-GFP) together with the corresponding composite image for *Lm*-GFP-infected cells pre-treated with SSO (shown in Fig. 7A). Scale bar: 50 µm.

#### 2. Supplementary Tables

| **TNFα and IL1β protein expression in clonal and mixed populations - 20x objective** | **RelA, RelB and CD36 protein expression in clonal and mixed populations - 20x objective** |
|---|---|
| **Nuclei detection**<br><br>channel=blue<br>staining_nuclear=1<br>normalization=0<br>denoising=1<br>denoising_strength=3<br>smoothing=1<br>smoothing_block_size=2<br>thresholding=1<br>thresholding_block_size=41<br>thresholding_baseline_offset=-1<br>opening=1<br>opening_repeats=1<br>opening_erosion_base_size=1<br>opening_dilation_base_size=2<br>closing=1<br>closing_repeats=1<br>closing_erosion_base_size=3<br>closing_dilation_base_size=3<br>uniformization=0<br>uniformization_brightness_threshold=25<br>contour_detection_threshold_low=80<br>contour_detection_thresholds_ratio=4<br>contour_detection_tolerance=1<br>contour_detection_reconstruct=0<br>min_solidity=0.96<br>min_area_as_median_fraction=0.3<br>max_area_as_median_fraction=1.1<br>splitting_adjacent=1<br>min_pocket_area=2<br>nuccut_channel_main=blue<br>nuccut_channel_aux1=(none)<br>nuccut_channel_aux2=(none)<br>nuccut_channel_main_weight=100<br>nuccut_channel_aux1_weight=50<br>nuccut_channel_aux2_weight=20<br>nuccut_color_similarity=5<br>nuccut_edge_impassability=5<br>nuccut_exclude_toggled=0<br><br>**Perinuclei derivatization**<br><br>inner_offset=0<br>width=2<br>neighbors_avoidance=0<br>overlap_avoidance=1<br>min_part_area=15<br>background_avoidance=1<br>background_avoidance_channel=green<br>background_avoidance_expansion=0<br>background_avoidance_smoothing=2 | **Nuclei detection**<br><br>channel=blue<br>staining_nuclear=1<br>normalization=0<br>denoising=1<br>denoising_strength=3<br>smoothing=1<br>smoothing_block_size=2<br>thresholding=1<br>thresholding_block_size=41<br>thresholding_baseline_offset=-1<br>opening=1<br>opening_repeats=1<br>opening_erosion_base_size=2<br>opening_dilation_base_size=1<br>closing=1<br>closing_repeats=1<br>closing_erosion_base_size=3<br>closing_dilation_base_size=3<br>uniformization=0<br>uniformization_brightness_threshold=25<br>contour_detection_threshold_low=80<br>contour_detection_thresholds_ratio=4<br>contour_detection_tolerance=1<br>contour_detection_reconstruct=0<br>min_solidity=0.96<br>min_area_as_median_fraction=0.3<br>max_area_as_median_fraction=1.1<br>splitting_adjacent=1<br>min_pocket_area=5<br>nuccut_channel_main=blue<br>nuccut_channel_aux1=(none)<br>nuccut_channel_aux2=(none)<br>nuccut_channel_main_weight=100<br>nuccut_channel_aux1_weight=50<br>nuccut_channel_aux2_weight=20<br>nuccut_color_similarity=5<br>nuccut_edge_impassability=5<br>nuccut_exclude_toggled=0<br><br>**Perinuclei derivatization**<br><br>inner_offset=0<br>width=2<br>neighbors_avoidance=0<br>overlap_avoidance=1<br>min_part_area=15<br>background_avoidance=0<br>background_avoidance_channel=magenta<br>background_avoidance_expansion=0<br>background_avoidance_smoothing=2 |

| **TNFR2 and F4/80 protein expression in clonal and mixed populations - 20x objective** | **IL1β and CD36 protein expression in high-density clonal and mixed populations - 20x objective** |
|---|---|
| **Nuclei detection**<br><br>channel=blue<br>staining_nuclear=1<br>normalization=1<br>denoising=1<br>denoising_strength=3<br>smoothing=1<br>smoothing_block_size=2<br>thresholding=1<br>thresholding_block_size=41<br>thresholding_baseline_offset=-1<br>opening=1<br>opening_repeats=1<br>opening_erosion_base_size=2<br>opening_dilation_base_size=1<br>closing=1<br>closing_repeats=1<br>closing_erosion_base_size=3<br>closing_dilation_base_size=3<br>uniformization=0<br>uniformization_brightness_threshold=25<br>contour_detection_threshold_low=80<br>contour_detection_thresholds_ratio=4<br>contour_detection_tolerance=1<br>contour_detection_reconstruct=0<br>min_solidity=0.96<br>min_area_as_median_fraction=0.3<br>max_area_as_median_fraction=1.1<br>splitting_adjacent=1<br>min_pocket_area=5<br>nuccut_channel_main=blue<br>nuccut_channel_aux1=(none)<br>nuccut_channel_aux2=(none)<br>nuccut_channel_main_weight=100<br>nuccut_channel_aux1_weight=50<br>nuccut_channel_aux2_weight=20<br>nuccut_color_similarity=5<br>nuccut_edge_impassability=5<br>nuccut_exclude_toggled=0<br><br>**Perinuclei derivatization**<br><br>inner_offset=0<br>width=2<br>neighbors_avoidance=0<br>overlap_avoidance=1<br>min_part_area=15<br>background_avoidance=0<br>background_avoidance_channel=magenta<br>background_avoidance_expansion=0<br>background_avoidance_smoothing=2 | **Nuclei detection**<br><br>channel=blue<br>staining_nuclear=1<br>normalization=0<br>denoising=1<br>denoising_strength=3<br>smoothing=1<br>smoothing_block_size=2<br>thresholding=1<br>thresholding_block_size=41<br>thresholding_baseline_offset=-1<br>opening=1<br>opening_repeats=1<br>opening_erosion_base_size=1<br>opening_dilation_base_size=2<br>closing=1<br>closing_repeats=1<br>closing_erosion_base_size=3<br>closing_dilation_base_size=3<br>uniformization=0<br>uniformization_brightness_threshold=25<br>contour_detection_threshold_low=80<br>contour_detection_thresholds_ratio=4<br>contour_detection_tolerance=1<br>contour_detection_reconstruct=0<br>min_solidity=0.96<br>min_area_as_median_fraction=0.3<br>max_area_as_median_fraction=1.1<br>splitting_adjacent=1<br>min_pocket_area=2<br>nuccut_channel_main=blue<br>nuccut_channel_aux1=(none)<br>nuccut_channel_aux2=(none)<br>nuccut_channel_main_weight=100<br>nuccut_channel_aux1_weight=50<br>nuccut_channel_aux2_weight=20<br>nuccut_color_similarity=5<br>nuccut_edge_impassability=5<br>nuccut_exclude_toggled=0<br><br>**Perinuclei derivatization**<br><br>inner_offset=0<br>width=2<br>neighbors_avoidance=0<br>overlap_avoidance=1<br>min_part_area=15<br>background_avoidance=1<br>background_avoidance_channel=green<br>background_avoidance_expansion=0<br>background_avoidance_smoothing=2 |
| **CD36 expression on *Listeria* infection - 40x objective** | |
| **Nuclei detection** | |

| |
|---|
| channel=blue<br>staining_nuclear=1<br>normalization=1<br>denoising=0<br>denoising_strength=10<br>smoothing=1<br>smoothing_block_size=5<br>thresholding=1<br>thresholding_block_size=121<br>thresholding_baseline_offset=0<br>opening=1<br>opening_repeats=1<br>opening_erosion_base_size=1<br>opening_dilation_base_size=1<br>closing=1<br>closing_repeats=1<br>closing_erosion_base_size=3<br>closing_dilation_base_size=3<br>uniformization=0<br>uniformization_brightness_threshold=25<br>contour_detection_threshold_low=80<br>contour_detection_thresholds_ratio=4<br>contour_detection_tolerance=1<br>contour_detection_reconstruct=0<br>min_solidity=0.95<br>min_area_as_median_fraction=0.4<br>max_area_as_median_fraction=1.2<br>splitting_adjacent=1<br>min_pocket_area=5<br>nuccut_channel_main=blue<br>nuccut_channel_aux1=(none)<br>nuccut_channel_aux2=(none)<br>nuccut_channel_main_weight=100<br>nuccut_channel_aux1_weight=50<br>nuccut_channel_aux2_weight=20<br>nuccut_color_similarity=5<br>nuccut_edge_impassability=5<br>nuccut_exclude_toggled=0<br><br>**Perinuclei derivatization**<br><br>inner_offset=2<br>width=5<br>neighbors_avoidance=2<br>overlap_avoidance=3<br>min_part_area=30<br>background_avoidance=0<br>background_avoidance_channel=green<br>background_avoidance_expansion=0<br>background_avoidance_smoothing=2 |

**Table S6.** Parameter sets used for nuclear detection and perinuclear derivatization in ShuttleTracker.

| | $N_0$ [cells] | N [cells] | Time [h] | $T_d$ [h] |
|---|---|---|---|---|
| 1 | 2.5x$10^5$ | 9.5 x$10^6$ | 72 | 13.72 |
| 2 | 5 x$10^5$ | 1.68 x$10^7$ | 72 | 14.21 |
| 3 | 5 x$10^5$ | 2.4 x$10^7$ | 72 | 12.89 |
| 4 | 5 x$10^5$ | 1.83 x$10^7$ | 72 | 13.87 |
| 5 | 2.5 x$10^5$ | 2.13 x$10^7$ | 96 | 14.98 |
| 6 | 5 x$10^5$ | 1.14 x$10^7$ | 72 | 15.96 |
| 7 | 2.5 x$10^5$ | 2.33 x$10^7$ | 96 | 14.68 |
| 8 | 5 x$10^5$ | 2.38 x$10^7$ | 72 | 12.93 |
| 9 | 5 x$10^5$ | 1.10 x$10^7$ | 72 | 16.15 |
| 10 | 2.5 x$10^5$ | 2.42 x$10^7$ | 96 | 14.55 |
| 11 | 5 x$10^5$ | 5 x$10^6$ | 48 | 14.45 |
| 12 | 2.5 x$10^5$ | 4.00 x$10^7$ | 96 | 13.11 |
| 13 | 2.5 x$10^5$ | 1.00 x$10^7$ | 72 | 13.53 |
| 14 | 5 x$10^5$ | 3.00 x$10^7$ | 72 | 12.19 |
| 15 | 5 x$10^5$ | 7.7 x$10^6$ | 72 | 18.25 |

**Table S7.** Estimation of the doubling time ($T_d$) for the iBMDM cells passaged under standard conditions ($N_0$- number of seeded cells, N- number of harvested cells, elapsed time in hours).

#### 3. Mathematical modelling of clonal expansion.

- **IL1β**

##### 1. Model 1 - Standard Two-State Model (Fig. 5G)

$IL1\beta$ expression was modelled using a two-state stochastic switching framework, in which cells transition between "ON" and "OFF" states at rates $k_{ON}$ and $k_{OFF}$, respectively (Fig. 5F, Model 1). Cells in both states proliferate at the same rate, $\lambda$. The following assumptions are made:.

1. The residence time in the ON state is independently and identically distributed (i.i.d), and follows an exponential distribution with a mean $\frac{1}{k_{OFF}}$.
2. The residence time in the OFF state is i.i.d. and follows an exponential distribution with a mean $\frac{1}{k_{ON}}$.
3. Cell cycle duration is i.i.d. and follows an exponential distribution with a mean $\frac{1}{\lambda}$.
4. Cell states are preserved upon division, such that both daughter cells inherit the state of their parent cell.

Following the framework established by Van Eyndhoven et al. (2025) [2], the switching rates $k_{ON}$ and $k_{OFF}$ were initially inferred from the odds ratio (*OR*) of daughter-cell pairs. At steady state, the mean fraction of cells in the ON state is given by:

$$f_{ON} = \frac{k_{ON}}{k_{ON}+k_{OFF}}.$$

*OR* quantifies the degree of association (memory) between daughter cells within a pair:

$$OR = \frac{4ab}{c^2} = \frac{(k_{ON}+\lambda)(k_{OFF}+\lambda)}{k_{ON}k_{OFF}},$$

where

- $a$ : probability that two selected cells are in ON State
- $b$ : probability that two selected cells are in OFF state
- $c$ : probability of one cell in ON State and the other cell in OFF State

*OR* together with the fraction of cells in the ON state were used as a system of equations to determine the switching rates $k_{ON}$ and $k_{OFF}$.

##### 2. Parameter Inference

For each set of daughter-cell pairs, *OR* and $f_{ON}$ are computed. *OR* = 1 indicates no memory (i.e., independent cell fates), while *OR > 1* indicates positive memory, with larger values corresponding to stronger association between daughter cells. The fraction of cells in the ON state was estimated as

$$\widehat{f_{ON}} = \frac{N_{ON}}{N},$$

where ($N_{ON}$) is the number of cells in the ON state and $N$ is the total number of cells. $OR$ was calculated from joint state probabilities of daughter-cell pairs, estimated from observed pair frequencies:

$$\hat{a} = \frac{N_{ON-ON}}{N_P},$$
$$\hat{b} = \frac{N_{OFF-OFF}}{N_P},$$
$$\hat{c} = \frac{N_{ON-OFF}}{N_P},$$

where $N_{ON-ON}$, $N_{OFF-OFF}$, and $N_{ON-OFF}$ denote the number of pairs in the corresponding states, and $N_P$ is the total number of daughter-cell pairs. The Odds Ratio ($\widehat{OR}$) is calculated as:

$$\widehat{OR} = \frac{4\hat{a}\hat{b}}{\hat{c}^2}.$$

The switching rates between the ON and OFF states were estimated by the following equations:

$$\widehat{f_{ON}} = \frac{k_{ON}}{k_{ON} + k_{OFF}},$$
$$\widehat{OR} = \frac{(k_{ON}+\lambda)(k_{OFF}+\lambda)}{k_{ON} * k_{OFF}}.$$

#### 3. Model fits

The cell proliferation rate, λ, was set to $\frac{\ln(2)}{14.4}\ h^{-1}$, according to the experimentally estimated value (Table S7). Bootstrapping was performed in two stages to estimate model parameters. In the first stage, resampling was performed 1,000 times to calculate $f_{ON}$ and $OR$ for each resample, along with 95% confidence intervals:

$$f_{ON} = 0.145\ (0.137, 0.152),$$
$$OR = 11.45\ (9.52, 12.97).$$

In the second stage, these bootstrapped estimates were used to calculate the switching rates $k_{ON}$ and $k_{OFF}$, using 1,000 bootstrapped datasets. The mean switching rates, and their corresponding 95% confidence intervals were then obtained from this second bootstrapping round:

$$k_{ON} = 0.00946\ h^{-1}\ (0.00935, 0.00957),$$
$$k_{OFF} = 0.05494\ h^{-1}\ (0.05444, 0.05561).$$

To validate the model against experimental observations, simulations were performed using a Stochastic Simulation Algorithm (SSA) with cell-doubling time of 14.4 h over 6 days. To enable a direct comparison between simulations and experimental observations, a nearest neighbor matching algorithm based on population size was applied. For each experimental colony, the specific simulated clone that minimized the absolute Euclidean distance in $\log_2 N$ space, thereby conditioning the phenotypic analysis of the fraction ON relative to the observed distribution of clone sizes, was identified.

- **CD36**

1. **Model 1 - Standard Two-State Model** (Fig. 5H, first panel)

First, the two-state stochastic switching model was applied to CD36 using *OR* and $f_{ON}$ to infer the constant switching rates $k_{ON}$ and $k_{OFF}$. Using the same inference framework as for IL1β, the following estimates (mean ± 95% confidence intervals) was obtained:

$$f_{ON} = 0.217\ (0.199, 0.235),$$
$$OR = 74.50\ (54.63, 105.80).$$

From those values, the corresponding switching rates were inferred:

$$k_{ON} = 0.00339\ h^{-1}\ (0.00338, 0.00342),$$
$$k_{OFF} = 0.01228\ h^{-1}\ (0.01222, 0.01236).$$

This model assumes a constant switching rate throughout the cell proliferation cycle, a simplification that fails to capture the population-dependent dynamics observed in CD36 expression, where the fraction of ON cells increases with clone size (Fig. 5B).

2. **Model 2 - Two-State Model switching dynamic $k_{ON}$** (Fig. 5H, second panel)

To better capture the dynamics observed in growing clonal populations, the model was extended by allowing the $k_{ON}$ rate to depend on population size ($N$). Consequently, time spent in the OFF state follows a non-stationary process, with the switching probabilities determined by the current population size. $k_{ON}(N)$ was defined a Hill functional form:

$$k_{ON}(N) = \mathrm{k}_{\mathrm{ON}}^{min} + \left(\mathrm{k}_{ON}^{max} - \mathrm{k}_{\mathrm{ON}}^{min}\right)\frac{N^n}{K^n+N^n},$$

where $\mathrm{k}_{\mathrm{ON}}^{min}$ and $\mathrm{k}_{ON}^{max}$ denote the minimal and maximal switching rates, respectively, *K* is the population size at which the switching rate reaches half-maximal activation, and $n$ is the Hill coefficient governing the steepness of the response.

The model was optimized using the following switching parameter set $k_{ON}^{min}, k_{ON}^{\max}, n, and\ K.$ $k_{OFF}$ was maintained as a constant throughout the simulations. Finally, λ, was set to $\frac{\ln(2)}{14.4}\ h^{-1}$, as before.

Model performance was assessed by comparing CV of the fraction ON cells, as a function of population size ($N$), between simulations and the experimental data, with additional validation based on *OR*. Both metrics were verified using a two-stage bootstrapping to ensure consistency within the 95% confidence intervals of the empirical results. Parameter estimation yielded the following values:

$$k_{ON}^{min} = 0.0016\ h^{-1}$$
$$k_{ON}^{max} = 0.0171\ h^{-1}$$
$$k_{OFF} = 0.0228\ h^{-1}$$

$$K = 84.45$$
$$n = 3$$

Although the model captured mean clonal CD36 expression, it failed to reproduce the observed inter-clonal variability,

3. **Model 3 - Two-State Model switching dynamic $k_{ON}$ with Beta distribution** (Fig. 5H, third panel, and 5I)

To capture the observed variability, the model was extended by allowing the maximal switching ON rate, $k_{ON}^{max}$, to vary across clones. Specifically, for each clone, $k_{ON}^{max}$ was modelled as a random variable drawn from a Beta distribution scaled by a factor γ. The population-dependent formulation of switching rate $k_{ON}(N)$ was retained as in the Model 2. Following previous methodology, parameter estimation yielded:

$$k_{ON}^{min} = 0.00067\ h^{-1},$$
$$K = 77.41,$$
$$\mathrm{n} = 8.65,$$
$$k_{ON}^{max} = Beta(\alpha = 0.38, \beta = 1.18),$$
$$\gamma = 0.152,$$
$$k_{OFF} = 0.028\ h^{-1}.$$

Model parameters were optimized using a weighted least squares objective function. The fitting process minimized the residuals between simulated and experimental values for both the mean CV of the 'Fraction ON' state. To ensure a direct comparison, simulated colonies were nearest neighbor matched to the experimental population by colony size, and errors were calculated across discrete $\log_2 N$ groups weighted by their bootstrapped confidence intervals.

Incorporating extrinsic noise in $k_{ON}^{max}$ substantially improved agreement with the experimental data, in terms of the observed variability in the fraction of ON cells across population sizes, while maintaining consistency with the measured odds ratio.

4. **Supplementary References**

1. Vaughan-Jackson A, Stodolak S, Ebrahimi KH, Johnson E, Reardon PK, Dupont M, et al. Density dependent regulation of inflammatory responses in macrophages. Front Immunol. 2022;13:895488. Epub 20221216. doi: 10.3389/fimmu.2022.895488. PubMed PMID: 36591218; PubMed Central PMCID: PMCPMC9800520.
2. Van Eyndhoven LC, Kinsler G, Zhang J, O'Farrell A, Abderrahim R, Srivastav A, et al. Lineage memory shapes viral resistance barriers in human skin. bioRxiv. 2025:2025.06.27.661951. doi: 10.1101/2025.06.27.661951.
